# Modeling colicin operons as genetic arms in plasmid-genome conflicts

**DOI:** 10.1101/2021.05.15.444200

**Authors:** Pavithra Anantharaman Sudhakari, Bhaskar Chandra Mohan Ramisetty

## Abstract

Plasmids are acellular propagating entities that depend, as molecular parasites, on bacteria for propagation. The conflict between the bacterial genome and the parasitic plasmids allows the emergence of ‘genetic arms’ such as Colicin (Col) operons. Endonuclease Col operons encode three proteins; an endonuclease colicin (cleaves nucleic acids), an immunity protein (inactivates its cognate colicin), and lysis protein (aids in colicin release via host cell lysis). Col operons are efficient plasmid-maintenance systems; (i) the plasmid cured cells are killed by the colicins; (ii) damaged cells lyse and release the colicins that eliminate the competitors; and (iii) the released plasmids invade new bacteria. Surprisingly, some bacterial genomes have Col operons. The eco-evolutionary drive and physiological relevance of genomic Col operons are unknown. We investigated plasmidic and genomic Col operons using sequence analyses from an eco-evolutionary perspective. We found 1,248 genomic and plasmidic colicins across 30 bacterial genera. Although 51% of the genomes harbor colicins, the majority of the genomic colicins lacked a functional lysis gene, suggesting the negative selection of lethal genes. The immunity gene of the Col operon protects the cured host, thereby eliminating the metabolic burden due to plasmid. We show mutual exclusivity of Col operons on genomes and plasmids. We propose an ‘anti-addiction’ hypothesis for genomic colicins. Using a stochastic agent-based model, we show that the genomic colicins confer an advantage to the host genome in terms of immunity to the toxin and elimination of plasmid burden. Col operons are ‘genetic arms’ that regulate the ecological interplay of bacterial genomes and plasmids.

## Introduction

The genetic conflict between propagating entities is never-ending genetic warfare that drives evolution and the emergence of ingenious ‘genetic arms’ [1]. Acellular propagating entities, such as plasmids, are stable outside the host but can propagate only upon parasitizing a bacterium. Plasmids encode strategic genetic arms, exemplified by Colicin (Col) operons, Restriction-modification systems (RMs), and Toxin-antitoxin systems (TAs), for stable maintenance in the host population [2]. Typically, an endonuclease Col operon comprises colicin activity gene (*cxa*), immunity gene (*cxi*), and lysis gene (*cxl*) [3, 4]. The *cxa* encodes colicin that cleaves nucleic acids; *cxi* encodes immunity protein that inactivates its cognate colicin, and *cxl* encodes lysis protein that aids in colicin released by lysing the host [5, 6]. Mutations in the lysis gene will hamper the colicin release [7], and hence the host survives. The immunity protein confers an advantage to its host as it inactivates both the endogenous and exogenous colicins (released by other colicin-producing bacteria), thereby protecting the host in its competitive niche [4].

Colicins (group A and B) are predominantly encoded on plasmids [8, 9]. Plasmidic colicin operons are implicated in plasmid maintenance [10], bacterial suicide [11], and other ecological phenomena [12-15]. However, the occurrence of colicin on bacterial genomes is intriguing. Firstly, carrying identical colicin on both genomes and plasmid is energetically expensive. Secondly, the lysis gene expression is counterproductive to the propagation of the genome; during colicin release, the cell is lysed, and then the genome is degraded.

The release of colicin determines the composition and stability of microbial populations [16, 17]. Simulation-based studies have elaborated on the factors such as time-point of colicin release, the concentration of colicin release, the initial abundance of the population, range of toxicity, stress conditions, resource availability, plasmid copy number, the concentration of lysis protein [12, 18, 19], etc., which determines the competition outcome.

This study explored the impact of genomic Col operons using sequence analyses and theoretical modeling from an eco-evolutionary perspective. Using nucleotide sequence homology and distribution patterns, we show that the occurrence of identical colicin is mutually exclusive on genomes and plasmids. Similar to toxin-antitoxin systems, we propose an ‘anti-addiction’ hypothesis for genomic colicins. Here, we provide a *proof-of-concept* that rationalizes our anti-addiction hypothesis for genomic colicins by simulating the competition between Col plasmids and genomes.

## Materials and methods

### Prevalence of Col operon

We analyzed nine nuclease colicins (E2, E3, E4, E6, E7, E8, E9, D, and cloacin DF13) (Supplementary file S1) in completely sequenced bacterial genomes and plasmids listed in the NCBI database as of July 2020. The above-mentioned colicin sequences were taken as reference sequences to perform a nucleotide homology search against all bacterial genomes and plasmids (Supplementary Sheet S1). We narrowed our search to specific colicins for which threshold values for sequence identity (∼>80%) and coverage (∼>90%) were set distinctively for each of the nine colicin types. The first hit list was created by filtering only “complete sequences”, “chromosome”, and “genomes” in MS-excel. The second hits list was created by performing individual TBLASTN for colicin, immunity, and lysis genes separately. Both the lists were compared to make a master list (Supplementary Sheet S2). Hits on genomes and plasmids were sorted separately and plotted as graphs. We then analyzed the colicin operons on genomes and plasmids for their conservation, prevalence, and multiplicity across genera. Genetic locus was examined for the conservation of operon (fig. S1). The distribution of colicins on both genomes and plasmids was plotted as graphs.

### Conservation of Col operon

Using the NCBI graphics of the hit sequences, genetic loci were checked for the completeness of the operon (Supplementary Sheet S3). The analysis was based on the sequence annotations of the respective strains submitted to NCBI. In the case of annotation artifacts for colicin activity gene, such as “HNH domain-containing gene” encoding <200 amino acids, were considered insignificant and were not included in this study. We analyzed the sequence similarity between the colicins for strains harboring colicins on both genome and their plasmid using the MAFFT alignment method in Jalview version 2.11.0 [20]. For visualization and presentation of the alignments, NCBI multiple sequence alignment viewer 1.16.2.

### Model description

We adopted a stochastic agent-based model to simulate the competition outcome between the strains: (i) sensitive to colicin (S); (ii) carrying colicin on the plasmid (C); and (iii) carrying colicin on the genome (C_g_). Within the C_g_ population, we show three variations: the population that (i) has complete operon (C_cil_); (ii) has colicin activity and immunity genes but not lysis gene (C_ci_); and (iii) has only immunity gene (C_i_). As a proof-of-concept, the framework of our model is adapted (with modifications) from a previously published dataset from *in vitro* competitive experiments with *E. coli* colicinogenic strains [12, 21]. Model parameters used for simulation were created following the experimental values. Initial communities were seeded in 100:5 (S:C) ratio in 250×250 lattice. Five different agents were used in this model: sensitive S cells, colicinogenic C, and C_g_ (C_cil_, C_ci_, and C_i_) cells. Each cell type is considered as an individual characterized by a particular status behavior. The model includes (i) reproduction of viable S, C, and C_g_ cells, (ii) lysis of C cells due to operon activation (**λ**_**c**_) and subsequent release of colicins and plasmids, (iii) death of C cells due to plasmid loss [22], (iv) switching of C to C_g_ (C_cil_ or C_ci_ or C_i_ at equal probability) cells (**λ**_**g**_), (vi) lysis of C_cil_ and C_ci_ cells with a release of colicin. We consider a small low-copy number (20 plasmids) plasmid containing endonuclease colicin operon.

### Simulation of the competition

Python 3.7 scripts were written to simulate the model. Libraries such as Matplotlib, pandas, and NumPy were used. The competition is modeled using a 2D-lattice, representing the width and height of the bacterial niche. Each grid in the lattice can contain one bacterial cell. We adapted an agent-based stochastic model to track the fate of each cell type (agents) as a function of space and time. The agents reproduce at specific growth rates. The multiplication of a cell is modeled using a Moore neighborhood (8 nearest neighbors) method. Each cell type is allowed to grow logistically yet limited by the carrying capacity and the total population at each instant. The logistic growth equation is calculated as,

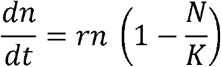

*r* is the growth rate, *n* is the total of each cell type, *N* is the total population size, and *K* is the carrying capacity. We performed a synchronous simulation where at each time step, the reaction conditions were applied for the corresponding population and not for the individual bacterial cell. The C, C_cil_, and rarely C_ci_ cells release toxin followed by their lysis. Once lysis occurred, the lattice site is now toxic to the nearby S cells. Our simulation governs the toxic interactions between the S cells and the toxic lattice sites by assuming exponential toxicity [19]. For exponential toxicity, the population of S cells within the radius of 5 is considered (exponential factor is 1/2^radius^), reducing the death probability exponentially with the increasing radius. The exponential toxicity is calculated as,

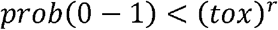

The state conversion probabilities are calculated binomially (*binomial* (*n,p,N*)),representing the binomial distribution with probability p and local trials n repeated N times). Model parameters, simulation conditions, and reaction rates are provided in the supplementary table with appropriate references (fig. 3b). The lack of experimental data is compensated by testing the competition outcome for a range of reaction rates. Uncertainty in the model is introduced through binomial probabilities for every parameter. The simulations with parameter sweeping were run 10 times for 15,000 minutes. The competition winner is assigned if they constitute more than 90% of the population in more than 5 out of 10 iterations. If none of the cell types attain 90% of the population, then the population is considered ‘disproportionately draw’. Our model predicts the relative success of plasmids in competitive environments. The success of plasmids and bacterial genomes is measured as the percentage of Col plasmid-containing cells (C) and genomic colicin-containing cells (C_g_) respectively in the population as a function of time.

## Results

### Distribution of colicin operons across bacterial genera

To examine the prevalence of genomic and plasmidic Col operons, we performed a nucleotide homology search for nine nuclease colicins against all bacterial genomes and plasmids. Within the limitations of available sequences, we obtained 1,248 genomes and plasmids hits across 30 bacterial genera harboring partial to complete Col operons (fig. 1a). All 30 genera are gram-negative *Gammaproteobacteria* and *Betaproteobacteria*. Plasmidic colicins were prevalent among the strains of *Klebsiella* (61%) and *Escherichia* (26%). Genomic colicins were prevalent among the strains of *Pseudomonas* (30%), *Salmonella* (28%), and *Yersinia* (15%) (table S1). The strains of *Klebsiella, Citrobacter, Salmonella*, and *Enterobacter* harbored Col operons on both genome and plasmid. By setting up the threshold values for sequence identity and coverage, we narrowed our analysis to nine type-specific colicins (E2, E3, E4, E6, E7, E8, E9, D, and cloacin DF13). Of the 240 strains encoding endonuclease colicins, cloacin DF13 is highly prevalent (78%) and distributed among the *Klebsiella* genus (fig. S3). We rarely found nuclease Col operon on the genome. Colicins E2, E8, and CloDF13 were found on genomes of *E. coli* and *Klebsiella*. Very few (3%) strains of *Shigella* harbored nuclease Col plasmids.

**Fig. 1.**
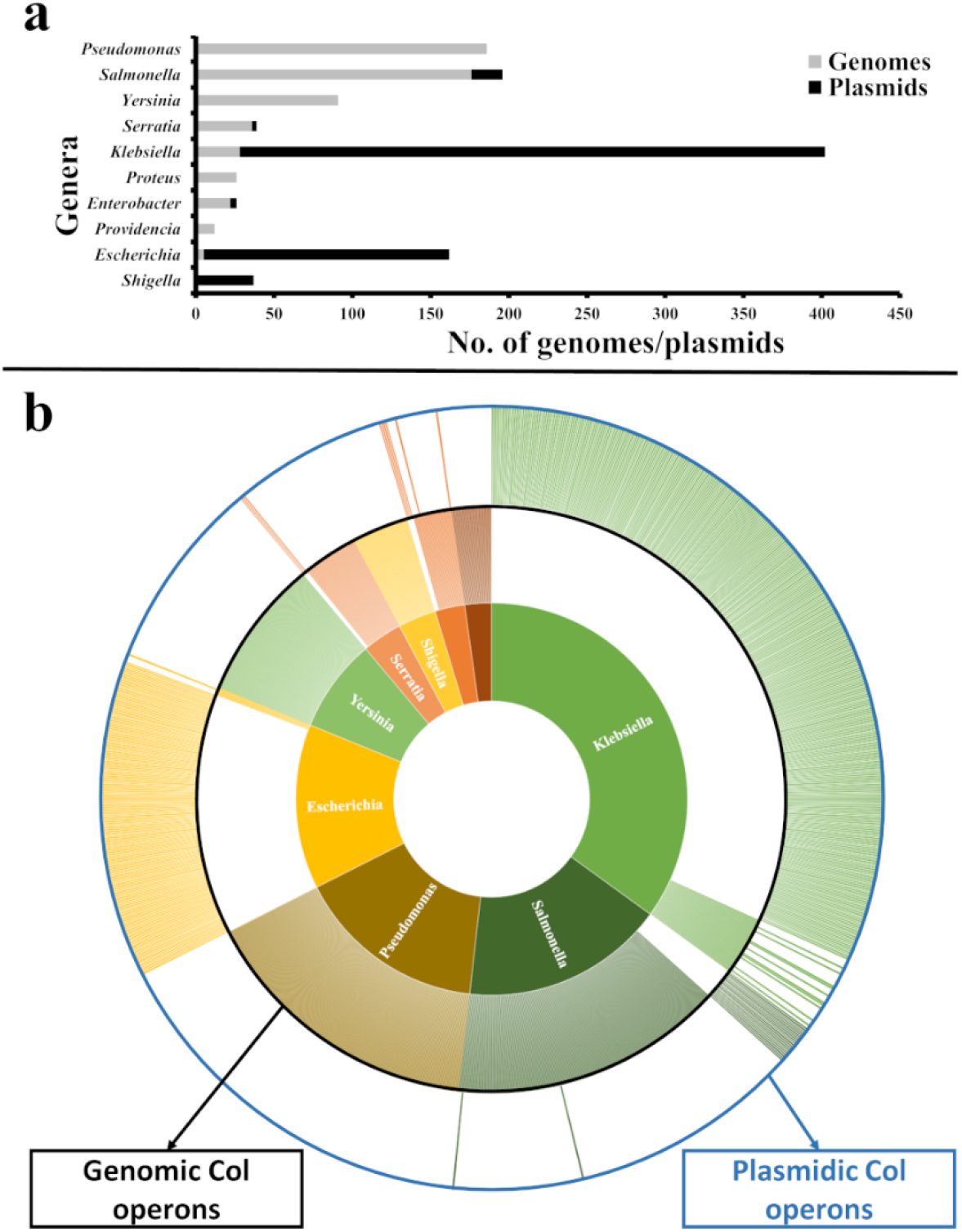
(a) Distribution of colicins across bacterial genera. We obtained 1248 genome and plasmid hits across 30 bacterial genera (belonging to families including *Enterobacteriaceae, Morganellaceae, Yersiniaceae, Pseudomonadaceae, Pectobacteriaceae, Vibrionaceae, Erwiniaceae*, and *Pasteurellaceae*, and *Burkholderiaceae*) (supplementary table 1). Here, we represent the distribution for genus comprising more than 10 colicinogenic strains. Plasmidic colicins were prevalent among the strains of *Klebsiella* (61%), *Escherichia* (26%), *Shigella* (6%), and *Salmonella* (3%). Genomic colicins were prevalent among the strains of *Pseudomonas* (30%), followed by *Salmonella* (28%), *Yersinia* (15%), *Serratia* (6%), and *Klebsiella* (4%). **(b) Mutual exclusivity of colicins on genomes and plasmids**. Hits list was sorted to distinguish whether strains carried colicins on the genome or plasmid. The data visualization is done using a sunburst graph. Different color code is given to each segment specific to genera (as labeled). For each genus, the outer segment represents the plasmids, and the inner segment represents the genomes.

### Mutual exclusivity of plasmidic and genomic colicin operons

Upon distribution pattern analyses, we found that some genera had Col operons only on genomes, and some had only on plasmids (fig. 1b). For example, we observed only genomic colicins in *Pseudomonas* strains while only plasmidic colicins in *Shigella*. Whereas in strains of *Klebsiella* (10 strains), *Escherichia* (1 strain), *Salmonella* (2 strains), and *Enterobacter* (2 strains), both genomic and plasmidic colicins were present (Supplementary Sheet S4). To rationalize the multiplicity of colicins within a strain, we examined the operon sequence similarity and conservation. We observed no strains carrying multiple copies of identical colicins (fig. S4). For example, *E. coli* BR10-DEC harbors colicins B and M on the genome, but colicin E1on its plasmid. *Citrobacter koseri* AR_0024 harbors incomplete colicin D on the genome but CloDF13-like operon on its plasmid. We also observed 85 strains harboring multiple Col plasmids but varied in the type of colicins they encode (Supplementary Sheet S5). For example, *E. coli* ACN001 harbors colicins K, E9-like, and E2-like operons on its plasmids pACN001-E, pACN001-D, and pACN001-B, respectively. *K. pneumoniae* NY9 harbors klebicin B and ColE3 on its plasmids pNY9_1 and pNY9_4, respectively. No bacterial strain had identical Col operons on genomes as well as plasmids.

### Degeneration of genomic Col operon

Colicin operon on genomes is highly degenerative. In our comparative gene conservation analysis, we found that most genomes either lacked lysis genes or harbored only immunity genes (fig. 2). To account for the prevalence of genomic colicins, as harboring lysis gene is strategically inefficient, we compared the conservation of genomic and plasmidic operons. Of the 620 genomic colicin operons, five were complete operons, 614 lacked lysis gene, and 323 lacked the colicin activity gene, whereas the immunity gene was conserved in almost all operons. For example, in *K. pneumoniae* TK421, the CloDF13 activity gene on the genome is replaced by a transposase gene, whereas its plasmid contains a complete CloDF13 operon. *Enterobacter roggenkampii* R11 harbors an S-type pyocin operon on its plasmid and an orphan immunity gene on its genome. Exceptionally, *K. pneumoniae* subsp. pneumoniae KPNIH24 harbors CloDF13 on its genome and plasmid. However, the integrity and functionality of these operons are unknown. The Col operons lacking functional lysis gene are either group B colicins that are released without host lysis or group A colicins with truncated lysis gene. We observed high conservation of the immunity gene on genomes (e.g., *Mixta theicola* SRCM103227) compared to colicin activity and lysis genes. Several truncations in the operon genes account for the evolutionary intermediates. Interestingly, we did not find any Col operon lacking immunity gene, as the absence of immunity protein is detrimental to the host.

**Fig. 2.**
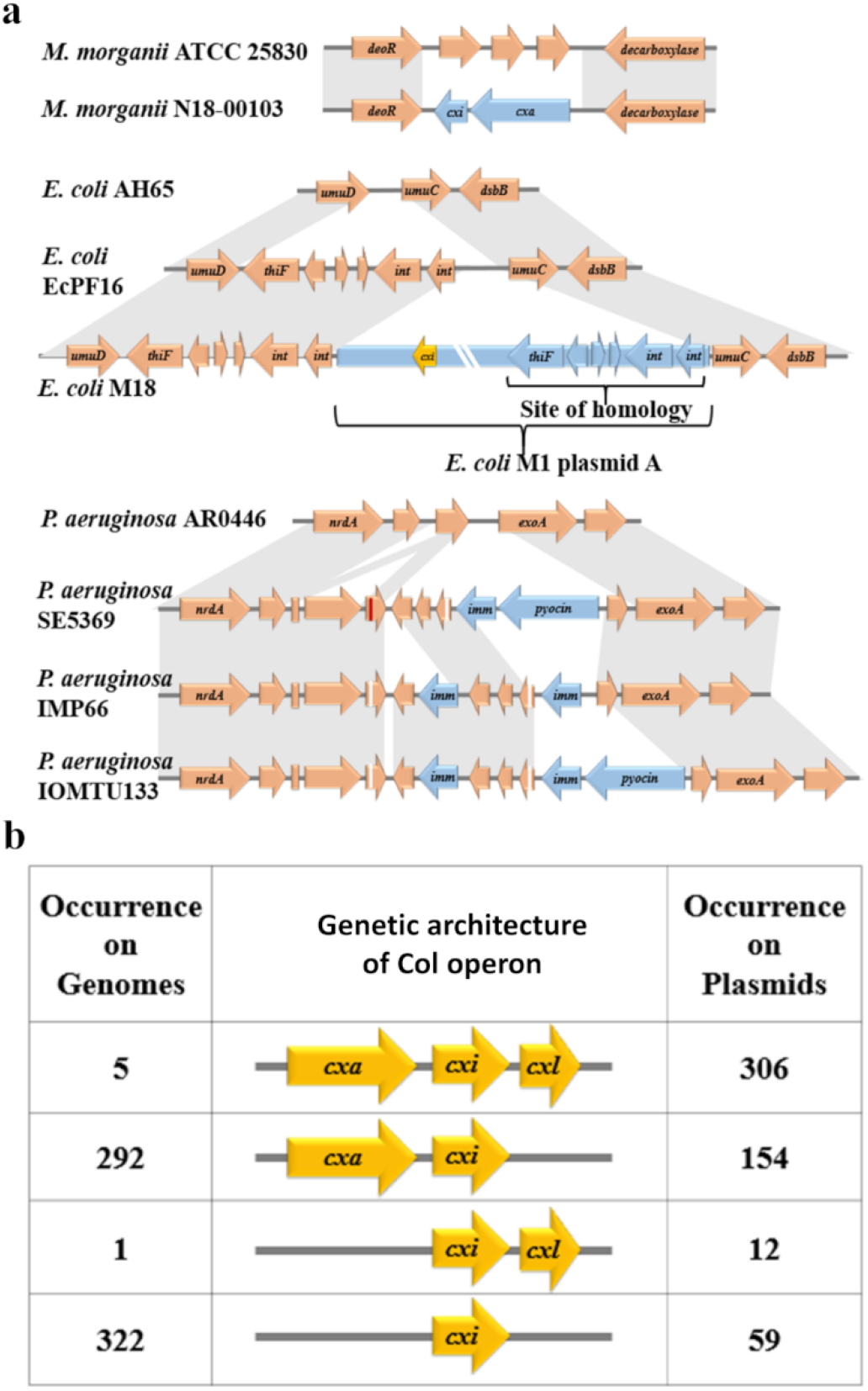
Genetic architecture of the operon. (a) Integration of Col operons on genomes. In *Morganella Morganii* N18-00103, the Col operon is flanked by *deoR* (deoxyribose operon repressor) and α-decarboxylase genes. We found *Morganella* strains lacking the operon (e.g., ATCC 25830) instead harbouring three hypothetical genes showing no sequence similarity with the Col operon. Likely that the operon integrated by replacing the region between the flanking genes. **(b)** In *Escherichia coli* M18, complete Col plasmid (*E. coli* M1 plasmid A) has integrated at homology site between the DNA polymerase V core genes (*umuD* and *umuC*). **(b) Degeneration of Col operon on genomes**. Here, we compare the different genetic organizations of the operons on genomes and plasmids. On the majority of the genomes, the immunity gene is orphaned. On the majority of the plasmids, the Col operon is complete, including the lysis gene. We observed 49% of plasmidic colicins and nearly 1% of genomic colicins are complete operons. Colicin operon on genomes is highly degenerative. For example, in *K. pneumoniae* TK421, the CloDF13 activity gene on the genome is replaced by a transposase gene. *Enterobacter roggenkampii* R11 harbors an orphan immunity gene on its genome. However, the integrity and functionality of the operon are unknown. Of the genomes carrying incomplete operon, 614 lacked a functional lysis gene. The Col operons lacking functional lysis gene are group B colicins or group A colicins with truncated lysis gene. We observed high conservation of the immunity gene on genomes (e.g., *Mixta theicola* SRCM103227) compared to colicin activity and lysis genes. Interestingly, we did not find any Col operon lacking immunity gene, as the absence of immunity protein is detrimental to the host.

### The competitive advantage of genomic colicins

Considering plasmids and genomes as competing entities, we developed a stochastic agent-based model [23, 24] to simulate the competition outcome between plasmids and genomes regulated by the Col operon. As a *proof-of-concept*, we adapted our model’s framework from a previously published dataset from *in vitro* competitive experiments with *E. coli* colicinogenic strains [12, 21]. Initially, we considered three cell types with the identical genetic background but differed in: (i) absence of Col operon (S-cell); (ii) carrying colicin on the plasmid (C-cell); and (iii) carrying colicin on the genome (C_g_-cells) (fig. 3a). Stochastic fluctuations in the parameters strongly affect the competition outcome [25]. We have incorporated such uncertainty into the model by testing a range of values for theoretical parameters (with binomial probabilities). Parameter sweeping tests the competition outcome for a range of parameter values abiding by Monte Carlo realizations, allowing statistical evaluation of the model (fig. 3b). Our model depicts the interactions between the colicinogenic and sensitive population and the competition outcome upon invasion by Col plasmids. In the theoretical model, the switching rate of S to C represents the plasmid invasion stage, the death rate of C due to plasmid loss represents the plasmid addiction stage, and the switching rate of C to C_g_ represents the integration stage. We did parameter sweeps to check the influence of invasion rate of Col plasmid (λ_s_), death of C due to plasmid loss rate, operon activation rate in C (λ_c_), acquisition rate of Col operon on the genome (λ_g_), and colicin toxicity (tox) on the competition outcome. We tested our model output for the otherwise possible range of values for each parameter to avoid biased observations and interpretations.

**Fig. 3.**
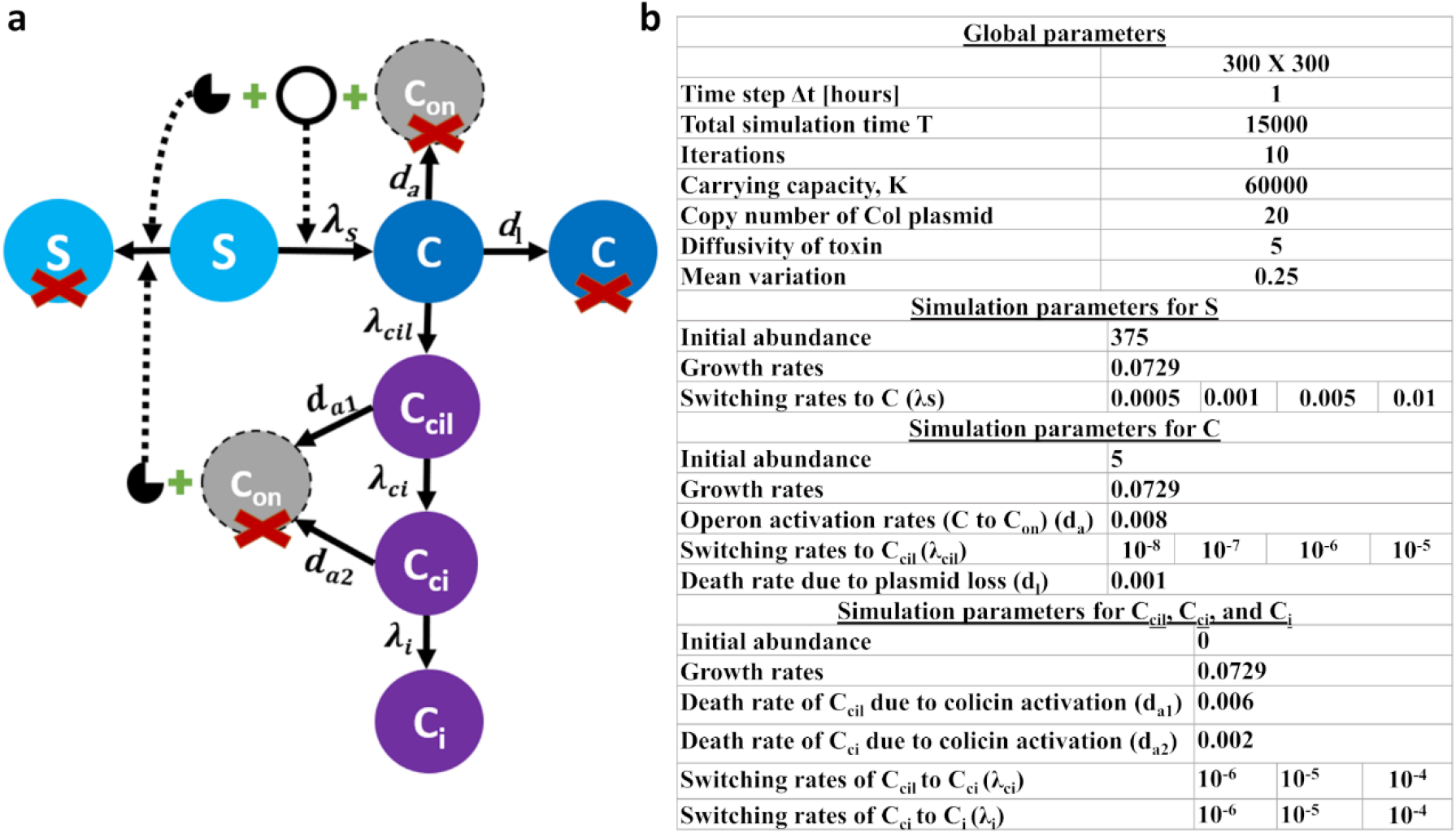
Theoretical model representing the interactions between the colicinogenic and sensitive strains. **(a)** Upon plasmid invasion, sensitive (S) cells transform into a colicinogenic (C) cell producing colicins. Once the Col operon integrates onto the genome (C_cil_ cells), though at a low frequency, the cells survive upon plasmid loss. However, with the accumulation of mutations in the operon, the degenerated operon retaining only the immunity genes is selected (C_i_ cells). λs-Switching rate of S to C; λ_cil_-Switching rate of C to C_cil_; λ_ci_-Switching rate of C_cil_ to C_ci_; λ_i_-Switching rate of C_ci_ to C_i_; d_a_-death rate of C due to operon activation; d_l_-death of C due to plasmid loss; d_a1_-death rate of C_cil_ due to operon activation; d_a2_-death rate of C_ci_ due to operon activation. **(b) Model parameters and reaction rates**. Parameter sweeps tested for each cell type are provided.

The reduction in the sensitive population represents the plasmid invasion where the plasmid is advantaged at the cost of the genome. The winning population of C represents the quasi-cooperation (Col plasmids benefit only the associated genomes by providing immunity to the toxin) by which both plasmids and genomes are benefitted (fig. 4, 6). Thus, genomic colicins are selected as a counter-strategy that eliminates the burden of Col plasmid imposed via addiction. In an evolutionary sense, the genomic colicins are no longer required after anti-addiction but for protection against the exogenous colicins. We then incorporated three genomic Col operon variations in the model: C_g_ population with (i) complete operon (C_cil_); (ii) colicin activity and immunity genes but not lysis gene (C_ci_); and (iii) only immunity gene (C_i_) (fig. 3a). Incoherence to *in-silico* analysis showing the high conservation of the immunity gene over colicin activity and lysis genes, our model predicts the selective conditions in which the C_i_ population outcompetes C_cil_ and C_ci_ (fig. 4).

**Fig. 4.**
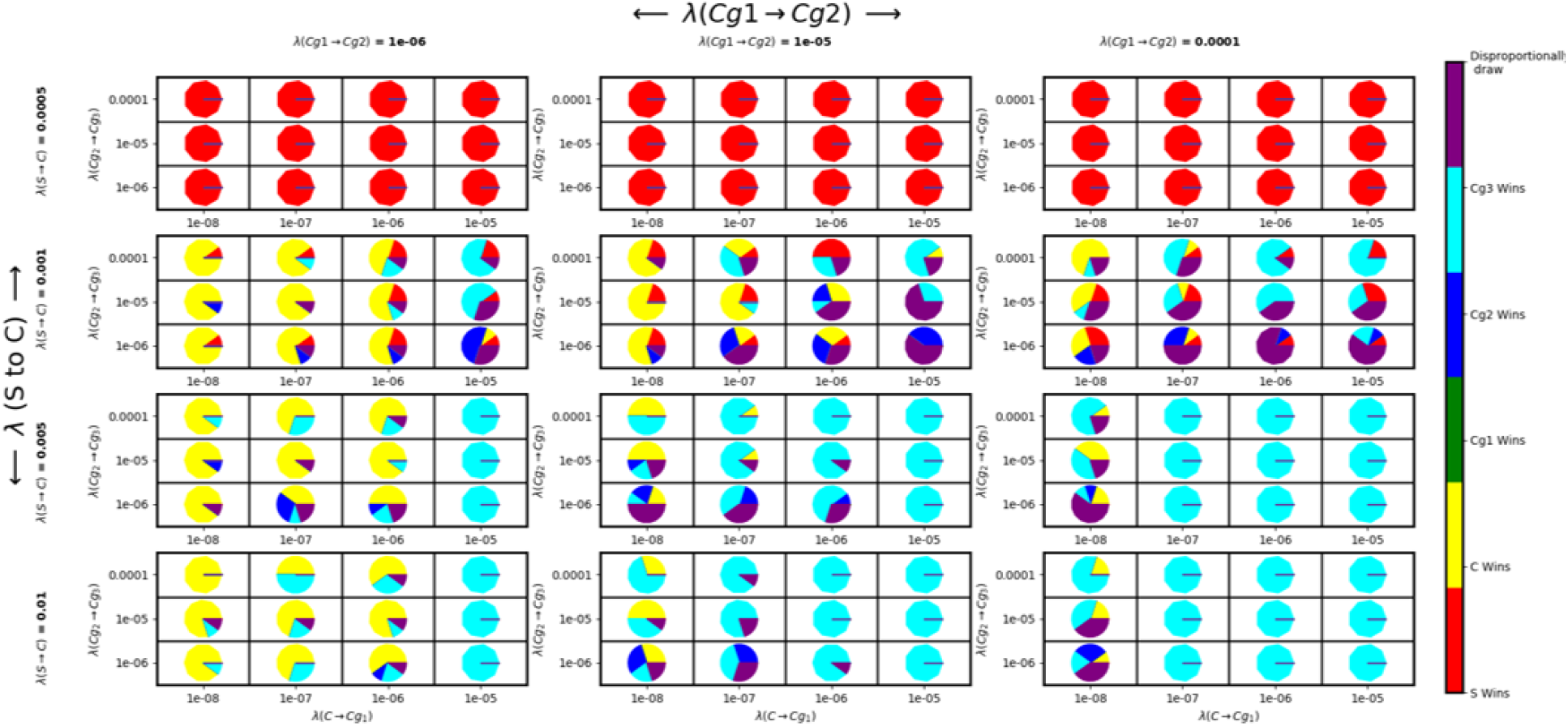
Simulation outcome of the competition between the cell types. Parameter sweeping for a range of values allows the uncertainty in the simulation outcome. Here, we tested the simulation outcome based on seven parameters, for each of which range of values was tested. Uncertainty in the model is introduced through binomial probabilities for every parameter. The simulations with parameter sweeping were run 10 times for 15,000 minutes. This combined pie chart shows the competition outcome for each of the tested parameter value. Colour code represents the respective cell type winning. A cell type is considered ‘winner’ if they constitute more than 90% of the population in more than 5 out of 10 iterations. If none reaches 90% proportion, then the population is considered as ‘disproportionately draw’.

**Fig. 5.**
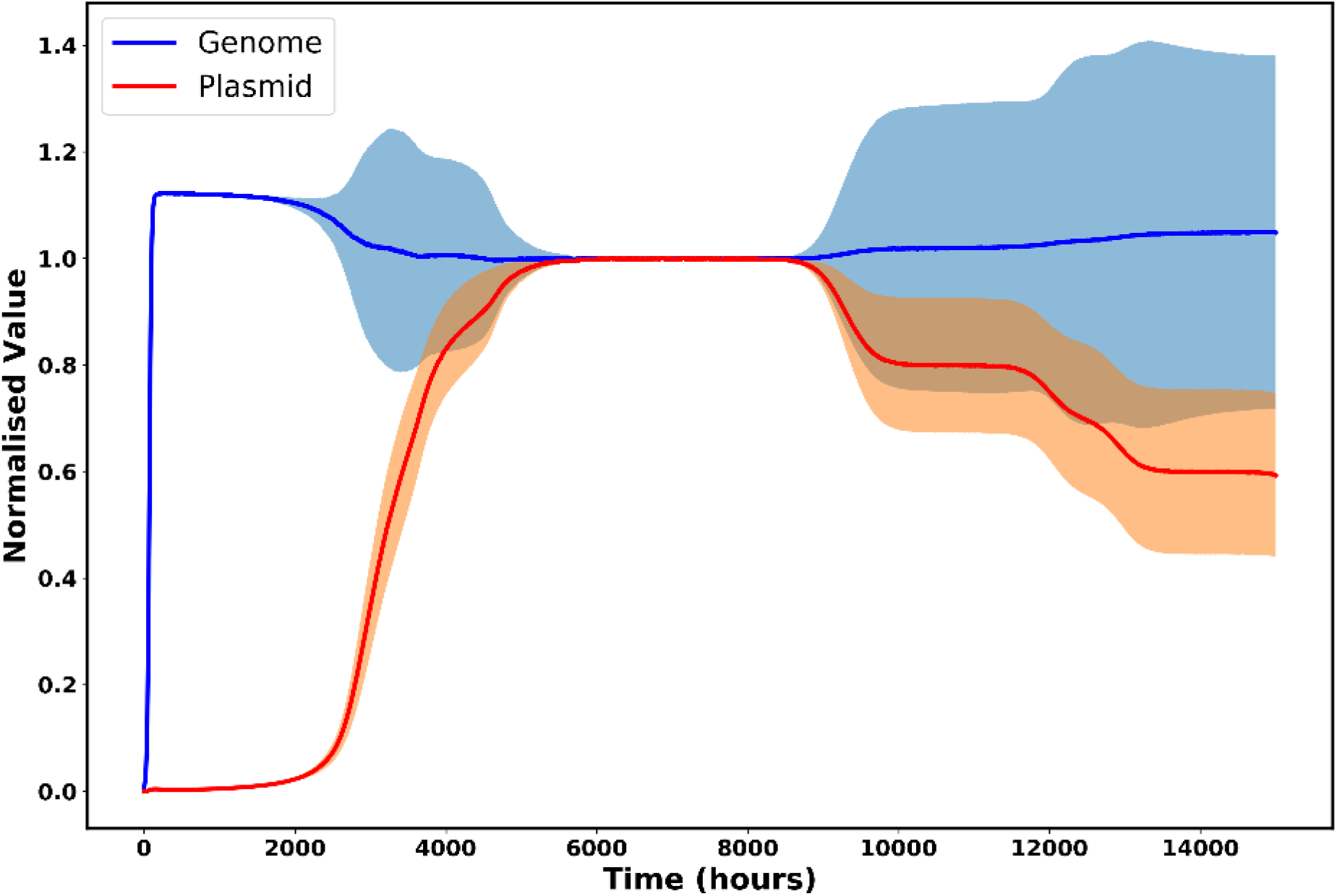
Modelling and simulating the implications of colicin in plasmids-genomes conflict. The dynamics of the number of plasmids and genomes over the period are plotted as a line chart. The total number of S, C, C_cil_, C_ci_, and C_i_ cells was taken to represent the total number of genomes and the number of C cells alone to represent the total number of plasmids. Simulations were iterated ten times, considering the parameter values that show uncertainty in the game outcome, and the statistical summary of the output is shown as a line. The additional lines above and below the mean line represent the variability (mean variation of 25%) in the data at each plotted point. Two subtle regions (between 2000-4000 hours and after 9000 hours) in the graph represent the uncertainty wherein genomes or plasmids get equal chances to outcompete the other. Parameters and values used: S to C conversion is 10^−3^, C death due to plasmid loss is 10^−4^, C to C_cil_ conversion is 10^−8^, C_cil_ to C_ci_ conversion is 10^−5^, C_ci_ to C_i_ conversion is 10^−5,^ and colicin toxicity is 50%.

**Fig. 6.**
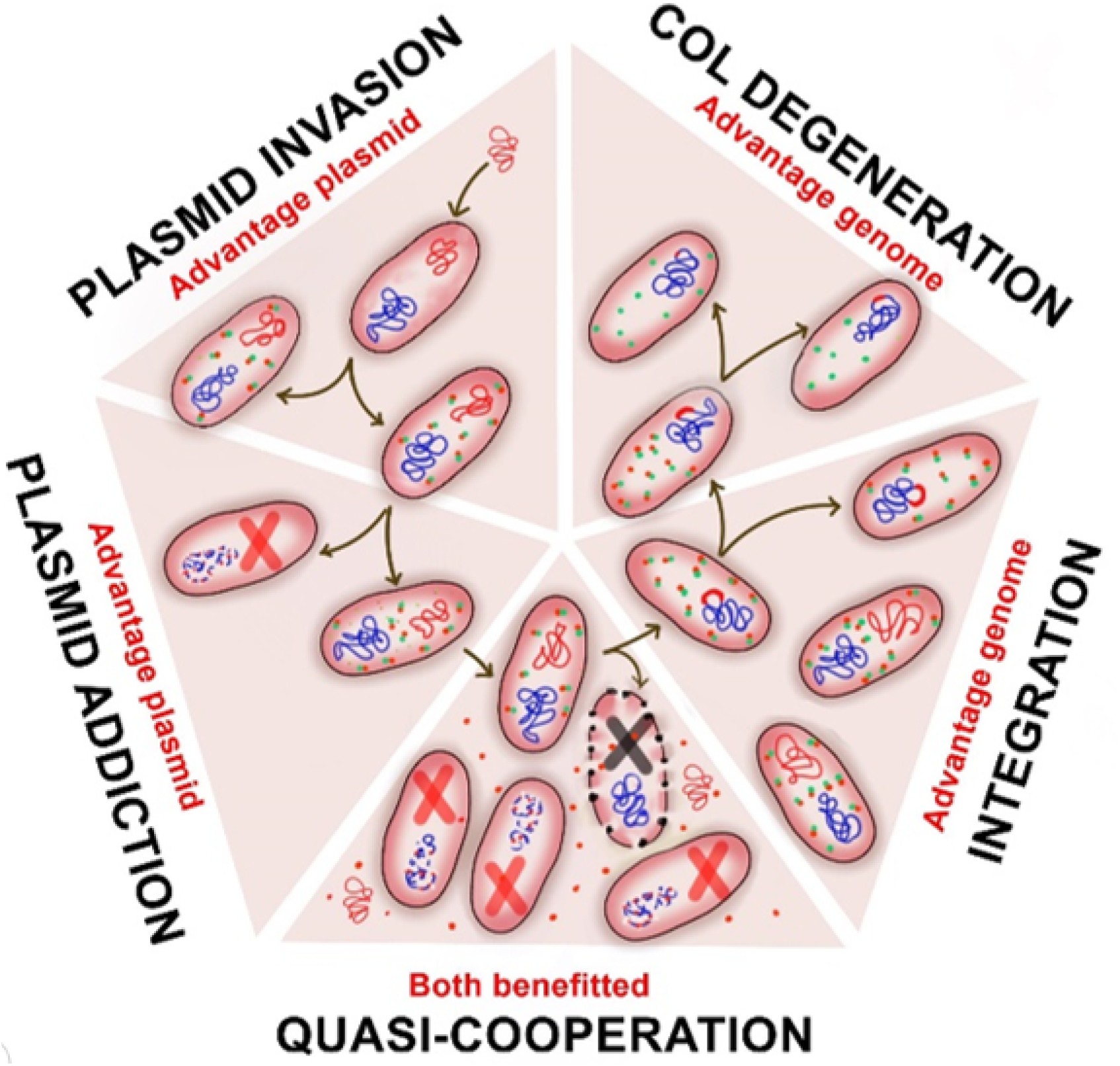
The eco-evolutionary cycle of colicin plasmids and host genomes. The ecological cycle can be divided into five theoretical stages. (i) Plasmid invasion. The plasmid is advantaged by its association with the host genome. (ii) Plasmid addiction. Cells cured of the plasmid are eliminated from the population. The plasmids are advantaged at the cost of host cells. (iii) Quasi-cooperation. The plasmid-containing cells are resistant to exogenous colicins due to the neutralizing effects of the immunity protein. They have increased genome propagation along with plasmid propagation. It is ‘quasi’ because the plasmid-free clones of the host cell are also killed. (iv) Chromosomal integration. The chromosomal colicin operons can act as anti-addiction systems by providing immunity protein to protect the host from exogenous colicins. The host genome is advantaged because the cured daughter cells are no longer killed by the plasmid-encoded colicins and reduce the metabolic burden of harboring the plasmid. (v) Colicin operon degeneration. Selection against the lethal gene (lysis gene) results in stabilizing the genomic colicins in the population.

## Discussion

This study investigated the abundance of Col operon on genomes and proposed an anti-addiction phenomenon for genomic colicins. We first analyzed the abundance of Col operons on plasmids as colicins are predominantly plasmidic [4]. Surprisingly, we found that genomic colicins are prevalently (51%) spread across the bacterial genera. However, most endonuclease colicins are plasmidic. Plasmidic Col operons impose ‘addiction’, a phenomenon where the cured daughter cell dies due to the toxic effects of the Col proteins. Despite the metabolic burden on the host, addiction ensures the maintenance of plasmid-bearing cells in the population. The best strategy to eliminate the metabolic burden of the plasmid is to carry Col operon on the genome. The vast number of genomic colicins indicate an eco-evolutionary advantage. In theory, all infected bacteria have to either retain the plasmid or die due to colicins. ‘Forced’ plasmid maintenance (addiction) is similar to being parasitized by phages like M13 [26]. A Col plasmid harboring bacterium cannot be cured of the plasmid because of addiction: a choice of maintaining the plasmid or death. The presence of Col operon on the genome will allow plasmid curing through ‘anti-addiction’, a phenomenon characterized by protection of the cured host by the genome encoded immunity protein. Based on the above rationales, we devised two premises. (i) A bacterium harboring Col plasmid would not have a Col operon on its genome, and (ii) a bacterium with genomic Col operon would not harbor a plasmid-encoding identical Col operon. Our observations fulfill both the premises supporting the anti-addiction hypothesis (fig. 1), corroborating the observations in the anti-addiction hypothesis of TAs [27]. The prevalence and mutual exclusivity of genomic colicins indicate a selective advantage of carrying colicin operons on genomes.

The comparative conservation analyses highlight the strategy of operon degeneration. We can interpret that the early loss of the immunity gene from the genome is detrimental to bacterial survival. In contrast, the early loss of lysis and colicin genes benefits the host strain [28]. In an evolutionary sense, the genomic colicins are no longer required after anti-addiction but for protection against the exogenous colicins. Of the three Col operon genes, only immunity protein is helpful in anti-addiction and protection against exogenous colicins. The colicin protein is harmful but can be antagonized by the immunity protein. The lysis protein is detrimental to the cell. Colicin and lysis genes have no evolutionary benefit to the cell. Therefore, the lysis and colicin genes should be less conserved relative to immunity genes.

Our theoretical analysis unravels the probable advantage of genomic colicins and their selection over plasmidic colicin. The competition outcome is influenced by a range of parameters such as plasmid invasion rate, degree of plasmid addiction, cost of lysis gene, chromosomal integration rate, and operon conservation rate. The preferential conservation of the immunity gene over Col and lysis genes allows the host to dispose of the burden imposed by Col plasmids and the cost incurred by *col* and lysis genes. Interpretation of the model in terms of plasmid-genome conflict gives a better picture of the eco-evolutionary cycle that governs the dynamicity in the population. In the conflict between plasmids and genomes, there exist dynamic time points wherein genomes or plasmids get equal chances to outcompete the other or exist in cooperation (fig. 6).

### The Plasmid-Genome conflict

The Plasmid-Genome conflict is exploited by the plasmid maintenance systems such as Col operons. The eco-evolutionary cycle of colicin plasmids and host genomes includes plasmid invasion, plasmid addiction, quasi-cooperation between the plasmids and genomes, plasmid loss via anti-addiction (fig. 6). During plasmid invasion, the plasmid is advantaged as it propagated along with its host genome. With the establishment of plasmid addiction, only plasmid-bearing cells are selected in the population against plasmid-free clones despite the metabolic burden incurred by the plasmid [29, 30]. A daughter cell that fails to inherit a copy of the plasmid becomes sensitive to the lethality of exogenous colicins as there is no source for the production of cognate immunity protein. The colicin operon acts as ‘genetic arms’ to induce the addiction to the plasmid. Through the elimination of cured cells, plasmids increase their maintenance within the population. Therefore, plasmids are advantaged at the cost of the host genome. But, in conditions of host DNA damage and the induction of SOS response, the plasmid-encoded colicins are the stress without which the bacteria would have grown better [31]. The lysis gene expression results in the host cell lysis and the release of colicin proteins and plasmids. The released plasmids have an opportunity to be taken up by other cells. The colicin proteins induce lethality in plasmid-free cells resulting in the elimination of competition. The plasmid-containing cells are resistant to exogenous colicins due to the neutralizing effects of the immunity protein. They have increased genome propagation along with plasmid propagation. The ultimate destiny of the Col plasmid-containing cell is to either remain parasitized by the plasmid, lyse due to lysis protein or die due to damage. Occasionally, the colicin operon may integrate into the chromosome through non-specific means, associated with transposons, or whole plasmid integration (fig. 2a, 5). In such cases, the host is relieved from plasmid addiction. The chromosomal colicin operons can act as anti-addiction systems by providing immunity protein to protect the host from exogenous colicins. Here, the host genome is advantaged because the cured daughter cells are resistant to the endogenous and exogenous colicins, and there is a reduction in the metabolic burden incurred by the plasmid. Interestingly, two of the three genes, the colicin gene, and lysis gene, are costly to maintain because of their respective lethal activities of DNA degradation and cell membrane lysis. The immunity gene-encoded protein is essential as long as the colicin gene is also encoded and immunity against the exogenous colicins. The lysis gene and the colicin genes are lost from the most chromosomal colicin operons. Therefore, the genome is advantaged.

## Conclusion

This study highlights the prevalence of colicins on the genomes and plasmids of bacterial genera. Mutual exclusivity of colicins on genomes and plasmids supports the ‘anti-addiction’ hypothesis [32]. Colicins are the ‘genetic arms’, which influence the ‘arms race’ between genomes and plasmids [13]. Plasmids occasionally provide an ecological advantage to the host; nevertheless, they are parasites and a metabolic burden. Colicin on plasmids ensures plasmid propagation via host addiction. This study highlights the prevalence of colicins on the genomes and plasmids of bacterial genera. The mutual exclusivity of colicins on genomes and plasmids supports the ‘anti-addiction’ hypothesis [32]. Genomic colicins enable anti-addiction to plasmids, thereby eliminating the metabolic burden of plasmids on the host. Empirical evidence is required to confirm these observations. Colicin operons play a significant role in the ecology and evolution of bacterial genomes and plasmids. The antagonistic role of colicins promotes genome diversity in bacteria [33]. Studying the phenomenon of anti-addiction may help mitigate the plasmid-mediated virulence and antibiotic resistance.

## Supporting information

Supplementary Sheet S1

## Acknowledgments

This work was supported by Prof T. R. Rajagopalan grants and Central Research Facility sanctioned by SASTRA Deemed University, Thanjavur.

## Code availability

The python Source code can be found at https://github.com/Pavi31/Plasmid-Genome_conflicts.git.

