## Supplementary material for "Modeling colicin operons as genetic arms in plasmid-genome conflicts": fig. S1

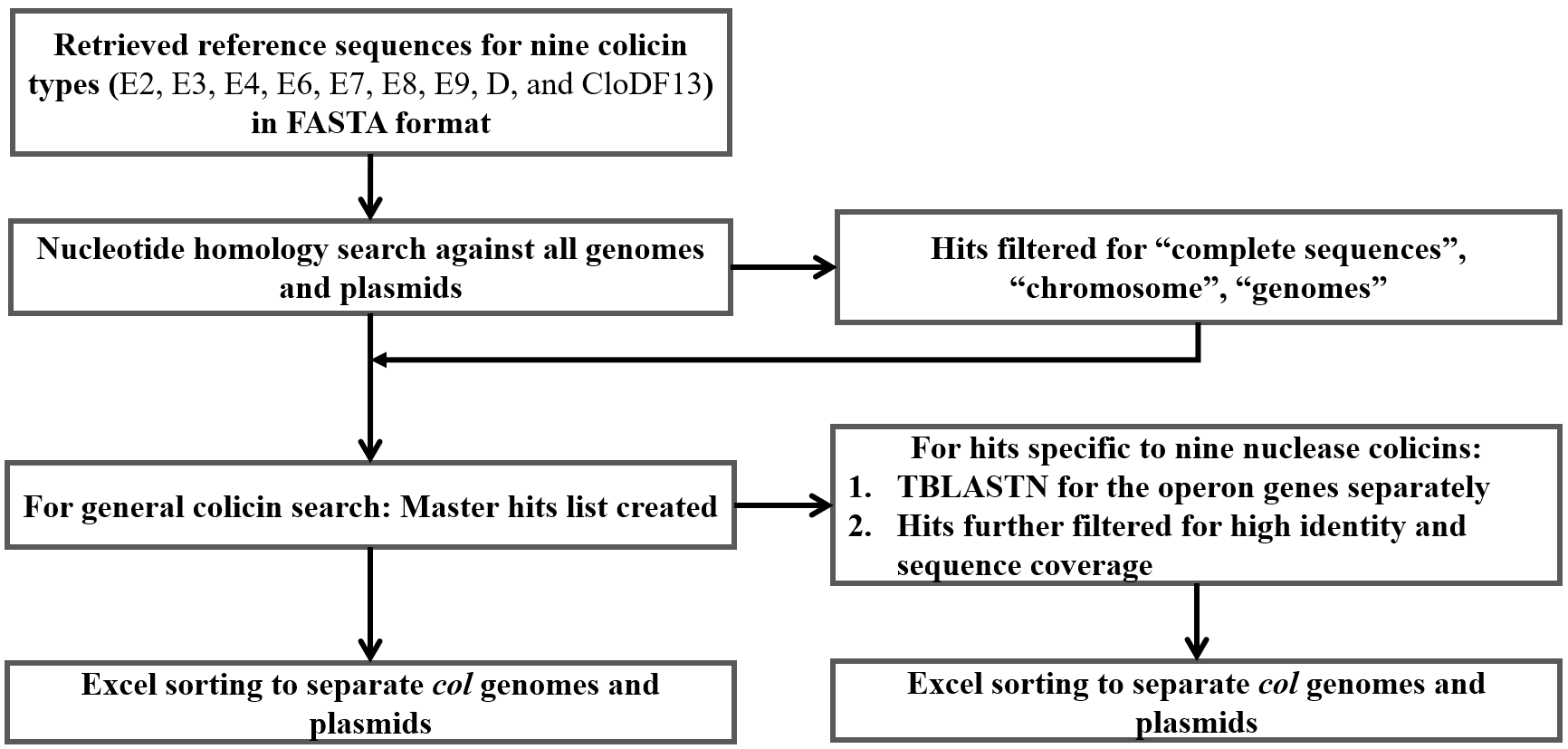

**Fig.** **S1** Flowchart briefing the *in-silico* methodology for investigating the distribution of colicins across bacterial genera.

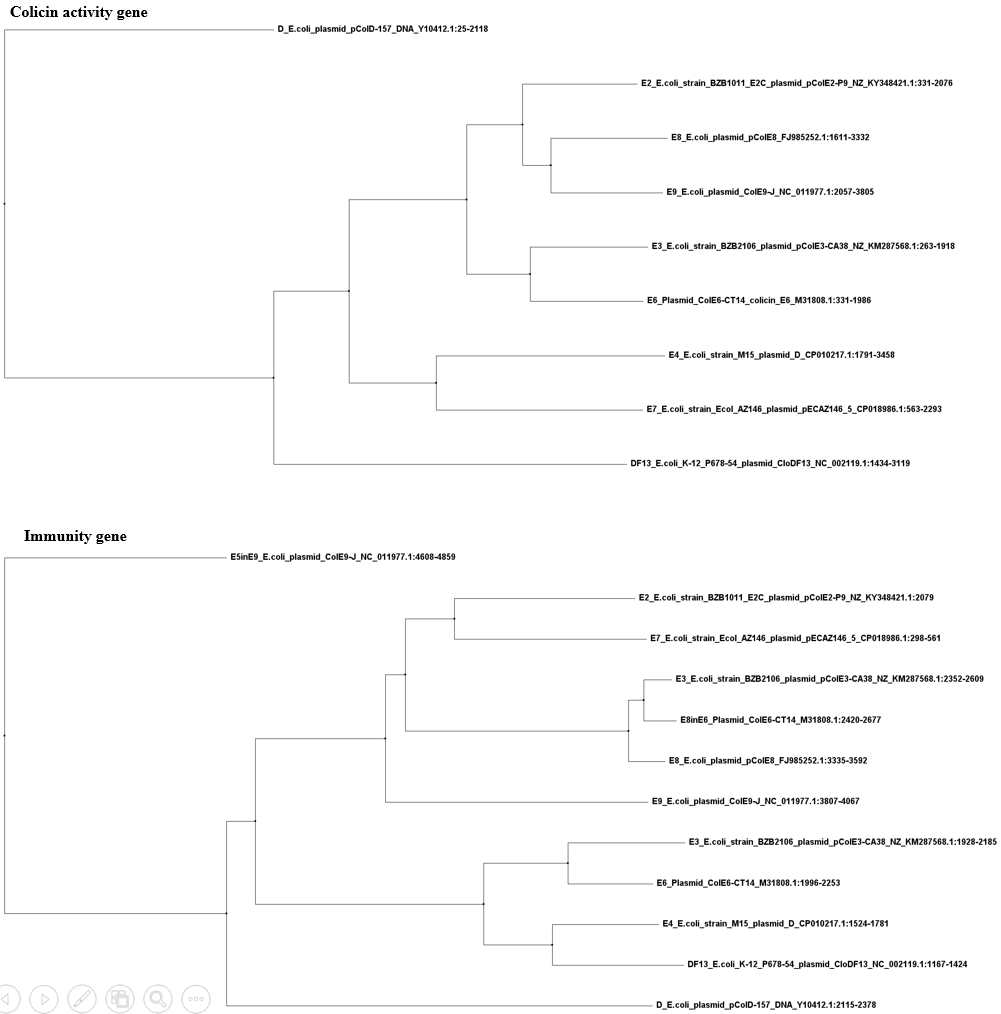

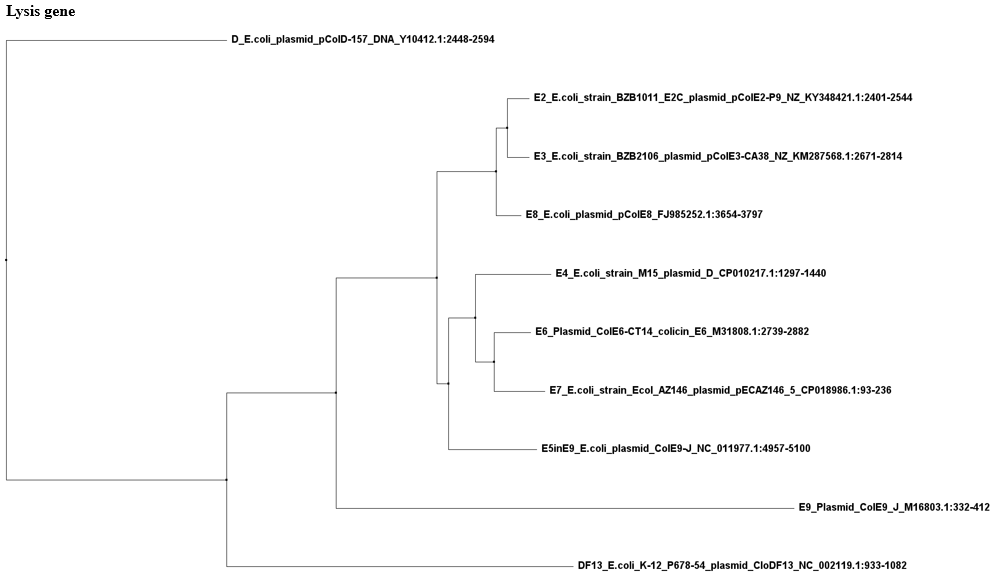

**Fig. S2 Diversification of colicin E-series.** We aligned the reference sequences of nine plasmid-encoding endonuclease colicin operons (E2, E3, E4, E6, E7, E8, E9, D, and CloDF13). The phylogenetic relationship for the colicin, immunity, and lysis genes were inferred using separate trees. The phylogeny includes the additional immunity and lysis genes encoded in E3, E6, and E9 operons. Similar branching pattern was observed for the activity and immunity genes of all colicins except E7, whose immunity gene (close to E2) showed distinct divergence from that of its activity gene (close to E4). DF13, which shares rRNase activity with E3 and E6, showed reduced similarity with E3 and E6 at the nucleotide level. Contrastingly, the activity genes of E4 with rRNase activity and E7 with DNase activity were more closely related than other functionally related colicins. ColD was more distantly related to colicin E-series. The lysis genes of nine colicins showed distinct branching patterns from that of their corresponding activity and immunity genes. But, lysis genes of endonuclease colicins share high sequence identity. Thus, lysis genes are more conserved across all the nine colicin types when compared to their activity and immunity genes.

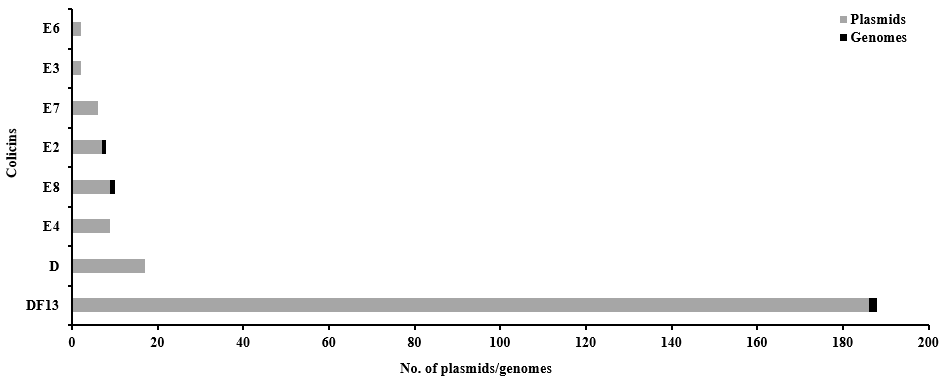

**Fig. S3** Prevalence and distribution of different endonuclease colicins (E2, E3, E4, E6, E7, E8, E9, D, and Clo DF13) on genomes and plasmids of different bacterial genera. Reference sequence of each colicin type was used as a query in standard blastn against all bacterial genera. The hits were filtered based on stringent threshold values of sequence identity and coverage. Hits were sorted based on their occurrence on genomes and plasmids. We obtained a total of 240 colicinogenic strains carrying (Supplementary data S1). The order of prevalence was CloDF13 followed by colicins D, E8, E4, E2, E7, E6, E3, and E9. CloDF13 was highly distributed on the plasmids of *Klebsiella*. However, other types were distributed among the strains of *Escherichia* and *Shigella*. We found that endonuclease colicins are plasmidic except for colicins E2, E8, and CloDF13.

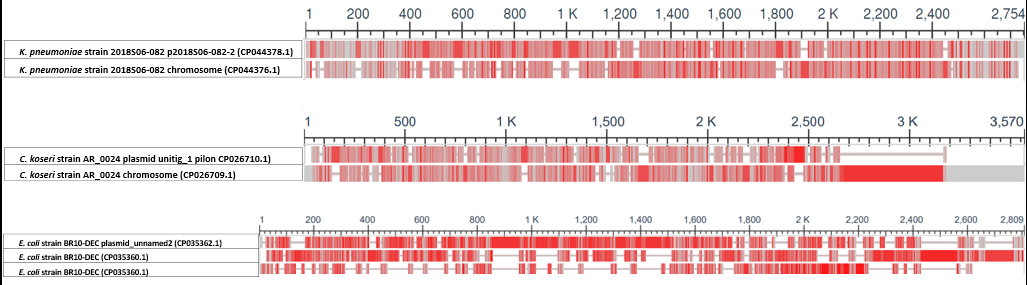

**Fig. S4** **Sequence dissimilarity between genomic and plasmidic colicins of a single strain.** For strains carrying colicins on both genome and plasmids, we analysed the colicins conservation and nucleotide sequence similarity. We found 18 strains harbouring colicins on both genome and plasmid (supplementary data S3). However, the colicins were not identical. Rather, the colicins differed significantly in terms of their nucleotide sequences. Sequences were aligned by MAFFT alignment method using python scripts. Alignment visualisation was done using NCBI MSA Viewer. Red markings indicate the sequence dissimilarity and grey markings indicate consensus sequences. *E. coli* BR10-DEC harbours colicins B and M on genome but colicin E1on its plasmid. *Citrobacter koseri* AR_0024 harbours incomplete colicin D on the genome but CloDF13-like operon on its plasmid. We also observed strains harbouring multiple Col plasmids carrying non-identical colicins (supplementary data S4).

**Table S1. Colicin distribution table.**

| **Class** | **Family** | **Genera** | **No. of strains carrying colicins on genome** | **No. of strains carrying colicins on plasmids** | **No. of strains carrying colicins on both genome & plasmid** |
| --- | --- | --- | --- | --- | --- |
| *Gammaproteobacteria* | *Enterobacteriaceae* | *Klebsiella* | 28 | 374 | 10 |
| *Gammaproteobacteria* | *Enterobacteriaceae* | *Salmonella* | 176 | 20 | 2 |
| *Gammaproteobacteria* | *Enterobacteriaceae* | *Escherichia* | 5 | 157 | 1 |
| *Gammaproteobacteria* | *Enterobacteriaceae* | *Enterobacter* | 22 | 4 | 2 |
| *Gammaproteobacteria* | *Yersiniaceae* | *Serratia* | 36 | 3 | 0 |
| *Gammaproteobacteria* | *Pseudomonadaceae* | *Pseudomonas* | 186 | 0 | 0 |
| *Gammaproteobacteria* | *Enterobacteriaceae* | *Shigella* | 0 | 37 | 0 |
| *Gammaproteobacteria* | *Morganellaceae* | *Providencia* | 12 | 0 | 0 |
| *Gammaproteobacteria* | *Morganellaceae* | *Proteus* | 26 | 0 | 0 |
| *Gammaproteobacteria* | *Yersiniaceae* | *Yersinia* | 91 | 0 | 0 |
| *Gammaproteobacteria* | *Morganellaceae* | *Morganella* | 7 | 2 | 0 |
| *Gammaproteobacteria* | *Pectobacteriaceae* | *Pectobacterium* | 7 | 0 | 0 |
| *Gammaproteobacteria* | *Enterobacteriaceae* | *Citrobacter* | 0 | 4 | 3 |
| *Gammaproteobacteria* | *Vibrionaceae* | *Vibrio* | 5 | 1 | 0 |
| *Gammaproteobacteria* | *Morganellaceae* | *Photorhabdus* | 5 | 0 | 0 |
| *Gammaproteobacteria* | *Yersiniaceae* | *Rahnella* | 3 | 0 | 0 |
| *Gammaproteobacteria* | *Enterobacteriaceae* | *Leclercia* | 2 | 0 | 0 |
| *Gammaproteobacteria* | *Erwiniaceae* | *Mixta* | 2 | 0 | 0 |
| *Gammaproteobacteria* | *Morganellaceae* | *Xenorhabdus* | 2 | 0 | 0 |
| *Gammaproteobacteria* | *Erwiniaceae* | *Erwinia* | 1 | 1 | 0 |
| *Gammaproteobacteria* | *Enterobacteriaceae* | *Edwardsiella* | 0 | 2 | 0 |
| *Betaproteobacteria* | *Burkholderiaceae* | *Burkholderia* | 1 | 0 | 0 |
| *Gammaproteobacteria* | *Enterobacteriaceae* | *Gibbsiella* | 1 | 0 | 0 |
| *Gammaproteobacteria* | *Pasteurellaceae* | *Actinobacillus* | 0 | 1 | 0 |
| *Gammaproteobacteria* | *Enterobacteriaceae* | *Hafnia* | 0 | 1 | 0 |
| *Gammaproteobacteria* | *Erwiniaceae* | *Pantoea* | 0 | 1 | 0 |
| *Gammaproteobacteria* | *Enterobacteriaceae* | *Raoultella* | 0 | 1 | 0 |
| *Gammaproteobacteria* | *Enterobacteriaceae* | *Obesumbacterium* | 1 | 0 | 0 |
|  |  | *Lonsdalea* | 1 | 0 | 0 |
|  |  | *Tolypothrix* | 1 | 0 | 0 |
