## Supplementary file S1 for "Modeling colicin operons as genetic arms in plasmid-genome conflicts"

| **Strain** | **Gene** | **Nuclease (Nucleotide)** | **Nuclease (Protein)** | **Immunity (Nucleotide)** | **Immunity (Protein)** | **Lysis (Nucleotide)** | **Lysis (protein)** | **operon** | **Nuclease + immunity (nucleotides)** | **Nuclease + immunity (protein)** | **Nuclease + immunity + lysis (protein)** |
| --- | --- | --- | --- | --- | --- | --- | --- | --- | --- | --- | --- |
| **Colicin E2**  **Escherichia coli strain BZB1011 E2C plasmid pColE2-P9**  <https://www.ncbi.nlm.nih.gov/nuccore/NZ_KY348421.1?report=graph> | ***ce2a***  **581 aa**  ***ce2i***  **86 aa**  ***ce2l***  **47 aa** | ATGAGCGGTGGCGATGGACGCGGCCATAACACGGGCGCGCATAGCACAAGTGGTAACATTAATGGTGGCCCGACCGGGCTTGGTGTAGGTGGTGGTGCTTCTGATGGCTCCGGATGGAGTTCGGAAAATAACCCGTGGGGTGGTGGTTCCGGTAGCGGCATTCACTGGGGTGGTGGTTCCGGTCATGGTAATGGCGGGGGGAATGGTAATTCCGGTGGTGGTTCGGGAACAGGCGGTAATCTGTCAGCAGTAGCTGCGCCAGTGGCATTTGGTTTTCCGGCACTTTCCACTCCAGGAGCTGGCGGTCTGGCGGTCAGTATTTCAGCGGGAGCATTATCGGCAGCTATTGCTGATATTATGGCTGCCCTGAAAGGACCGTTTAAATTTGGTCTTTGGGGGGTGGCTTTATATGGTGTATTGCCATCACAAATAGCGAAAGATGACCCCAATATGATGTCAAAGATTGTGACGTCATTACCCGCAGATGATATTACTGAATCACCTGTCAGTTCATTACCTCTCGATAAGGCAACAGTAAACGTAAATGTTCGTGTTGTTGATGATGTAAAAGACGAACGACAGAATATTTCGGTTGTTTCAGGTGTTCCGATGAGTGTTCCGGTGGTTGATGCAAAACCTACCGAACGTCCAGGTGTTTTTACGGCATCAATTCCAGGTGCACCTGTTCTGAATATTTCAGTTAATAACAGTACGCCAGAAGTACAGACATTAAGCCCAGGTGTTACAAATAATACTGATAAGGATGTTCGCCCGGCAGGATTTACTCAGGGTGGTAATACCAGGGATGCAGTTATTCGATTCCCGAAGGACAGCGGTCATAATGCCGTATATGTTTCAGTGAGTGATGTTCTTAGTCCTGACCAGGTAAAACAACGTCAGGATGAAGAAAATCGCCGTCAGCAGGAATGGGATGCTACGCATCCGGTTGAAGCGGCTGAGCGAAATTATGAACGCGCGCGTGCAGAGCTGAATCAGGCAAATGAAGATGTTGCCAGAAATCAGGAGCGACAGGCTAAAGCTGTTCAGGTTTATAATTCGCGTAAAAGCGAACTTGATGCAGCGAATAAAACTCTTGCTGATGCAATAGCTGAAATAAAACAATTTAATCGATTTGCCCATGACCCAATGGCTGGCGGTCACAGAATGTGGCAAATGGCCGGACTTAAAGCTCAGCGGGCGCAGACGGATGTAAATAATAAGCAGGCTGCATTTGATGCTGCTGCAAAAGAGAAGTCAGATGCTGATGCTGCATTAAGTGCCGCGCAGGAGCGCCGCAAACAGAAGGAAAATAAAGAAAAGGACGCTAAGGATAAATTAGATAAGGAGAGTAAACGGAATAAGCCAGGGAAGGCGACAGGTAAAGGTAAACCAGTTGGTGATAAATGGCTGGATGATGCAGGTAAAGATTCAGGAGCGCCAATTCCAGATCGCATTGCTGATAAGTTGCGTGATAAAGAATTTAAAAACTTTGACGATTTCCGGAAGAAATTCTGGGAAGAAGTGTCAAAAGATCCCGATCTTAGTAAGCAATTTAAAGGCAGTAATAAGACGAACATTCAAAAGGGAAAAGCACCTTTTGCAAGGAAGAAAGACCAAGTAGGTGGTAGGGAACGCTTTGAATTACATCATGATAAACCAATCAGTCAGGATGGTGGTGTCTATGATATGAATAATATCAGAGTGACCACACCTAAGCGACATATTGATATTCATCGGGGTAAGTAA | MSGGDGRGHNTGAHSTSGNINGGPTGLGVGGGASGSGWSSENNPWGGGSGSGIHWGGGSGHGNGGGNGNSGGGSGTGGNLSAVAAPVAFGFPALSTPGAGGLAVSISAGALSAAIADIMAALKGPFKFGLWGVALYGVLPSQIAKDDPNMMSKIVTSLPADDITESPVSSLPLDKATVNVNVRVVDDVKDERQNISVVSGVPMSVPVVDAKPTERPGVFTASIPGAPVLNISVNNSTPEVQTLSPGVTNNTDKDVRPAGFTQGGNTRDAVIRFPKDSGHNAVYVSVSDVLSPDQVKQRQDEENRRQQEWDATHPVEAAERNYERARAELNQANEDVARNQERQAKAVQVYNSRKSELDAANKTLADAIAEIKQFNRFAHDPMAGGHRMWQMAGLKAQRAQTDVNNKQAAFDAAAKEKSDADAALSAAQERRKQKENKEKDAKDKLDKESKRNKPGKATGKGKPVGDKWLDDAGKDSGAPIPDRIADKLRDKEFKNFDDFRKKFWEEVSKDPDLSKQFKGSNKTNIQKGKAPFARKKDQVGGRERFELHHDKPISQDGGVYDMNNIRVTTPKRHIDIHRGK | ATGGAACTGAAACATAGTATTAGTGATTATACCGAGGCTGAATTTCTGGAGTTTGTAAAAAAAATATGTAGAGCTGAAGGTGCTACTGAAGAGGATGACAATAAATTAGTGAGAGAGTTTGAGCGATTAACTGAGCACCCAGATGGTTCAGATCTGATTTATTATCCTCGCGATGACAGGGAAGATAGTCCTGAAGGGATTGTCAAGGAAATTAAAGAATGGCGAGCTGCTAACGGTAAGTCAGGATTTAAACAGGGCTGA | MELKHSISDYTEAEFLEFVKKICRAEGATEEDDNKLVREFERLTEHPDGSDLIYYPRDDREDSPEGIVKEIKEWRAANGKSGFKQG | ATGAAAAAAATAACAGGGATTATTTTATTGCTTCTTGCAGTCATTATTCTGTCTGCATGTCAGGCAAACTATATCCGGGATGTTCAGGGCGGGACCGTATCTCCGTCATCAACAGCTGAAGTGACCGGATTAGCAACGCAGTAA | MKKITGIILLLLAVIILSACQANYIRDVQGGTVSPSSTAEVTGLATQ | >E2 operon  ATGAGCGGTGGCGATGGACGCGGCCATAACACGGGCGCGCATAGCACAAGTGGTAACATTAATGGTGGCC  CGACCGGGCTTGGTGTAGGTGGTGGTGCTTCTGATGGCTCCGGATGGAGTTCGGAAAATAACCCGTGGGG  TGGTGGTTCCGGTAGCGGCATTCACTGGGGTGGTGGTTCCGGTCATGGTAATGGCGGGGGGAATGGTAAT  TCCGGTGGTGGTTCGGGAACAGGCGGTAATCTGTCAGCAGTAGCTGCGCCAGTGGCATTTGGTTTTCCGG  CACTTTCCACTCCAGGAGCTGGCGGTCTGGCGGTCAGTATTTCAGCGGGAGCATTATCGGCAGCTATTGC  TGATATTATGGCTGCCCTGAAAGGACCGTTTAAATTTGGTCTTTGGGGGGTGGCTTTATATGGTGTATTG  CCATCACAAATAGCGAAAGATGACCCCAATATGATGTCAAAGATTGTGACGTCATTACCCGCAGATGATA  TTACTGAATCACCTGTCAGTTCATTACCTCTCGATAAGGCAACAGTAAACGTAAATGTTCGTGTTGTTGA  TGATGTAAAAGACGAACGACAGAATATTTCGGTTGTTTCAGGTGTTCCGATGAGTGTTCCGGTGGTTGAT  GCAAAACCTACCGAACGTCCAGGTGTTTTTACGGCATCAATTCCAGGTGCACCTGTTCTGAATATTTCAG  TTAATAACAGTACGCCAGAAGTACAGACATTAAGCCCAGGTGTTACAAATAATACTGATAAGGATGTTCG  CCCGGCAGGATTTACTCAGGGTGGTAATACCAGGGATGCAGTTATTCGATTCCCGAAGGACAGCGGTCAT  AATGCCGTATATGTTTCAGTGAGTGATGTTCTTAGTCCTGACCAGGTAAAACAACGTCAGGATGAAGAAA  ATCGCCGTCAGCAGGAATGGGATGCTACGCATCCGGTTGAAGCGGCTGAGCGAAATTATGAACGCGCGCG  TGCAGAGCTGAATCAGGCAAATGAAGATGTTGCCAGAAATCAGGAGCGACAGGCTAAAGCTGTTCAGGTT  TATAATTCGCGTAAAAGCGAACTTGATGCAGCGAATAAAACTCTTGCTGATGCAATAGCTGAAATAAAAC  AATTTAATCGATTTGCCCATGACCCAATGGCTGGCGGTCACAGAATGTGGCAAATGGCCGGACTTAAAGC  TCAGCGGGCGCAGACGGATGTAAATAATAAGCAGGCTGCATTTGATGCTGCTGCAAAAGAGAAGTCAGAT  GCTGATGCTGCATTAAGTGCCGCGCAGGAGCGCCGCAAACAGAAGGAAAATAAAGAAAAGGACGCTAAGG  ATAAATTAGATAAGGAGAGTAAACGGAATAAGCCAGGGAAGGCGACAGGTAAAGGTAAACCAGTTGGTGA  TAAATGGCTGGATGATGCAGGTAAAGATTCAGGAGCGCCAATTCCAGATCGCATTGCTGATAAGTTGCGT  GATAAAGAATTTAAAAACTTTGACGATTTCCGGAAGAAATTCTGGGAAGAAGTGTCAAAAGATCCCGATC  TTAGTAAGCAATTTAAAGGCAGTAATAAGACGAACATTCAAAAGGGAAAAGCACCTTTTGCAAGGAAGAA  AGACCAAGTAGGTGGTAGGGAACGCTTTGAATTACATCATGATAAACCAATCAGTCAGGATGGTGGTGTC  TATGATATGAATAATATCAGAGTGACCACACCTAAGCGACATATTGATATTCATCGGGGTAAGTAAAAAT  GGAACTGAAACATAGTATTAGTGATTATACCGAGGCTGAATTTCTGGAGTTTGTAAAAAAAATATGTAGA  GCTGAAGGTGCTACTGAAGAGGATGACAATAAATTAGTGAGAGAGTTTGAGCGATTAACTGAGCACCCAG  ATGGTTCAGATCTGATTTATTATCCTCGCGATGACAGGGAAGATAGTCCTGAAGGGATTGTCAAGGAAAT  TAAAGAATGGCGAGCTGCTAACGGTAAGTCAGGATTTAAACAGGGCTGAAATATGAATGCCGGTTGTTTA  TGGATGAATGGCTGGCATTCTTTCACAACAAGGAGTCGTTATGAAAAAAATAACAGGGATTATTTTATTG  CTTCTTGCAGTCATTATTCTGTCTGCATGTCAGGCAAACTATATCCGGGATGTTCAGGGCGGGACCGTAT  CTCCGTCATCAACAGCTGAAGTGACCGGATTAGCAACGCAGTAA | ATGAGCGGTGGCGATGGACGCGGCCATAACACGGGCGCGCATAGCACAAGTGGTAACATTAATGGTGGCCCGACCGGGCTTGGTGTAGGTGGTGGTGCTTCTGATGGCTCCGGATGGAGTTCGGAAAATAACCCGTGGGGTGGTGGTTCCGGTAGCGGCATTCACTGGGGTGGTGGTTCCGGTCATGGTAATGGCGGGGGGAATGGTAATTCCGGTGGTGGTTCGGGAACAGGCGGTAATCTGTCAGCAGTAGCTGCGCCAGTGGCATTTGGTTTTCCGGCACTTTCCACTCCAGGAGCTGGCGGTCTGGCGGTCAGTATTTCAGCGGGAGCATTATCGGCAGCTATTGCTGATATTATGGCTGCCCTGAAAGGACCGTTTAAATTTGGTCTTTGGGGGGTGGCTTTATATGGTGTATTGCCATCACAAATAGCGAAAGATGACCCCAATATGATGTCAAAGATTGTGACGTCATTACCCGCAGATGATATTACTGAATCACCTGTCAGTTCATTACCTCTCGATAAGGCAACAGTAAACGTAAATGTTCGTGTTGTTGATGATGTAAAAGACGAACGACAGAATATTTCGGTTGTTTCAGGTGTTCCGATGAGTGTTCCGGTGGTTGATGCAAAACCTACCGAACGTCCAGGTGTTTTTACGGCATCAATTCCAGGTGCACCTGTTCTGAATATTTCAGTTAATAACAGTACGCCAGAAGTACAGACATTAAGCCCAGGTGTTACAAATAATACTGATAAGGATGTTCGCCCGGCAGGATTTACTCAGGGTGGTAATACCAGGGATGCAGTTATTCGATTCCCGAAGGACAGCGGTCATAATGCCGTATATGTTTCAGTGAGTGATGTTCTTAGTCCTGACCAGGTAAAACAACGTCAGGATGAAGAAAATCGCCGTCAGCAGGAATGGGATGCTACGCATCCGGTTGAAGCGGCTGAGCGAAATTATGAACGCGCGCGTGCAGAGCTGAATCAGGCAAATGAAGATGTTGCCAGAAATCAGGAGCGACAGGCTAAAGCTGTTCAGGTTTATAATTCGCGTAAAAGCGAACTTGATGCAGCGAATAAAACTCTTGCTGATGCAATAGCTGAAATAAAACAATTTAATCGATTTGCCCATGACCCAATGGCTGGCGGTCACAGAATGTGGCAAATGGCCGGACTTAAAGCTCAGCGGGCGCAGACGGATGTAAATAATAAGCAGGCTGCATTTGATGCTGCTGCAAAAGAGAAGTCAGATGCTGATGCTGCATTAAGTGCCGCGCAGGAGCGCCGCAAACAGAAGGAAAATAAAGAAAAGGACGCTAAGGATAAATTAGATAAGGAGAGTAAACGGAATAAGCCAGGGAAGGCGACAGGTAAAGGTAAACCAGTTGGTGATAAATGGCTGGATGATGCAGGTAAAGATTCAGGAGCGCCAATTCCAGATCGCATTGCTGATAAGTTGCGTGATAAAGAATTTAAAAACTTTGACGATTTCCGGAAGAAATTCTGGGAAGAAGTGTCAAAAGATCCCGATCTTAGTAAGCAATTTAAAGGCAGTAATAAGACGAACATTCAAAAGGGAAAAGCACCTTTTGCAAGGAAGAAAGACCAAGTAGGTGGTAGGGAACGCTTTGAATTACATCATGATAAACCAATCAGTCAGGATGGTGGTGTCTATGATATGAATAATATCAGAGTGACCACACCTAAGCGACATATTGATATTCATCGGGGTAAGTAA  ATGGAACTGAAACATAGTATTAGTGATTATACCGAGGCTGAATTTCTGGAGTTTGTAAAAAAAATATGTAGAGCTGAAGGTGCTACTGAAGAGGATGACAATAAATTAGTGAGAGAGTTTGAGCGATTAACTGAGCACCCAGATGGTTCAGATCTGATTTATTATCCTCGCGATGACAGGGAAGATAGTCCTGAAGGGATTGTCAAGGAAATTAAAGAATGGCGAGCTGCTAACGGTAAGTCAGGATTTAAACAGGGCTGA | MSGGDGRGHNTGAHSTSGNINGGPTGLGVGGGASGSGWSSENNPWGGGSGSGIHWGGGSGHGNGGGNGNSGGGSGTGGNLSAVAAPVAFGFPALSTPGAGGLAVSISAGALSAAIADIMAALKGPFKFGLWGVALYGVLPSQIAKDDPNMMSKIVTSLPADDITESPVSSLPLDKATVNVNVRVVDDVKDERQNISVVSGVPMSVPVVDAKPTERPGVFTASIPGAPVLNISVNNSTPEVQTLSPGVTNNTDKDVRPAGFTQGGNTRDAVIRFPKDSGHNAVYVSVSDVLSPDQVKQRQDEENRRQQEWDATHPVEAAERNYERARAELNQANEDVARNQERQAKAVQVYNSRKSELDAANKTLADAIAEIKQFNRFAHDPMAGGHRMWQMAGLKAQRAQTDVNNKQAAFDAAAKEKSDADAALSAAQERRKQKENKEKDAKDKLDKESKRNKPGKATGKGKPVGDKWLDDAGKDSGAPIPDRIADKLRDKEFKNFDDFRKKFWEEVSKDPDLSKQFKGSNKTNIQKGKAPFARKKDQVGGRERFELHHDKPISQDGGVYDMNNIRVTTPKRHIDIHRGK  MELKHSISDYTEAEFLEFVKKICRAEGATEEDDNKLVREFERLTEHPDGSDLIYYPRDDREDSPEGIVKEIKEWRAANGKSGFKQG | MSGGDGRGHNTGAHSTSGNINGGPTGLGVGGGASGSGWSSENNPWGGGSGSGIHWGGGSGHGNGGGNGNSGGGSGTGGNLSAVAAPVAFGFPALSTPGAGGLAVSISAGALSAAIADIMAALKGPFKFGLWGVALYGVLPSQIAKDDPNMMSKIVTSLPADDITESPVSSLPLDKATVNVNVRVVDDVKDERQNISVVSGVPMSVPVVDAKPTERPGVFTASIPGAPVLNISVNNSTPEVQTLSPGVTNNTDKDVRPAGFTQGGNTRDAVIRFPKDSGHNAVYVSVSDVLSPDQVKQRQDEENRRQQEWDATHPVEAAERNYERARAELNQANEDVARNQERQAKAVQVYNSRKSELDAANKTLADAIAEIKQFNRFAHDPMAGGHRMWQMAGLKAQRAQTDVNNKQAAFDAAAKEKSDADAALSAAQERRKQKENKEKDAKDKLDKESKRNKPGKATGKGKPVGDKWLDDAGKDSGAPIPDRIADKLRDKEFKNFDDFRKKFWEEVSKDPDLSKQFKGSNKTNIQKGKAPFARKKDQVGGRERFELHHDKPISQDGGVYDMNNIRVTTPKRHIDIHRGK  MELKHSISDYTEAEFLEFVKKICRAEGATEEDDNKLVREFERLTEHPDGSDLIYYPRDDREDSPEGIVKEIKEWRAANGKSGFKQG  MKKITGIILLLLAVIILSACQANYIRDVQGGTVSPSSTAEVTGLATQ |
| Colicin E3  Escherichia coli strain BZB2106 plasmid pColE3-CA38, complete sequence  <https://www.ncbi.nlm.nih.gov/nuccore/NZ_KM287568.1?report=graph> | Colicin E3  (551 aa)  Immunity  (85 aa)  Immunity  (85 aa)  Lysis  (47 aa) | ATGAGCGGTGGCGATGGACGCGGCCATAACACGGGCGCGCATAGCACAAGTGGTAACATTAATGGTGGCC  CGACCGGGCTTGGTGTAGGTGGTGGTGCTTCTGATGGCTCCGGATGGAGTTCGGAAAATAACCCGTGGGG  TGGTGGTTCCGGTAGCGGCATTCACTGGGGTGGTGGTTCCGGTCATGGTAATGGCGGGGGGAATGGTAAT  TCCGGTGGTGGTTCGGGAACAGGCGGTAATCTGTCAGCAGTAGCTGCGCCAGTGGCATTTGGTTTTCCGG  CACTTTCCACTCCAGGAGCTGGCGGTCTGGCGGTCAGTATTTCAGCGGGAGCATTATCGGCAGCTATTGC  TGATATTATGGCTGCCCTGAAAGGACCGTTTAAATTTGGTCTTTGGGGGGTGGCTTTATATGGTGTATTG  CCATCACAAATAGCGAAAGATGACCCCAATATGATGTCAAAGATTGTGACGTCATTACCCGCAGATGATA  TTACTGAATCACCTGTCAGTTCATTACCTCTCGATAAGGCAACAGTAAACGTAAATGTTCGTGTTGTTGA  TGATGTAAAAGACGAGCGACAGAATATTTCGGTTGTTTCAGGTGTTCCGATGAGTGTTCCGGTGGTTGAT  GCAAAACCTACCGAACGTCCGGGTGTTTTTACGGCATCAATTCCAGGTGCACCTGTTCTGAATATTTCAG  TTAATAACAGTACGCCAGCAGTACAGACATTAAGCCCAGGTGTTACAAATAATACTGATAAGGATGTTCG  CCCGGCAGGATTTACTCAGGGTGGTAATACCAGGGATGCAGTTATTCGATTCCCGAAGGACAGCGGTCAT  AATGCCGTATATGTTTCAGTGAGTGATGTTCTTAGCCCTGACCAGGTAAAACAACGTCAAGATGAAGAAA  ATCGCCGTCAGCAGGAATGGGATGCTACGCATCCGGTTGAAGCGGCTGAGCGAAATTATGAACGCGCGCG  TGCAGAGCTGAATCAGGCAAATGAAGATGTTGCCAGAAATCAGGAGCGACAGGCTAAAGCTGTTCAGGTT  TATAATTCGCGTAAAAGCGAACTTGATGCAGCGAATAAAACTCTTGCTGATGCAATAGCTGAAATAAAAC  AATTTAATCGATTTGCCCATGACCCAATGGCTGGCGGTCACAGAATGTGGCAAATGGCCGGGCTTAAAGC  CCAGCGGGCGCAGACGGATGTAAATAATAAGCAGGCTGCATTTGATGCTGCTGCAAAAGAGAAGTCAGAT  GCTGATGCTGCATTGAGTTCTGCTATGGAAAGCAGGAAGAAGAAAGAAGATAAGAAAAGGAGTGCTGAAA  ATAATTTAAACGATGAAAAGAATAAGCCCAGAAAAGGTTTTAAAGATTACGGGCATGATTATCATCCAGC  TCCGAAAACTGAGAATATTAAAGGGCTTGGTGATCTTAAGCCTGGGATACCAAAAACACCAAAGCAGAAT  GGTGGTGGAAAACGCAAGCGCTGGACTGGAGATAAAGGGCGTAAGATTTATGAGTGGGATTCTCAGCATG  GTGAGCTTGAGGGGTATCGTGCCAGTGATGGTCAGCATCTTGGCTCATTTGACCCTAAAACAGGCAATCA  GTTGAAAGGTCCAGATCCGAAACGAAATATCAAGAAATATCTTTGA | MSGGDGRGHNTGAHSTSGNINGGPTGLGVGGGASDGSGWSSENNPWGGGSGSGIHWGGGSGHGNGGGNGN  SGGGSGTGGNLSAVAAPVAFGFPALSTPGAGGLAVSISAGALSAAIADIMAALKGPFKFGLWGVALYGVL  PSQIAKDDPNMMSKIVTSLPADDITESPVSSLPLDKATVNVNVRVVDDVKDERQNISVVSGVPMSVPVVD  AKPTERPGVFTASIPGAPVLNISVNNSTPAVQTLSPGVTNNTDKDVRPAGFTQGGNTRDAVIRFPKDSGH  NAVYVSVSDVLSPDQVKQRQDEENRRQQEWDATHPVEAAERNYERARAELNQANEDVARNQERQAKAVQV  YNSRKSELDAANKTLADAIAEIKQFNRFAHDPMAGGHRMWQMAGLKAQRAQTDVNNKQAAFDAAAKEKSD  ADAALSSAMESRKKKEDKKRSAENNLNDEKNKPRKGFKDYGHDYHPAPKTENIKGLGDLKPGIPKTPKQN  GGGKRKRWTGDKGRKIYEWDSQHGELEGYRASDGQHLGSFDPKTGNQLKGPDPKRNIKKYL | ATGGGACTTAAATTGGATTTAACTTGGTTTGATAAAAGTACAGAAGATTTTAAGGGTGAGGAGTATTCAA  AAGATTTTGGAGATGACGGTTCAGTTATGGAAAGTCTAGGTGTGCCTTTTAAGGATAATGTTAATAACGG  TTGCTTTGATGTTATAGCTGAATGGGTACCTTTGCTACAACCATACTTTAATCATCAAATTGATATTTCC  GATAATGAGTATTTTGTTTCGTTTGATTATCGTGATGGTGATTGGTGA  GTGGAGCTAAAAAAAAGTATTGGTGATTACACTGAAACCGAATTCAAAAAATTTATTGAAGACATCATCA  ATTGTGAAGGTGATGAAAAAAAACAGGATGATAACCTCGAGTATTTTATAAATGTTACTGAGCATCCTAG  TGGTTCTGATCTGATTTATTACCCAGAAGGTAATAATGATGGTAGCCCTGAAGGTGTTATTAAAGAGATT  AAAGAATGGCGAGCCGCTAACGGTAAGTCAGGATTTAAACAGGGCTGA | MGLKLDLTWFDKSTEDFKGEEYSKDFGDDGSVMESLGVPFKDNVNNGCFDVIAEWVPLLQPYFNHQIDIS  DNEYFVSFDYRDGDW  MELKKSIGDYTETEFKKFIEDIINCEGDEKKQDDNLEYFINVTEHPSGSDLIYYPEGNNDGSPEGVIKEI  KEWRAANGKSGFKQG | ATGAAAAAAATAACAGGGATTATTTTATTGCTTCTTGCAGTCATTATTCTGTCTGCATGTCAGGCAAACT  ATATCCGGGATGTTCAGGGCGGGACCGTATCTCCGTCATCAACAGCTGAAGTGACCGGATTAGCAACGCA  GTAA | MKKITGIILLLLAVIILSACQANYIRDVQGGTVSPSSTAEVTGLATQ | ATGAGCGGTGGCGATGGACGCGGCCATAACACGGGCGCGCATAGCACAAGTGGTAACATTAATGGTGGCC  CGACCGGGCTTGGTGTAGGTGGTGGTGCTTCTGATGGCTCCGGATGGAGTTCGGAAAATAACCCGTGGGG  TGGTGGTTCCGGTAGCGGCATTCACTGGGGTGGTGGTTCCGGTCATGGTAATGGCGGGGGGAATGGTAAT  TCCGGTGGTGGTTCGGGAACAGGCGGTAATCTGTCAGCAGTAGCTGCGCCAGTGGCATTTGGTTTTCCGG  CACTTTCCACTCCAGGAGCTGGCGGTCTGGCGGTCAGTATTTCAGCGGGAGCATTATCGGCAGCTATTGC  TGATATTATGGCTGCCCTGAAAGGACCGTTTAAATTTGGTCTTTGGGGGGTGGCTTTATATGGTGTATTG  CCATCACAAATAGCGAAAGATGACCCCAATATGATGTCAAAGATTGTGACGTCATTACCCGCAGATGATA  TTACTGAATCACCTGTCAGTTCATTACCTCTCGATAAGGCAACAGTAAACGTAAATGTTCGTGTTGTTGA  TGATGTAAAAGACGAGCGACAGAATATTTCGGTTGTTTCAGGTGTTCCGATGAGTGTTCCGGTGGTTGAT  GCAAAACCTACCGAACGTCCGGGTGTTTTTACGGCATCAATTCCAGGTGCACCTGTTCTGAATATTTCAG  TTAATAACAGTACGCCAGCAGTACAGACATTAAGCCCAGGTGTTACAAATAATACTGATAAGGATGTTCG  CCCGGCAGGATTTACTCAGGGTGGTAATACCAGGGATGCAGTTATTCGATTCCCGAAGGACAGCGGTCAT  AATGCCGTATATGTTTCAGTGAGTGATGTTCTTAGCCCTGACCAGGTAAAACAACGTCAAGATGAAGAAA  ATCGCCGTCAGCAGGAATGGGATGCTACGCATCCGGTTGAAGCGGCTGAGCGAAATTATGAACGCGCGCG  TGCAGAGCTGAATCAGGCAAATGAAGATGTTGCCAGAAATCAGGAGCGACAGGCTAAAGCTGTTCAGGTT  TATAATTCGCGTAAAAGCGAACTTGATGCAGCGAATAAAACTCTTGCTGATGCAATAGCTGAAATAAAAC  AATTTAATCGATTTGCCCATGACCCAATGGCTGGCGGTCACAGAATGTGGCAAATGGCCGGGCTTAAAGC  CCAGCGGGCGCAGACGGATGTAAATAATAAGCAGGCTGCATTTGATGCTGCTGCAAAAGAGAAGTCAGAT  GCTGATGCTGCATTGAGTTCTGCTATGGAAAGCAGGAAGAAGAAAGAAGATAAGAAAAGGAGTGCTGAAA  ATAATTTAAACGATGAAAAGAATAAGCCCAGAAAAGGTTTTAAAGATTACGGGCATGATTATCATCCAGC  TCCGAAAACTGAGAATATTAAAGGGCTTGGTGATCTTAAGCCTGGGATACCAAAAACACCAAAGCAGAAT  GGTGGTGGAAAACGCAAGCGCTGGACTGGAGATAAAGGGCGTAAGATTTATGAGTGGGATTCTCAGCATG  GTGAGCTTGAGGGGTATCGTGCCAGTGATGGTCAGCATCTTGGCTCATTTGACCCTAAAACAGGCAATCA  GTTGAAAGGTCCAGATCCGAAACGAAATATCAAGAAATATCTTTGAGAGGAAGTTATGGGACTTAAATTG  GATTTAACTTGGTTTGATAAAAGTACAGAAGATTTTAAGGGTGAGGAGTATTCAAAAGATTTTGGAGATG  ACGGTTCAGTTATGGAAAGTCTAGGTGTGCCTTTTAAGGATAATGTTAATAACGGTTGCTTTGATGTTAT  AGCTGAATGGGTACCTTTGCTACAACCATACTTTAATCATCAAATTGATATTTCCGATAATGAGTATTTT  GTTTCGTTTGATTATCGTGATGGTGATTGGTGATCAAATATTATCAGGGATGAGTTGATATACGGGCTTC  TAGTGTTCATGGATGAACGCTGGAGCCTCCAAATGTAGAAATGTTATATTTTTTATTGAGTTCTTGGTTA  TAATTGCTCCGCAATGATTTAAATAAGCATTATTTAAAACATTCTCAGGAGAGGTGAAGGTGGAGCTAAA  AAAAAGTATTGGTGATTACACTGAAACCGAATTCAAAAAATTTATTGAAGACATCATCAATTGTGAAGGT  GATGAAAAAAAACAGGATGATAACCTCGAGTATTTTATAAATGTTACTGAGCATCCTAGTGGTTCTGATC  TGATTTATTACCCAGAAGGTAATAATGATGGTAGCCCTGAAGGTGTTATTAAAGAGATTAAAGAATGGCG  AGCCGCTAACGGTAAGTCAGGATTTAAACAGGGCTGAAATATGAATGCCGGTTGTTTATGGATGAATGGC  TGGCATTCTTTCACAACAAGGAGTCGTTATGAAAAAAATAACAGGGATTATTTTATTGCTTCTTGCAGTC  ATTATTCTGTCTGCATGTCAGGCAAACTATATCCGGGATGTTCAGGGCGGGACCGTATCTCCGTCATCAA  CAGCTGAAGTGACCGGATTAGCAACGCAGTAA | ATGAGCGGTGGCGATGGACGCGGCCATAACACGGGCGCGCATAGCACAAGTGGTAACATTAATGGTGGCC  CGACCGGGCTTGGTGTAGGTGGTGGTGCTTCTGATGGCTCCGGATGGAGTTCGGAAAATAACCCGTGGGG  TGGTGGTTCCGGTAGCGGCATTCACTGGGGTGGTGGTTCCGGTCATGGTAATGGCGGGGGGAATGGTAAT  TCCGGTGGTGGTTCGGGAACAGGCGGTAATCTGTCAGCAGTAGCTGCGCCAGTGGCATTTGGTTTTCCGG  CACTTTCCACTCCAGGAGCTGGCGGTCTGGCGGTCAGTATTTCAGCGGGAGCATTATCGGCAGCTATTGC  TGATATTATGGCTGCCCTGAAAGGACCGTTTAAATTTGGTCTTTGGGGGGTGGCTTTATATGGTGTATTG  CCATCACAAATAGCGAAAGATGACCCCAATATGATGTCAAAGATTGTGACGTCATTACCCGCAGATGATA  TTACTGAATCACCTGTCAGTTCATTACCTCTCGATAAGGCAACAGTAAACGTAAATGTTCGTGTTGTTGA  TGATGTAAAAGACGAGCGACAGAATATTTCGGTTGTTTCAGGTGTTCCGATGAGTGTTCCGGTGGTTGAT  GCAAAACCTACCGAACGTCCGGGTGTTTTTACGGCATCAATTCCAGGTGCACCTGTTCTGAATATTTCAG  TTAATAACAGTACGCCAGCAGTACAGACATTAAGCCCAGGTGTTACAAATAATACTGATAAGGATGTTCG  CCCGGCAGGATTTACTCAGGGTGGTAATACCAGGGATGCAGTTATTCGATTCCCGAAGGACAGCGGTCAT  AATGCCGTATATGTTTCAGTGAGTGATGTTCTTAGCCCTGACCAGGTAAAACAACGTCAAGATGAAGAAA  ATCGCCGTCAGCAGGAATGGGATGCTACGCATCCGGTTGAAGCGGCTGAGCGAAATTATGAACGCGCGCG  TGCAGAGCTGAATCAGGCAAATGAAGATGTTGCCAGAAATCAGGAGCGACAGGCTAAAGCTGTTCAGGTT  TATAATTCGCGTAAAAGCGAACTTGATGCAGCGAATAAAACTCTTGCTGATGCAATAGCTGAAATAAAAC  AATTTAATCGATTTGCCCATGACCCAATGGCTGGCGGTCACAGAATGTGGCAAATGGCCGGGCTTAAAGC  CCAGCGGGCGCAGACGGATGTAAATAATAAGCAGGCTGCATTTGATGCTGCTGCAAAAGAGAAGTCAGAT  GCTGATGCTGCATTGAGTTCTGCTATGGAAAGCAGGAAGAAGAAAGAAGATAAGAAAAGGAGTGCTGAAA  ATAATTTAAACGATGAAAAGAATAAGCCCAGAAAAGGTTTTAAAGATTACGGGCATGATTATCATCCAGC  TCCGAAAACTGAGAATATTAAAGGGCTTGGTGATCTTAAGCCTGGGATACCAAAAACACCAAAGCAGAAT  GGTGGTGGAAAACGCAAGCGCTGGACTGGAGATAAAGGGCGTAAGATTTATGAGTGGGATTCTCAGCATG  GTGAGCTTGAGGGGTATCGTGCCAGTGATGGTCAGCATCTTGGCTCATTTGACCCTAAAACAGGCAATCA  GTTGAAAGGTCCAGATCCGAAACGAAATATCAAGAAATATCTTTGA  ATGGGACTTAAATTGGATTTAACTTGGTTTGATAAAAGTACAGAAGATTTTAAGGGTGAGGAGTATTCAA  AAGATTTTGGAGATGACGGTTCAGTTATGGAAAGTCTAGGTGTGCCTTTTAAGGATAATGTTAATAACGG  TTGCTTTGATGTTATAGCTGAATGGGTACCTTTGCTACAACCATACTTTAATCATCAAATTGATATTTCC  GATAATGAGTATTTTGTTTCGTTTGATTATCGTGATGGTGATTGGTGA  GTGGAGCTAAAAAAAAGTATTGGTGATTACACTGAAACCGAATTCAAAAAATTTATTGAAGACATCATCA  ATTGTGAAGGTGATGAAAAAAAACAGGATGATAACCTCGAGTATTTTATAAATGTTACTGAGCATCCTAG  TGGTTCTGATCTGATTTATTACCCAGAAGGTAATAATGATGGTAGCCCTGAAGGTGTTATTAAAGAGATT  AAAGAATGGCGAGCCGCTAACGGTAAGTCAGGATTTAAACAGGGCTGA | MSGGDGRGHNTGAHSTSGNINGGPTGLGVGGGASDGSGWSSENNPWGGGSGSGIHWGGGSGHGNGGGNGN  SGGGSGTGGNLSAVAAPVAFGFPALSTPGAGGLAVSISAGALSAAIADIMAALKGPFKFGLWGVALYGVL  PSQIAKDDPNMMSKIVTSLPADDITESPVSSLPLDKATVNVNVRVVDDVKDERQNISVVSGVPMSVPVVD  AKPTERPGVFTASIPGAPVLNISVNNSTPAVQTLSPGVTNNTDKDVRPAGFTQGGNTRDAVIRFPKDSGH  NAVYVSVSDVLSPDQVKQRQDEENRRQQEWDATHPVEAAERNYERARAELNQANEDVARNQERQAKAVQV  YNSRKSELDAANKTLADAIAEIKQFNRFAHDPMAGGHRMWQMAGLKAQRAQTDVNNKQAAFDAAAKEKSD  ADAALSSAMESRKKKEDKKRSAENNLNDEKNKPRKGFKDYGHDYHPAPKTENIKGLGDLKPGIPKTPKQN  GGGKRKRWTGDKGRKIYEWDSQHGELEGYRASDGQHLGSFDPKTGNQLKGPDPKRNIKKYL  MGLKLDLTWFDKSTEDFKGEEYSKDFGDDGSVMESLGVPFKDNVNNGCFDVIAEWVPLLQPYFNHQIDIS  DNEYFVSFDYRDGDW  MELKKSIGDYTETEFKKFIEDIINCEGDEKKQDDNLEYFINVTEHPSGSDLIYYPEGNNDGSPEGVIKEI  KEWRAANGKSGFKQG | MSGGDGRGHNTGAHSTSGNINGGPTGLGVGGGASDGSGWSSENNPWGGGSGSGIHWGGGSGHGNGGGNGN  SGGGSGTGGNLSAVAAPVAFGFPALSTPGAGGLAVSISAGALSAAIADIMAALKGPFKFGLWGVALYGVL  PSQIAKDDPNMMSKIVTSLPADDITESPVSSLPLDKATVNVNVRVVDDVKDERQNISVVSGVPMSVPVVD  AKPTERPGVFTASIPGAPVLNISVNNSTPAVQTLSPGVTNNTDKDVRPAGFTQGGNTRDAVIRFPKDSGH  NAVYVSVSDVLSPDQVKQRQDEENRRQQEWDATHPVEAAERNYERARAELNQANEDVARNQERQAKAVQV  YNSRKSELDAANKTLADAIAEIKQFNRFAHDPMAGGHRMWQMAGLKAQRAQTDVNNKQAAFDAAAKEKSD  ADAALSSAMESRKKKEDKKRSAENNLNDEKNKPRKGFKDYGHDYHPAPKTENIKGLGDLKPGIPKTPKQN  GGGKRKRWTGDKGRKIYEWDSQHGELEGYRASDGQHLGSFDPKTGNQLKGPDPKRNIKKYL  MGLKLDLTWFDKSTEDFKGEEYSKDFGDDGSVMESLGVPFKDNVNNGCFDVIAEWVPLLQPYFNHQIDIS  DNEYFVSFDYRDGDW  MELKKSIGDYTETEFKKFIEDIINCEGDEKKQDDNLEYFINVTEHPSGSDLIYYPEGNNDGSPEGVIKEI  KEWRAANGKSGFKQG  MKKITGIILLLLAVIILSACQANYIRDVQGGTVSPSSTAEVTGLATQ |
| Escherichia coli strain M15 plasmid D, complete sequence  [https://www.ncbi.nlm.nih.gov/nuccore/CP010217.1?report=graph&rid=EKBKRTVX016[CP010217.1]&tracks=[key:sequence_track,name:Sequence,display_name:Sequence,id:STD1,category:Sequence,annots:Sequence,ShowLabel:true][key:gene_model_track,CDSProductFeats:false][key:alignment_track,name:other%20alignments,annots:NG%20Alignments\|Refseq%20Alignments\|Gnomon%20Alignments\|Unnamed,shown:false]&v=1765:2347&appname=ncbiblast&link_loc=fromHSP](https://www.ncbi.nlm.nih.gov/nuccore/CP010217.1?report=graph&rid=EKBKRTVX016%5bCP010217.1%5d&tracks=%5bkey:sequence_track,name:Sequence,display_name:Sequence,id:STD1,category:Sequence,annots:Sequence,ShowLabel:true%5d%5bkey:gene_model_track,CDSProductFeats:false%5d%5bkey:alignment_track,name:other%20alignments,annots:NG%20Alignments\|Refseq%20Alignments\|Gnomon%20Alignments\|Unnamed,shown:false%5d&v=1765:2347&appname=ncbiblast&link_loc=fromHSP) | Colicin E4  (555 aa)  Immunity  (85 aa)  Lysis  (47 aa) | ATGAGTGGTGGAGATGGAAAAGGTCCGGGTAATACAGGCCTGGGGCATAGCGGCGGTAAA  CCCAGTGGGAATGTGAATGGCGGCCCAACCGGGCTTGGTGGTAGTGGTAACAGCGGTGGT  AATAATCCAAACAGTGGGCCGGGCTGGGGAACGACACATACTCCTAATGGGGATATTCAC  AACTACAATCCGGGTGAGTTTGGTAATGGCGGGAATAAACCTGGGGGCAATAGCGGTAAT  CATGATGGTCATGGGTCGGGAGTCCAGTCGGCAATGGCATTTGGTTTTCCGGCGCTGGCA  GCACCTGGTGCCGGAACACTGGGTATCGCTGTCTCTGGTGAGGCTTTATCTGCGGCAATA  GCGGATATCTTTGCAGCCCTGAAAGGCCCGTTTAAATTCAGTGCATGGGGTATTGCGCTT  TACGGCATTCTGCCATCTGAAATAGCAAAAGATGACCCGAAAATGATGTCAAAGATTGTG  ACGTCATTACCGGCAGAAACAGTGACGAATGTTCAGGTCAGTACACTTCCGCTGGACCAG  GCAACTGTCAGCGTGACGAAACGTGTGACAGATGTTGTGAAGGACACACGACAGCATATT  GCCGTAGTTGCAGGTGTGCCTATGAGTGTGCCTGTTGTGAATGCCAAACCAACACGTACT  CCAGGTGTATTTCACGCATCATTTCCTGGTGTCCCTTCTCTGACAGTCAGCACTGTCAAA  GGCCTTCCGGTGTCAACAACCCTTCCCCGTGGTATTATGGAGGATAAAGGCCGGACTGCC  GTTCCTGCGGGATTTACCTTTGGTGGTGGTTCACATGAAGCGGTGATTCGCTTTCCGAAA  GAAAGTGGGCAGAAGCCGGTTTATGTATCAGTGACAGATGTTCTTACCCCAGCACAGGTA  AAACAACGTCAGGATGAGGAAAAACGCCTCCAGCAGGAATGGAATGACGCACATCCGGTG  GAAGTGGCTGAACGCAATTATGAACAGGCCCGCGCAGAACTGAATCAGGCTAATAAAGAT  GTTGCCAGAAACCAGGAACGACAGGCTAAAGCTGTTCAGGTTTATAATTCTCGTAAAAGT  GAACTTGATGCAGCGAATAAAACTCTTGCTGATGCAAAGGCTGAAATAAAACAATTCGAG  CGATTTGCCCGAGAACCAATGGCTGCTGGTCACAGAATGTGGCAAATGGCAGGGCTTAAG  GCCCAGCGGGCACAGACGGATGTAAATAATAAGAAGGCTGCATTTGATGCTGCCGCAAAA  GAGAAGTCAGCAGCAGATGCTGCATTGAGTTCTGCGATGGAAAGCAGGAAGAAGAAAGAA  GATAAGAAAAGGAGTGCTGAAAATAAATTAAACGAAGAGAAAAATAAACGCCGAAAAGGC  ACTAAAGATTATGGCCATGATTATCATCCAGCTCCTAAAACTGAGGATATTAAAGGTTTG  GGGGAATTAAAAGAGGGGAGGCCAAAAACTCCAAAGCAAGGTGGTGGAGGTAAGCGTGCC  AGATGGTATGGAGATAAGGGGCGTAAAATTTATGAGTGGGACTCTCAGCATGGTGAGCTT  GAGGGCTATCGAGCCAGTGATGGTCAGCATCTTGGTTCATTTGACCCTAAAACCGGTAAG  CAGTTGAAAGGTCCAGATCCGAAACGAAATATTAAAAAGTATCTTTAA | MSGGDGKGPGNTGLGHSGGKPSGNVNGGPTGLGGSGNSGGNNPNSGPGWGTTHTPNGDIHNYNPGEFGNG  GNKPGGNSGNHDGHGSGVQSAMAFGFPALAAPGAGTLGIAVSGEALSAAIADIFAALKGPFKFSAWGIAL  YGILPSEIAKDDPKMMSKIVTSLPAETVTNVQVSTLPLDQATVSVTKRVTDVVKDTRQHIAVVAGVPMSV  PVVNAKPTRTPGVFHASFPGVPSLTVSTVKGLPVSTTLPRGIMEDKGRTAVPAGFTFGGGSHEAVIRFPK  ESGQKPVYVSVTDVLTPAQVKQRQDEEKRLQQEWNDAHPVEVAERNYEQARAELNQANKDVARNQERQAK  AVQVYNSRKSELDAANKTLADAKAEIKQFERFAREPMAAGHRMWQMAGLKAQRAQTDVNNKKAAFDAAAK  EKSAADAALSSAMESRKKKEDKKRSAENKLNEEKNKRRKGTKDYGHDYHPAPKTEDIKGLGELKEGRPKT  PKQGGGGKRARWYGDKGRKIYEWDSQHGELEGYRASDGQHLGSFDPKTGKQLKGPDPKRNIKKYL | ATGGGACTTAAATTACACATTCATTGGTTTGATAAGAAGAATGAGGAATTTAAAGGTGGT  GAGTATTCAAAGGATTTCGGTGATGATGGTTCGGTTATTGAACGTCTTGGAATGCCTTTA  AAGGATAATATTAATAACGGTTGGTTTGATGTTGAAAAGAGATGGGTGCCAATATTACAG  CCATACTTTAAAAATGTTATTGATATTAGTAAGTTCGATTACTTTGTTTCTTTCGATTAT  CGTGATGGTGACTGGTAA | MGLKLHIHWFDKKNEEFKGGEYSKDFGDDGSVIERLGMPLKDNINNGWFDVEKRWVPILQPYFKNVIDIS  KFDYFVSFDYRDGDW | ATGAAGAAAATAACAGGGATTATTTTGTTGCTTCTTGCAGCCATTATTCTGGCTGCATGT  CAGGCAAACTATATCCGTGATGTTCAGGGCGGGACTGTATCACCGTCATCCTCTGCTGAA  CTGACCGGAGTGGAAACGCAGTAA | MKKITGIILLLLAAIILAACQANYIRDVQGGTVSPSSSAELTGVETQ | ATGAGTGGTGGAGATGGAAAAGGTCCGGGTAATACAGGCCTGGGGCATAGCGGCGGTAAA  CCCAGTGGGAATGTGAATGGCGGCCCAACCGGGCTTGGTGGTAGTGGTAACAGCGGTGGT  AATAATCCAAACAGTGGGCCGGGCTGGGGAACGACACATACTCCTAATGGGGATATTCAC  AACTACAATCCGGGTGAGTTTGGTAATGGCGGGAATAAACCTGGGGGCAATAGCGGTAAT  CATGATGGTCATGGGTCGGGAGTCCAGTCGGCAATGGCATTTGGTTTTCCGGCGCTGGCA  GCACCTGGTGCCGGAACACTGGGTATCGCTGTCTCTGGTGAGGCTTTATCTGCGGCAATA  GCGGATATCTTTGCAGCCCTGAAAGGCCCGTTTAAATTCAGTGCATGGGGTATTGCGCTT  TACGGCATTCTGCCATCTGAAATAGCAAAAGATGACCCGAAAATGATGTCAAAGATTGTG  ACGTCATTACCGGCAGAAACAGTGACGAATGTTCAGGTCAGTACACTTCCGCTGGACCAG  GCAACTGTCAGCGTGACGAAACGTGTGACAGATGTTGTGAAGGACACACGACAGCATATT  GCCGTAGTTGCAGGTGTGCCTATGAGTGTGCCTGTTGTGAATGCCAAACCAACACGTACT  CCAGGTGTATTTCACGCATCATTTCCTGGTGTCCCTTCTCTGACAGTCAGCACTGTCAAA  GGCCTTCCGGTGTCAACAACCCTTCCCCGTGGTATTATGGAGGATAAAGGCCGGACTGCC  GTTCCTGCGGGATTTACCTTTGGTGGTGGTTCACATGAAGCGGTGATTCGCTTTCCGAAA  GAAAGTGGGCAGAAGCCGGTTTATGTATCAGTGACAGATGTTCTTACCCCAGCACAGGTA  AAACAACGTCAGGATGAGGAAAAACGCCTCCAGCAGGAATGGAATGACGCACATCCGGTG  GAAGTGGCTGAACGCAATTATGAACAGGCCCGCGCAGAACTGAATCAGGCTAATAAAGAT  GTTGCCAGAAACCAGGAACGACAGGCTAAAGCTGTTCAGGTTTATAATTCTCGTAAAAGT  GAACTTGATGCAGCGAATAAAACTCTTGCTGATGCAAAGGCTGAAATAAAACAATTCGAG  CGATTTGCCCGAGAACCAATGGCTGCTGGTCACAGAATGTGGCAAATGGCAGGGCTTAAG  GCCCAGCGGGCACAGACGGATGTAAATAATAAGAAGGCTGCATTTGATGCTGCCGCAAAA  GAGAAGTCAGCAGCAGATGCTGCATTGAGTTCTGCGATGGAAAGCAGGAAGAAGAAAGAA  GATAAGAAAAGGAGTGCTGAAAATAAATTAAACGAAGAGAAAAATAAACGCCGAAAAGGC  ACTAAAGATTATGGCCATGATTATCATCCAGCTCCTAAAACTGAGGATATTAAAGGTTTG  GGGGAATTAAAAGAGGGGAGGCCAAAAACTCCAAAGCAAGGTGGTGGAGGTAAGCGTGCC  AGATGGTATGGAGATAAGGGGCGTAAAATTTATGAGTGGGACTCTCAGCATGGTGAGCTT  GAGGGCTATCGAGCCAGTGATGGTCAGCATCTTGGTTCATTTGACCCTAAAACCGGTAAG  CAGTTGAAAGGTCCAGATCCGAAACGAAATATTAAAAAGTATCTTTAAGGTGATGTTATG  GGACTTAAATTACACATTCATTGGTTTGATAAGAAGAATGAGGAATTTAAAGGTGGTGAG  TATTCAAAGGATTTCGGTGATGATGGTTCGGTTATTGAACGTCTTGGAATGCCTTTAAAG  GATAATATTAATAACGGTTGGTTTGATGTTGAAAAGAGATGGGTGCCAATATTACAGCCA  TACTTTAAAAATGTTATTGATATTAGTAAGTTCGATTACTTTGTTTCTTTCGATTATCGT  GATGGTGACTGGTAATTAAACTATCAGGGACGAGTAATAAAGTGTTTGATTCAGGTGTTC  AAGGATGAATTCCTGAATCCATATATACAGGGAATCGTATGAAGAAAATAACAGGGATTA  TTTTGTTGCTTCTTGCAGCCATTATTCTGGCTGCATGTCAGGCAAACTATATCCGTGATG  TTCAGGGCGGGACTGTATCACCGTCATCCTCTGCTGAACTGACCGGAGTGGAAACGCAGT  AA | ATGAGTGGTGGAGATGGAAAAGGTCCGGGTAATACAGGCCTGGGGCATAGCGGCGGTAAA  CCCAGTGGGAATGTGAATGGCGGCCCAACCGGGCTTGGTGGTAGTGGTAACAGCGGTGGT  AATAATCCAAACAGTGGGCCGGGCTGGGGAACGACACATACTCCTAATGGGGATATTCAC  AACTACAATCCGGGTGAGTTTGGTAATGGCGGGAATAAACCTGGGGGCAATAGCGGTAAT  CATGATGGTCATGGGTCGGGAGTCCAGTCGGCAATGGCATTTGGTTTTCCGGCGCTGGCA  GCACCTGGTGCCGGAACACTGGGTATCGCTGTCTCTGGTGAGGCTTTATCTGCGGCAATA  GCGGATATCTTTGCAGCCCTGAAAGGCCCGTTTAAATTCAGTGCATGGGGTATTGCGCTT  TACGGCATTCTGCCATCTGAAATAGCAAAAGATGACCCGAAAATGATGTCAAAGATTGTG  ACGTCATTACCGGCAGAAACAGTGACGAATGTTCAGGTCAGTACACTTCCGCTGGACCAG  GCAACTGTCAGCGTGACGAAACGTGTGACAGATGTTGTGAAGGACACACGACAGCATATT  GCCGTAGTTGCAGGTGTGCCTATGAGTGTGCCTGTTGTGAATGCCAAACCAACACGTACT  CCAGGTGTATTTCACGCATCATTTCCTGGTGTCCCTTCTCTGACAGTCAGCACTGTCAAA  GGCCTTCCGGTGTCAACAACCCTTCCCCGTGGTATTATGGAGGATAAAGGCCGGACTGCC  GTTCCTGCGGGATTTACCTTTGGTGGTGGTTCACATGAAGCGGTGATTCGCTTTCCGAAA  GAAAGTGGGCAGAAGCCGGTTTATGTATCAGTGACAGATGTTCTTACCCCAGCACAGGTA  AAACAACGTCAGGATGAGGAAAAACGCCTCCAGCAGGAATGGAATGACGCACATCCGGTG  GAAGTGGCTGAACGCAATTATGAACAGGCCCGCGCAGAACTGAATCAGGCTAATAAAGAT  GTTGCCAGAAACCAGGAACGACAGGCTAAAGCTGTTCAGGTTTATAATTCTCGTAAAAGT  GAACTTGATGCAGCGAATAAAACTCTTGCTGATGCAAAGGCTGAAATAAAACAATTCGAG  CGATTTGCCCGAGAACCAATGGCTGCTGGTCACAGAATGTGGCAAATGGCAGGGCTTAAG  GCCCAGCGGGCACAGACGGATGTAAATAATAAGAAGGCTGCATTTGATGCTGCCGCAAAA  GAGAAGTCAGCAGCAGATGCTGCATTGAGTTCTGCGATGGAAAGCAGGAAGAAGAAAGAA  GATAAGAAAAGGAGTGCTGAAAATAAATTAAACGAAGAGAAAAATAAACGCCGAAAAGGC  ACTAAAGATTATGGCCATGATTATCATCCAGCTCCTAAAACTGAGGATATTAAAGGTTTG  GGGGAATTAAAAGAGGGGAGGCCAAAAACTCCAAAGCAAGGTGGTGGAGGTAAGCGTGCC  AGATGGTATGGAGATAAGGGGCGTAAAATTTATGAGTGGGACTCTCAGCATGGTGAGCTT  GAGGGCTATCGAGCCAGTGATGGTCAGCATCTTGGTTCATTTGACCCTAAAACCGGTAAG  CAGTTGAAAGGTCCAGATCCGAAACGAAATATTAAAAAGTATCTTTAA  ATGGGACTTAAATTACACATTCATTGGTTTGATAAGAAGAATGAGGAATTTAAAGGTGGT  GAGTATTCAAAGGATTTCGGTGATGATGGTTCGGTTATTGAACGTCTTGGAATGCCTTTA  AAGGATAATATTAATAACGGTTGGTTTGATGTTGAAAAGAGATGGGTGCCAATATTACAG  CCATACTTTAAAAATGTTATTGATATTAGTAAGTTCGATTACTTTGTTTCTTTCGATTAT  CGTGATGGTGACTGGTAA | MSGGDGKGPGNTGLGHSGGKPSGNVNGGPTGLGGSGNSGGNNPNSGPGWGTTHTPNGDIHNYNPGEFGNG  GNKPGGNSGNHDGHGSGVQSAMAFGFPALAAPGAGTLGIAVSGEALSAAIADIFAALKGPFKFSAWGIAL  YGILPSEIAKDDPKMMSKIVTSLPAETVTNVQVSTLPLDQATVSVTKRVTDVVKDTRQHIAVVAGVPMSV  PVVNAKPTRTPGVFHASFPGVPSLTVSTVKGLPVSTTLPRGIMEDKGRTAVPAGFTFGGGSHEAVIRFPK  ESGQKPVYVSVTDVLTPAQVKQRQDEEKRLQQEWNDAHPVEVAERNYEQARAELNQANKDVARNQERQAK  AVQVYNSRKSELDAANKTLADAKAEIKQFERFAREPMAAGHRMWQMAGLKAQRAQTDVNNKKAAFDAAAK  EKSAADAALSSAMESRKKKEDKKRSAENKLNEEKNKRRKGTKDYGHDYHPAPKTEDIKGLGELKEGRPKT  PKQGGGGKRARWYGDKGRKIYEWDSQHGELEGYRASDGQHLGSFDPKTGKQLKGPDPKRNIKKYL  MGLKLHIHWFDKKNEEFKGGEYSKDFGDDGSVIERLGMPLKDNINNGWFDVEKRWVPILQPYFKNVIDIS  KFDYFVSFDYRDGDW | MSGGDGKGPGNTGLGHSGGKPSGNVNGGPTGLGGSGNSGGNNPNSGPGWGTTHTPNGDIHNYNPGEFGNG  GNKPGGNSGNHDGHGSGVQSAMAFGFPALAAPGAGTLGIAVSGEALSAAIADIFAALKGPFKFSAWGIAL  YGILPSEIAKDDPKMMSKIVTSLPAETVTNVQVSTLPLDQATVSVTKRVTDVVKDTRQHIAVVAGVPMSV  PVVNAKPTRTPGVFHASFPGVPSLTVSTVKGLPVSTTLPRGIMEDKGRTAVPAGFTFGGGSHEAVIRFPK  ESGQKPVYVSVTDVLTPAQVKQRQDEEKRLQQEWNDAHPVEVAERNYEQARAELNQANKDVARNQERQAK  AVQVYNSRKSELDAANKTLADAKAEIKQFERFAREPMAAGHRMWQMAGLKAQRAQTDVNNKKAAFDAAAK  EKSAADAALSSAMESRKKKEDKKRSAENKLNEEKNKRRKGTKDYGHDYHPAPKTEDIKGLGELKEGRPKT  PKQGGGGKRARWYGDKGRKIYEWDSQHGELEGYRASDGQHLGSFDPKTGKQLKGPDPKRNIKKYL  MGLKLHIHWFDKKNEEFKGGEYSKDFGDDGSVIERLGMPLKDNINNGWFDVEKRWVPILQPYFKNVIDIS  KFDYFVSFDYRDGDW  MKKITGIILLLLAAIILAACQANYIRDVQGGTVSPSSSAELTGVETQ |
| Plasmid ColE5-099 genes for colicin E5, immunity and lysis protein  <https://www.ncbi.nlm.nih.gov/nuccore/X15857.1?report=graph> | Colicin E5 |  |  |  |  |  |  |  |  |  |  |
| Plasmid ColE6-CT14 colicin E6, immunity proteins E6 and E8, and lysis protein genes, complete cds  <https://www.ncbi.nlm.nih.gov/nuccore/M31808.1?report=graph> | Colicin E6  (551 aa)  Immunity E6  (85 aa)  Immunity E8  (85 aa)  Lysis  (47 aa) | ATGAGCGGTGGCGATGGACGCGGCCATAACACGGGCGCGCATAGCACAAGTGGTAACATTAATGGTGGCC  CGACCGGGCTTGGTGTAGGTGGTGGTGCTTCTGATGGCTCCGGATGGAGTTCGGAAAATAACCCGTGGGG  TGGTGGTTCCGGTAGCGGCATTCACTGGGGTGGTGGTTCCGGTCATGGTAATGGCGGGGGGAATGGTAAT  TCCGGTGGTGGCTCGGGAACAGGCGGTAATCTGTCAGCAGTAGCTGCGCCAGTGGCATTTGGTTTTCCGG  CACTTTCCACTCCAGGAGCTGGCGGTCTGGCGGTCAGTATTTCAGCGGGAGCATTATCGGCAGCTATTGC  TGATATTATGGCTGCCCTGAAAGGACCGTTTAAATTTGGTCTTTGGGGGGTGGCTTTATATGGTGTATTG  CCATCACAAATAGCGAAAGATGACCCCAATATGATGTCAAAGATTGTGACGTCATTACCCGCAGATGATA  TTACTGAATCACCTGTCAGTTCATTACCTCTCGATAAGGCAACAGTAAACGTAAATGTTCGTGTTGTTGA  TGATGTAAAAGACGAACGACAGAATATTTCGGTTGTTTCAGGTGTTCCGATGAGTGTTCCGGTGGTTGAT  GCAAAACCTACCGAACGTCCAGGTGTTTTTACGGCATCAATTCCAGGTGCACCTGTTCTGAATATTTCAG  TTAATAACAGTACGCCAGCAGTACAGACATTAAGCCCAGGTGTTACAAATAATACTGATAAGGATGTTCG  CCCGGCAGGATTTACTCAGGGGGGTAATACCAGGGATGCAGTTATTCGATTCCCGAAGGACAGCGGTCAT  AATGCCGTATATGTTTCAGTGAGTGATGTTCTTAGCCCTGACCAGGTAAAACAACGTCAGGATGAAGAAA  ATCGCCGTCAGCAGGAATGGGATGCTACGCATCCGGTTGAAGCGGCTGAGCGAAATTATGAACGCGCGCG  TGCAGAGCTGAATCAGGCAAATGAAGATGTTGCCAGAAATCAGGAGCGACAGGCTAAAGCTGTTCAGGTT  TATAATTCGCGTAAAAGCGAACTTGATGCAGCGAATAAAACTCTTGCTGATGCAATAGCTGAAATAAAAC  AATTTAATCGATTTGCCCATGACCCAATGGCTGGCGGTCACAGAATGTGGCAAATGGCCGGGCTTAAAGC  CCAGCGGGCGCAGACGGATGTAAATAATAAGCAGGCTGCATTTGATGCTGCTGCAAAAGAGAAGTCAGAT  GCTGATGCTGCATTGAGTTCTGCTATGGAAAGCAGGAAGAAGAAAGAAGATAAGAAAAGGAGCGCTGAAA  ATAAATTAAACGAGGAAAAAAACAAGCCTCGCAAGGGAGTTAAAGATTACGGTCATGATTATCATCCAGA  TCCTAAAACTGAAGATATAAAAGGGCTGGGTGAGTTAAAAGAGGGTAAACCAAAAACTCCAAAGCAAGGT  GGTGGCGGTAAACGTGCTAGATGGTATGGAGATAAAGGGCGTAAGATTTATGAGTGGGACTCTCAGCATG  GTGAGCTTGAGGGGTATCGTGCCAGTGATGGTCAGCATCTTGGCTCATTCGAGCCTAAGACTGGTAATCA  GTTGAAAGGACCTGATCCAAAACGAAATATCAAAAAGTATCTTTGA | MSGGDGRGHNTGAHSTSGNINGGPTGLGVGGGASDGSGWSSENNPWGGGSGSGIHWGGGSGHGNGGGNGN  SGGGSGTGGNLSAVAAPVAFGFPALSTPGAGGLAVSISAGALSAAIADIMAALKGPFKFGLWGVALYGVL  PSQIAKDDPNMMSKIVTSLPADDITESPVSSLPLDKATVNVNVRVVDDVKDERQNISVVSGVPMSVPVVD  AKPTERPGVFTASIPGAPVLNISVNNSTPAVQTLSPGVTNNTDKDVRPAGFTQGGNTRDAVIRFPKDSGH  NAVYVSVSDVLSPDQVKQRQDEENRRQQEWDATHPVEAAERNYERARAELNQANEDVARNQERQAKAVQV  YNSRKSELDAANKTLADAIAEIKQFNRFAHDPMAGGHRMWQMAGLKAQRAQTDVNNKQAAFDAAAKEKSD  ADAALSSAMESRKKKEDKKRSAENKLNEEKNKPRKGVKDYGHDYHPDPKTEDIKGLGELKEGKPKTPKQG  GGGKRARWYGDKGRKIYEWDSQHGELEGYRASDGQHLGSFEPKTGNQLKGPDPKRNIKKYL | ATGGGGCTTAAATTACATATTAATTGGTTTGATAAGACGACCGAGGAATTTAAAGGTGGTGAGTATTCAA  AAGATTTTGGAGATGATGGCTCGGTCATTGAACGTCTTGGAATGCCTTTAAAAGATAATATCAATAATGG  TTGGTTTGATGTTATAGCTGAATGGGTACCTTTGCTACAACCATACTTTAATCATCAAATTGATATTTCC  GATAATGAGTATTTTGTTTCGTTTGATTATCGTGATGGTGATTGGTGA  GTGGAGCTAAAGAAAAGTATTGGTGATTACACTGAAACCGAATTCAAAAAAATTATTGAAAACATCATCA  ATTGTGAAGGTGATGAAAAAAAACAGGATGATAACCTCGAGCATTTTATAAGTGTTACTGAGCATCCTAG  TGGTTCTGATCTGATTTATTACCCAGAAGGTAATAATGATGGTAGCCCTGAAGCTGTTATTAAAGAGATT  AAAGAATGGCGAGCTGCTAACGGTAAGTCAGGATTTAAACAGGGCTGA | MGLKLHINWFDKTTEEFKGGEYSKDFGDDGSVIERLGMPLKDNINNGWFDVIAEWVPLLQPYFNHQIDIS  DNEYFVSFDYRDGDW  MELKKSIGDYTETEFKKIIENIINCEGDEKKQDDNLEHFISVTEHPSGSDLIYYPEGNNDGSPEAVIKEI  KEWRAANGKSGFKQG | ATGAAAAAAATAACAGGGATTATTTTATTGCTTCTTGCAGTCATTATTCTGGCTGCATGTCAGGCAAACT  ATATCCGTGATGTTCAGGGCGGGACTGTATCACCGTCGTCAACTGCTGAACTGACCGGAGTGGAAACGCA  GTAA | MKKITGIILLLLAVIILAACQANYIRDVQGGTVSPSSTAELTGVETQ | ATGAGCGGTGGCGATGGACGCGGCCATAACACGGGCGCGCATAGCACAAGTGGTAACATTAATGGTGGCC  CGACCGGGCTTGGTGTAGGTGGTGGTGCTTCTGATGGCTCCGGATGGAGTTCGGAAAATAACCCGTGGGG  TGGTGGTTCCGGTAGCGGCATTCACTGGGGTGGTGGTTCCGGTCATGGTAATGGCGGGGGGAATGGTAAT  TCCGGTGGTGGCTCGGGAACAGGCGGTAATCTGTCAGCAGTAGCTGCGCCAGTGGCATTTGGTTTTCCGG  CACTTTCCACTCCAGGAGCTGGCGGTCTGGCGGTCAGTATTTCAGCGGGAGCATTATCGGCAGCTATTGC  TGATATTATGGCTGCCCTGAAAGGACCGTTTAAATTTGGTCTTTGGGGGGTGGCTTTATATGGTGTATTG  CCATCACAAATAGCGAAAGATGACCCCAATATGATGTCAAAGATTGTGACGTCATTACCCGCAGATGATA  TTACTGAATCACCTGTCAGTTCATTACCTCTCGATAAGGCAACAGTAAACGTAAATGTTCGTGTTGTTGA  TGATGTAAAAGACGAACGACAGAATATTTCGGTTGTTTCAGGTGTTCCGATGAGTGTTCCGGTGGTTGAT  GCAAAACCTACCGAACGTCCAGGTGTTTTTACGGCATCAATTCCAGGTGCACCTGTTCTGAATATTTCAG  TTAATAACAGTACGCCAGCAGTACAGACATTAAGCCCAGGTGTTACAAATAATACTGATAAGGATGTTCG  CCCGGCAGGATTTACTCAGGGGGGTAATACCAGGGATGCAGTTATTCGATTCCCGAAGGACAGCGGTCAT  AATGCCGTATATGTTTCAGTGAGTGATGTTCTTAGCCCTGACCAGGTAAAACAACGTCAGGATGAAGAAA  ATCGCCGTCAGCAGGAATGGGATGCTACGCATCCGGTTGAAGCGGCTGAGCGAAATTATGAACGCGCGCG  TGCAGAGCTGAATCAGGCAAATGAAGATGTTGCCAGAAATCAGGAGCGACAGGCTAAAGCTGTTCAGGTT  TATAATTCGCGTAAAAGCGAACTTGATGCAGCGAATAAAACTCTTGCTGATGCAATAGCTGAAATAAAAC  AATTTAATCGATTTGCCCATGACCCAATGGCTGGCGGTCACAGAATGTGGCAAATGGCCGGGCTTAAAGC  CCAGCGGGCGCAGACGGATGTAAATAATAAGCAGGCTGCATTTGATGCTGCTGCAAAAGAGAAGTCAGAT  GCTGATGCTGCATTGAGTTCTGCTATGGAAAGCAGGAAGAAGAAAGAAGATAAGAAAAGGAGCGCTGAAA  ATAAATTAAACGAGGAAAAAAACAAGCCTCGCAAGGGAGTTAAAGATTACGGTCATGATTATCATCCAGA  TCCTAAAACTGAAGATATAAAAGGGCTGGGTGAGTTAAAAGAGGGTAAACCAAAAACTCCAAAGCAAGGT  GGTGGCGGTAAACGTGCTAGATGGTATGGAGATAAAGGGCGTAAGATTTATGAGTGGGACTCTCAGCATG  GTGAGCTTGAGGGGTATCGTGCCAGTGATGGTCAGCATCTTGGCTCATTCGAGCCTAAGACTGGTAATCA  GTTGAAAGGACCTGATCCAAAACGAAATATCAAAAAGTATCTTTGAGAGGATGTTATGGGGCTTAAATTA  CATATTAATTGGTTTGATAAGACGACCGAGGAATTTAAAGGTGGTGAGTATTCAAAAGATTTTGGAGATG  ATGGCTCGGTCATTGAACGTCTTGGAATGCCTTTAAAAGATAATATCAATAATGGTTGGTTTGATGTTAT  AGCTGAATGGGTACCTTTGCTACAACCATACTTTAATCATCAAATTGATATTTCCGATAATGAGTATTTT  GTTTCGTTTGATTATCGTGATGGTGATTGGTGATCAAATATTATCAGGGATGAGTTGATGTACGGGCTTC  TAGTGTTCATGGATGAACGCTGGAGCCTCCAAATGTAGAAGTGTTATATTTTTTATTGAGTTCTTGGTTA  TAATTGCTCCGCAATAATTTAAATAGGCATTATTTAAAACATTCTCAGGAGAGGTGAAGGTGGAGCTAAA  GAAAAGTATTGGTGATTACACTGAAACCGAATTCAAAAAAATTATTGAAAACATCATCAATTGTGAAGGT  GATGAAAAAAAACAGGATGATAACCTCGAGCATTTTATAAGTGTTACTGAGCATCCTAGTGGTTCTGATC  TGATTTATTACCCAGAAGGTAATAATGATGGTAGCCCTGAAGCTGTTATTAAAGAGATTAAAGAATGGCG  AGCTGCTAACGGTAAGTCAGGATTTAAACAGGGCTGAAATATGAATGCCGGTTGTTTAAGGATGAATGAC  TGGCATTCTTTCACAACAAGGAGTCGTTATGAAAAAAATAACAGGGATTATTTTATTGCTTCTTGCAGTC  ATTATTCTGGCTGCATGTCAGGCAAACTATATCCGTGATGTTCAGGGCGGGACTGTATCACCGTCGTCAA  CTGCTGAACTGACCGGAGTGGAAACGCAGTAA | ATGAGCGGTGGCGATGGACGCGGCCATAACACGGGCGCGCATAGCACAAGTGGTAACATTAATGGTGGCC  CGACCGGGCTTGGTGTAGGTGGTGGTGCTTCTGATGGCTCCGGATGGAGTTCGGAAAATAACCCGTGGGG  TGGTGGTTCCGGTAGCGGCATTCACTGGGGTGGTGGTTCCGGTCATGGTAATGGCGGGGGGAATGGTAAT  TCCGGTGGTGGCTCGGGAACAGGCGGTAATCTGTCAGCAGTAGCTGCGCCAGTGGCATTTGGTTTTCCGG  CACTTTCCACTCCAGGAGCTGGCGGTCTGGCGGTCAGTATTTCAGCGGGAGCATTATCGGCAGCTATTGC  TGATATTATGGCTGCCCTGAAAGGACCGTTTAAATTTGGTCTTTGGGGGGTGGCTTTATATGGTGTATTG  CCATCACAAATAGCGAAAGATGACCCCAATATGATGTCAAAGATTGTGACGTCATTACCCGCAGATGATA  TTACTGAATCACCTGTCAGTTCATTACCTCTCGATAAGGCAACAGTAAACGTAAATGTTCGTGTTGTTGA  TGATGTAAAAGACGAACGACAGAATATTTCGGTTGTTTCAGGTGTTCCGATGAGTGTTCCGGTGGTTGAT  GCAAAACCTACCGAACGTCCAGGTGTTTTTACGGCATCAATTCCAGGTGCACCTGTTCTGAATATTTCAG  TTAATAACAGTACGCCAGCAGTACAGACATTAAGCCCAGGTGTTACAAATAATACTGATAAGGATGTTCG  CCCGGCAGGATTTACTCAGGGGGGTAATACCAGGGATGCAGTTATTCGATTCCCGAAGGACAGCGGTCAT  AATGCCGTATATGTTTCAGTGAGTGATGTTCTTAGCCCTGACCAGGTAAAACAACGTCAGGATGAAGAAA  ATCGCCGTCAGCAGGAATGGGATGCTACGCATCCGGTTGAAGCGGCTGAGCGAAATTATGAACGCGCGCG  TGCAGAGCTGAATCAGGCAAATGAAGATGTTGCCAGAAATCAGGAGCGACAGGCTAAAGCTGTTCAGGTT  TATAATTCGCGTAAAAGCGAACTTGATGCAGCGAATAAAACTCTTGCTGATGCAATAGCTGAAATAAAAC  AATTTAATCGATTTGCCCATGACCCAATGGCTGGCGGTCACAGAATGTGGCAAATGGCCGGGCTTAAAGC  CCAGCGGGCGCAGACGGATGTAAATAATAAGCAGGCTGCATTTGATGCTGCTGCAAAAGAGAAGTCAGAT  GCTGATGCTGCATTGAGTTCTGCTATGGAAAGCAGGAAGAAGAAAGAAGATAAGAAAAGGAGCGCTGAAA  ATAAATTAAACGAGGAAAAAAACAAGCCTCGCAAGGGAGTTAAAGATTACGGTCATGATTATCATCCAGA  TCCTAAAACTGAAGATATAAAAGGGCTGGGTGAGTTAAAAGAGGGTAAACCAAAAACTCCAAAGCAAGGT  GGTGGCGGTAAACGTGCTAGATGGTATGGAGATAAAGGGCGTAAGATTTATGAGTGGGACTCTCAGCATG  GTGAGCTTGAGGGGTATCGTGCCAGTGATGGTCAGCATCTTGGCTCATTCGAGCCTAAGACTGGTAATCA  GTTGAAAGGACCTGATCCAAAACGAAATATCAAAAAGTATCTTTGA  ATGGGGCTTAAATTACATATTAATTGGTTTGATAAGACGACCGAGGAATTTAAAGGTGGTGAGTATTCAA  AAGATTTTGGAGATGATGGCTCGGTCATTGAACGTCTTGGAATGCCTTTAAAAGATAATATCAATAATGG  TTGGTTTGATGTTATAGCTGAATGGGTACCTTTGCTACAACCATACTTTAATCATCAAATTGATATTTCC  GATAATGAGTATTTTGTTTCGTTTGATTATCGTGATGGTGATTGGTGA  GTGGAGCTAAAGAAAAGTATTGGTGATTACACTGAAACCGAATTCAAAAAAATTATTGAAAACATCATCA  ATTGTGAAGGTGATGAAAAAAAACAGGATGATAACCTCGAGCATTTTATAAGTGTTACTGAGCATCCTAG  TGGTTCTGATCTGATTTATTACCCAGAAGGTAATAATGATGGTAGCCCTGAAGCTGTTATTAAAGAGATT  AAAGAATGGCGAGCTGCTAACGGTAAGTCAGGATTTAAACAGGGCTGA | MSGGDGRGHNTGAHSTSGNINGGPTGLGVGGGASDGSGWSSENNPWGGGSGSGIHWGGGSGHGNGGGNGN  SGGGSGTGGNLSAVAAPVAFGFPALSTPGAGGLAVSISAGALSAAIADIMAALKGPFKFGLWGVALYGVL  PSQIAKDDPNMMSKIVTSLPADDITESPVSSLPLDKATVNVNVRVVDDVKDERQNISVVSGVPMSVPVVD  AKPTERPGVFTASIPGAPVLNISVNNSTPAVQTLSPGVTNNTDKDVRPAGFTQGGNTRDAVIRFPKDSGH  NAVYVSVSDVLSPDQVKQRQDEENRRQQEWDATHPVEAAERNYERARAELNQANEDVARNQERQAKAVQV  YNSRKSELDAANKTLADAIAEIKQFNRFAHDPMAGGHRMWQMAGLKAQRAQTDVNNKQAAFDAAAKEKSD  ADAALSSAMESRKKKEDKKRSAENKLNEEKNKPRKGVKDYGHDYHPDPKTEDIKGLGELKEGKPKTPKQG  GGGKRARWYGDKGRKIYEWDSQHGELEGYRASDGQHLGSFEPKTGNQLKGPDPKRNIKKYL  MGLKLHINWFDKTTEEFKGGEYSKDFGDDGSVIERLGMPLKDNINNGWFDVIAEWVPLLQPYFNHQIDIS  DNEYFVSFDYRDGDW  MELKKSIGDYTETEFKKIIENIINCEGDEKKQDDNLEHFISVTEHPSGSDLIYYPEGNNDGSPEAVIKEI  KEWRAANGKSGFKQG | MSGGDGRGHNTGAHSTSGNINGGPTGLGVGGGASDGSGWSSENNPWGGGSGSGIHWGGGSGHGNGGGNGN  SGGGSGTGGNLSAVAAPVAFGFPALSTPGAGGLAVSISAGALSAAIADIMAALKGPFKFGLWGVALYGVL  PSQIAKDDPNMMSKIVTSLPADDITESPVSSLPLDKATVNVNVRVVDDVKDERQNISVVSGVPMSVPVVD  AKPTERPGVFTASIPGAPVLNISVNNSTPAVQTLSPGVTNNTDKDVRPAGFTQGGNTRDAVIRFPKDSGH  NAVYVSVSDVLSPDQVKQRQDEENRRQQEWDATHPVEAAERNYERARAELNQANEDVARNQERQAKAVQV  YNSRKSELDAANKTLADAIAEIKQFNRFAHDPMAGGHRMWQMAGLKAQRAQTDVNNKQAAFDAAAKEKSD  ADAALSSAMESRKKKEDKKRSAENKLNEEKNKPRKGVKDYGHDYHPDPKTEDIKGLGELKEGKPKTPKQG  GGGKRARWYGDKGRKIYEWDSQHGELEGYRASDGQHLGSFEPKTGNQLKGPDPKRNIKKYL  MGLKLHINWFDKTTEEFKGGEYSKDFGDDGSVIERLGMPLKDNINNGWFDVIAEWVPLLQPYFNHQIDIS  DNEYFVSFDYRDGDW  MELKKSIGDYTETEFKKIIENIINCEGDEKKQDDNLEHFISVTEHPSGSDLIYYPEGNNDGSPEAVIKEI  KEWRAANGKSGFKQG  MKKITGIILLLLAVIILAACQANYIRDVQGGTVSPSSTAELTGVETQ |
| Escherichia coli strain Ecol_AZ146 plasmid pECAZ146_5, complete sequence  [https://www.ncbi.nlm.nih.gov/nuccore/CP018986.1?report=graph&rid=EEK75R9G016[CP018986.1]&tracks=[key:sequence_track,name:Sequence,display_name:Sequence,id:STD1,category:Sequence,annots:Sequence,ShowLabel:true][key:gene_model_track,CDSProductFeats:false][key:alignment_track,name:other%20alignments,annots:NG%20Alignments\|Refseq%20Alignments\|Gnomon%20Alignments\|Unnamed,shown:false]&v=2196:2747&appname=ncbiblast&link_loc=fromHSP](https://www.ncbi.nlm.nih.gov/nuccore/CP018986.1?report=graph&rid=EEK75R9G016%5bCP018986.1%5d&tracks=%5bkey:sequence_track,name:Sequence,display_name:Sequence,id:STD1,category:Sequence,annots:Sequence,ShowLabel:true%5d%5bkey:gene_model_track,CDSProductFeats:false%5d%5bkey:alignment_track,name:other%20alignments,annots:NG%20Alignments\|Refseq%20Alignments\|Gnomon%20Alignments\|Unnamed,shown:false%5d&v=2196:2747&appname=ncbiblast&link_loc=fromHSP) | Colicin E7  (576 aa)  Immunity  (87 aa)  Lysis  (47 aa) | ATGAGCGGTGGAGATGGACGTGGCCACAACAGTGGCGCACATAACACAGGTGGTAACATT  AATGGCGGCCCTACGGGGCTTGGTGGAAATGGTGGGGCTTCTGACGGCTCCGGATGGAGT  TCGGAAAATAACCCATGGGGTGGCGGTTCCGGTAGTGGTGTTCACTGGGGAGGTGGCTCC  GGCCATGGCAATGGCGGGGGGAATAGTAATTCCGGTGGTGGCAGCAATTCATCCGTAGCA  GCACCTATGGCATTTGGTTTTCCTGCTTTGGCTGCTCCTGGTGCCGGAACTCTTGGTATT  TCCGTATCTGGTGAGGCTTTATCTGCGGCAATAGCGGATATCTTTGCAGCCCTGAAAGGT  CCGTTTAAATTCAGCGCATGGGGTATTGCGCTTTACGGCATTCTGCCATCTGAAATAGCA  AAAGATGACCCGAATATGATGTCAAAGATTGTGACGTCATTACCGGCAGAAACAGTGACG  AATGTTCAGGTCAGTACACTTCCGCTGGACCAGGCAACTGTCAGCGTGACGAAACGTGTG  ACAGATGTTGTGAAGGACACACGACAGCATATTGCCGTAGTTGCAGGTGTGCCTATGAGT  GTGCCTGTTGTGAATGCCAAACCAACACGTACTCCAGGTGTATTTCACGCATCATTTCCT  GGTGTCCCTTCTCTGACAGTCAGCACTGTCAAAGGCCTTCCGGTGTCAACAACCCTTCCC  CGTGGTATTACGGAGGATAAAGGCCGGACTGCCGTTCCTGCGGGATTTACCTTTGGTGGT  GGTTCACATGAAGCGGTGATTCGCTTTCCGAAAGAAAGTGGGCAGAAGCCGGTTTATGTA  TCAGTGACAGATGTTCTTACCCCAGCACAGGTAAAACAACGTCAGGATGAGGAAAAACGC  CTCCAGCAGGAATGGAATGACGCACATCCGGTGGAAGTGGCTGAACGCAATTATGAACAG  GCCCGCGCAGAACTGAATCAGGCTAATAAAGATGTTGCCAGAAACCAGGAACGACAGGCT  AAAGCTGTTCAGGTTTATAATTCTCGTAAAAGTGAACTTGATGCAGCGAATAAAACTCTT  GCTGATGCAAAGGCTGAAATAAAACAATTCGAGCGATTTGCCCGAGAACCAATGGCTGCT  GGTCACAGAATGTGGCAAATGGCAGGGCTTAAGGCCCAGCGGGCACAGACGGATGTAAAT  AATAAGAAGGCTGCATTTGATGCTGCTGCAAAAGAAAAGTCAGATGCTGATGTTGCATTG  AGTTCTGCGTTGGAGCGCCGCAAACAGAAAGAAAATAAAGAAAAGGACGCTAAAGCCAAA  TTAGATAAGGAGAGTAAACGGAATAAGCCAGGGAAGGCAACAGGTAAAGGAAAACCTGTC  AATAATAAGTGGTTAAATAATGCAGGTAAAGACTTAGGTTCTCCTGTTCCAGATCGTATA  GCTAATAAACTACGTGATAAGGAGTTTAAAAGTTTCGATGATTTTCGTAAGAAATTCTGG  GAAGAAGTGTCAAAAGATCCTGAGTTAAGTAAACAATTTAGTAGGAACAATAATGATCGA  ATGAAGGTTGGAAAAGCGCCCAAGACTAGAACCCAGGATGTTTCAGGGAAGAGAACTTCA  TTCGAGCTTCATCATGAGAAGCCGATCAGCCAAAATGGTGGTGTCTATGATATGGATAAC  ATCAGCGTGGTAACACCTAAAAGGCATATTGATATTCACCGAGGTAAATAA | MSGGDGRGHNSGAHNTGGNINGGPTGLGGNGGASDGSGWSSENNPWGGGSGSGVHWGGGSGHGNGGGNSN  SGGGSNSSVAAPMAFGFPALAAPGAGTLGISVSGEALSAAIADIFAALKGPFKFSAWGIALYGILPSEIA  KDDPNMMSKIVTSLPAETVTNVQVSTLPLDQATVSVTKRVTDVVKDTRQHIAVVAGVPMSVPVVNAKPTR  TPGVFHASFPGVPSLTVSTVKGLPVSTTLPRGITEDKGRTAVPAGFTFGGGSHEAVIRFPKESGQKPVYV  SVTDVLTPAQVKQRQDEEKRLQQEWNDAHPVEVAERNYEQARAELNQANKDVARNQERQAKAVQVYNSRK  SELDAANKTLADAKAEIKQFERFAREPMAAGHRMWQMAGLKAQRAQTDVNNKKAAFDAAAKEKSDADVAL  SSALERRKQKENKEKDAKAKLDKESKRNKPGKATGKGKPVNNKWLNNAGKDLGSPVPDRIANKLRDKEFK  SFDDFRKKFWEEVSKDPELSKQFSRNNNDRMKVGKAPKTRTQDVSGKRTSFELHHEKPISQNGGVYDMDN  ISVVTPKRHIDIHRGK | ATGGAACTGAAAAATAGTATTAGTGATTACACAGAGGCTGAGTTTGTTCAACTTCTTAAG  GAAATTGAAAAAGAGAATGTTGCTGCAACTGATGATGTGTTAGATGTGTTACTCGAACAC  TTTGTAAAAATTACTGAGCATCCAGATGGAACGGATCTGATTTATTATCCTAGTGATAAT  AGAGACGATAGCCCCGAAGGGATTGTCAAGGAAATTAAAGAATGGCGAGCTGCTAACGGT  AAGCCAGGATTTAAACAGGGCTAA | MELKNSISDYTEAEFVQLLKEIEKENVAATDDVLDVLLEHFVKITEHPDGTDLIYYPSDNRDDSPEGIVK  EIKEWRAANGKPGFKQG | ATGAAAAAAATAACAGGGATTATTTTATTGCTTCTTGCAGCCATTATTCTTGCTGCATGT  CAGGCAAACTATATCCGTGATGTTCAGGGCGGGACAGTATCACCGTCGTCAACTGCTGAA  CTGACCGGAGTGGAAACGCAGTAA | MKKITGIILLLLAAIILAACQANYIRDVQGGTVSPSSTAELTGVETQ | ATGAGCGGTGGAGATGGACGTGGCCACAACAGTGGCGCACATAACACAGGTGGTAACATT  AATGGCGGCCCTACGGGGCTTGGTGGAAATGGTGGGGCTTCTGACGGCTCCGGATGGAGT  TCGGAAAATAACCCATGGGGTGGCGGTTCCGGTAGTGGTGTTCACTGGGGAGGTGGCTCC  GGCCATGGCAATGGCGGGGGGAATAGTAATTCCGGTGGTGGCAGCAATTCATCCGTAGCA  GCACCTATGGCATTTGGTTTTCCTGCTTTGGCTGCTCCTGGTGCCGGAACTCTTGGTATT  TCCGTATCTGGTGAGGCTTTATCTGCGGCAATAGCGGATATCTTTGCAGCCCTGAAAGGT  CCGTTTAAATTCAGCGCATGGGGTATTGCGCTTTACGGCATTCTGCCATCTGAAATAGCA  AAAGATGACCCGAATATGATGTCAAAGATTGTGACGTCATTACCGGCAGAAACAGTGACG  AATGTTCAGGTCAGTACACTTCCGCTGGACCAGGCAACTGTCAGCGTGACGAAACGTGTG  ACAGATGTTGTGAAGGACACACGACAGCATATTGCCGTAGTTGCAGGTGTGCCTATGAGT  GTGCCTGTTGTGAATGCCAAACCAACACGTACTCCAGGTGTATTTCACGCATCATTTCCT  GGTGTCCCTTCTCTGACAGTCAGCACTGTCAAAGGCCTTCCGGTGTCAACAACCCTTCCC  CGTGGTATTACGGAGGATAAAGGCCGGACTGCCGTTCCTGCGGGATTTACCTTTGGTGGT  GGTTCACATGAAGCGGTGATTCGCTTTCCGAAAGAAAGTGGGCAGAAGCCGGTTTATGTA  TCAGTGACAGATGTTCTTACCCCAGCACAGGTAAAACAACGTCAGGATGAGGAAAAACGC  CTCCAGCAGGAATGGAATGACGCACATCCGGTGGAAGTGGCTGAACGCAATTATGAACAG  GCCCGCGCAGAACTGAATCAGGCTAATAAAGATGTTGCCAGAAACCAGGAACGACAGGCT  AAAGCTGTTCAGGTTTATAATTCTCGTAAAAGTGAACTTGATGCAGCGAATAAAACTCTT  GCTGATGCAAAGGCTGAAATAAAACAATTCGAGCGATTTGCCCGAGAACCAATGGCTGCT  GGTCACAGAATGTGGCAAATGGCAGGGCTTAAGGCCCAGCGGGCACAGACGGATGTAAAT  AATAAGAAGGCTGCATTTGATGCTGCTGCAAAAGAAAAGTCAGATGCTGATGTTGCATTG  AGTTCTGCGTTGGAGCGCCGCAAACAGAAAGAAAATAAAGAAAAGGACGCTAAAGCCAAA  TTAGATAAGGAGAGTAAACGGAATAAGCCAGGGAAGGCAACAGGTAAAGGAAAACCTGTC  AATAATAAGTGGTTAAATAATGCAGGTAAAGACTTAGGTTCTCCTGTTCCAGATCGTATA  GCTAATAAACTACGTGATAAGGAGTTTAAAAGTTTCGATGATTTTCGTAAGAAATTCTGG  GAAGAAGTGTCAAAAGATCCTGAGTTAAGTAAACAATTTAGTAGGAACAATAATGATCGA  ATGAAGGTTGGAAAAGCGCCCAAGACTAGAACCCAGGATGTTTCAGGGAAGAGAACTTCA  TTCGAGCTTCATCATGAGAAGCCGATCAGCCAAAATGGTGGTGTCTATGATATGGATAAC  ATCAGCGTGGTAACACCTAAAAGGCATATTGATATTCACCGAGGTAAATAATATGGAACT  GAAAAATAGTATTAGTGATTACACAGAGGCTGAGTTTGTTCAACTTCTTAAGGAAATTGA  AAAAGAGAATGTTGCTGCAACTGATGATGTGTTAGATGTGTTACTCGAACACTTTGTAAA  AATTACTGAGCATCCAGATGGAACGGATCTGATTTATTATCCTAGTGATAATAGAGACGA  TAGCCCCGAAGGGATTGTCAAGGAAATTAAAGAATGGCGAGCTGCTAACGGTAAGCCAGG  ATTTAAACAGGGCTAAAATATGAATGCCGGTTGTTTAAGGATGAATGACTGGCATTCTTT  CACAACAAGGAGTCGTTATGAAAAAAATAACAGGGATTATTTTATTGCTTCTTGCAGCCA  TTATTCTTGCTGCATGTCAGGCAAACTATATCCGTGATGTTCAGGGCGGGACAGTATCAC  CGTCGTCAACTGCTGAACTGACCGGAGTGGAAACGCAGTAA | ATGAGCGGTGGAGATGGACGTGGCCACAACAGTGGCGCACATAACACAGGTGGTAACATT  AATGGCGGCCCTACGGGGCTTGGTGGAAATGGTGGGGCTTCTGACGGCTCCGGATGGAGT  TCGGAAAATAACCCATGGGGTGGCGGTTCCGGTAGTGGTGTTCACTGGGGAGGTGGCTCC  GGCCATGGCAATGGCGGGGGGAATAGTAATTCCGGTGGTGGCAGCAATTCATCCGTAGCA  GCACCTATGGCATTTGGTTTTCCTGCTTTGGCTGCTCCTGGTGCCGGAACTCTTGGTATT  TCCGTATCTGGTGAGGCTTTATCTGCGGCAATAGCGGATATCTTTGCAGCCCTGAAAGGT  CCGTTTAAATTCAGCGCATGGGGTATTGCGCTTTACGGCATTCTGCCATCTGAAATAGCA  AAAGATGACCCGAATATGATGTCAAAGATTGTGACGTCATTACCGGCAGAAACAGTGACG  AATGTTCAGGTCAGTACACTTCCGCTGGACCAGGCAACTGTCAGCGTGACGAAACGTGTG  ACAGATGTTGTGAAGGACACACGACAGCATATTGCCGTAGTTGCAGGTGTGCCTATGAGT  GTGCCTGTTGTGAATGCCAAACCAACACGTACTCCAGGTGTATTTCACGCATCATTTCCT  GGTGTCCCTTCTCTGACAGTCAGCACTGTCAAAGGCCTTCCGGTGTCAACAACCCTTCCC  CGTGGTATTACGGAGGATAAAGGCCGGACTGCCGTTCCTGCGGGATTTACCTTTGGTGGT  GGTTCACATGAAGCGGTGATTCGCTTTCCGAAAGAAAGTGGGCAGAAGCCGGTTTATGTA  TCAGTGACAGATGTTCTTACCCCAGCACAGGTAAAACAACGTCAGGATGAGGAAAAACGC  CTCCAGCAGGAATGGAATGACGCACATCCGGTGGAAGTGGCTGAACGCAATTATGAACAG  GCCCGCGCAGAACTGAATCAGGCTAATAAAGATGTTGCCAGAAACCAGGAACGACAGGCT  AAAGCTGTTCAGGTTTATAATTCTCGTAAAAGTGAACTTGATGCAGCGAATAAAACTCTT  GCTGATGCAAAGGCTGAAATAAAACAATTCGAGCGATTTGCCCGAGAACCAATGGCTGCT  GGTCACAGAATGTGGCAAATGGCAGGGCTTAAGGCCCAGCGGGCACAGACGGATGTAAAT  AATAAGAAGGCTGCATTTGATGCTGCTGCAAAAGAAAAGTCAGATGCTGATGTTGCATTG  AGTTCTGCGTTGGAGCGCCGCAAACAGAAAGAAAATAAAGAAAAGGACGCTAAAGCCAAA  TTAGATAAGGAGAGTAAACGGAATAAGCCAGGGAAGGCAACAGGTAAAGGAAAACCTGTC  AATAATAAGTGGTTAAATAATGCAGGTAAAGACTTAGGTTCTCCTGTTCCAGATCGTATA  GCTAATAAACTACGTGATAAGGAGTTTAAAAGTTTCGATGATTTTCGTAAGAAATTCTGG  GAAGAAGTGTCAAAAGATCCTGAGTTAAGTAAACAATTTAGTAGGAACAATAATGATCGA  ATGAAGGTTGGAAAAGCGCCCAAGACTAGAACCCAGGATGTTTCAGGGAAGAGAACTTCA  TTCGAGCTTCATCATGAGAAGCCGATCAGCCAAAATGGTGGTGTCTATGATATGGATAAC  ATCAGCGTGGTAACACCTAAAAGGCATATTGATATTCACCGAGGTAAATAA  ATGGAACTGAAAAATAGTATTAGTGATTACACAGAGGCTGAGTTTGTTCAACTTCTTAAG  GAAATTGAAAAAGAGAATGTTGCTGCAACTGATGATGTGTTAGATGTGTTACTCGAACAC  TTTGTAAAAATTACTGAGCATCCAGATGGAACGGATCTGATTTATTATCCTAGTGATAAT  AGAGACGATAGCCCCGAAGGGATTGTCAAGGAAATTAAAGAATGGCGAGCTGCTAACGGT  AAGCCAGGATTTAAACAGGGCTAA | MSGGDGRGHNSGAHNTGGNINGGPTGLGGNGGASDGSGWSSENNPWGGGSGSGVHWGGGSGHGNGGGNSN  SGGGSNSSVAAPMAFGFPALAAPGAGTLGISVSGEALSAAIADIFAALKGPFKFSAWGIALYGILPSEIA  KDDPNMMSKIVTSLPAETVTNVQVSTLPLDQATVSVTKRVTDVVKDTRQHIAVVAGVPMSVPVVNAKPTR  TPGVFHASFPGVPSLTVSTVKGLPVSTTLPRGITEDKGRTAVPAGFTFGGGSHEAVIRFPKESGQKPVYV  SVTDVLTPAQVKQRQDEEKRLQQEWNDAHPVEVAERNYEQARAELNQANKDVARNQERQAKAVQVYNSRK  SELDAANKTLADAKAEIKQFERFAREPMAAGHRMWQMAGLKAQRAQTDVNNKKAAFDAAAKEKSDADVAL  SSALERRKQKENKEKDAKAKLDKESKRNKPGKATGKGKPVNNKWLNNAGKDLGSPVPDRIANKLRDKEFK  SFDDFRKKFWEEVSKDPELSKQFSRNNNDRMKVGKAPKTRTQDVSGKRTSFELHHEKPISQNGGVYDMDN  ISVVTPKRHIDIHRGK  MELKNSISDYTEAEFVQLLKEIEKENVAATDDVLDVLLEHFVKITEHPDGTDLIYYPSDNRDDSPEGIVK  EIKEWRAANGKPGFKQG | MSGGDGRGHNSGAHNTGGNINGGPTGLGGNGGASDGSGWSSENNPWGGGSGSGVHWGGGSGHGNGGGNSN  SGGGSNSSVAAPMAFGFPALAAPGAGTLGISVSGEALSAAIADIFAALKGPFKFSAWGIALYGILPSEIA  KDDPNMMSKIVTSLPAETVTNVQVSTLPLDQATVSVTKRVTDVVKDTRQHIAVVAGVPMSVPVVNAKPTR  TPGVFHASFPGVPSLTVSTVKGLPVSTTLPRGITEDKGRTAVPAGFTFGGGSHEAVIRFPKESGQKPVYV  SVTDVLTPAQVKQRQDEEKRLQQEWNDAHPVEVAERNYEQARAELNQANKDVARNQERQAKAVQVYNSRK  SELDAANKTLADAKAEIKQFERFAREPMAAGHRMWQMAGLKAQRAQTDVNNKKAAFDAAAKEKSDADVAL  SSALERRKQKENKEKDAKAKLDKESKRNKPGKATGKGKPVNNKWLNNAGKDLGSPVPDRIANKLRDKEFK  SFDDFRKKFWEEVSKDPELSKQFSRNNNDRMKVGKAPKTRTQDVSGKRTSFELHHEKPISQNGGVYDMDN  ISVVTPKRHIDIHRGK  MELKNSISDYTEAEFVQLLKEIEKENVAATDDVLDVLLEHFVKITEHPDGTDLIYYPSDNRDDSPEGIVK  EIKEWRAANGKPGFKQG  MKKITGIILLLLAAIILAACQANYIRDVQGGTVSPSSTAELTGVETQ |
| Escherichia coli plasmid pColE8, complete sequence  <https://www.ncbi.nlm.nih.gov/nuccore/FJ985252.1?report=graph> | colicin E8  *ce8a*  573 aa  immunity E8  *ce8i*  85 aa  lysis E8  *ce8l*  47 aa | ATGAGCGGTGGAGATGGACGCGGCCATAACACGGGCGCGCATAGCACAAGTGGTAACATTAATGGTGGCCCGACCGGGATTGGTGTAAGTGGTGGTGCTTCTGATGGTTCAGGATGGAGTTCGGAAAATAACCCGTGGGGTGGTGGTTCCGGTAGCGGCATTCACTGGGGAGGTGGCTCCGGTCGTGGTAATGGCGGGGGTAATGGCAATTCCGGTGGTGGCTCGGGAACAGGCGGTAATTTGTCAGCAGTAGCTGCGCCAGTGGCATTTGGTTTTCCGGCTCTTTCCACTCCAGGAGCTGGCGGTCTGGCTGTCAGTATTTCTGCAAGCGAATTATCGGCAGCTATTGCTGGTATTATTGCTAAATTAAAAAAAGTAAATCTTAAATTCACTCCTTTTGGGGTTGTCTTATCTTCATTAATTCCGTCGGAAATAGCGAAAGATGACCCCAATATGATGTCAAAGATTGTGACGTCATTACCCGCAGATGATATTACTGAATCACCTGTCAGTTCATTACCTCTCGATAAGGCAACAGTAAACGTAAATGTTCGTGTTGTTGATGATGTAAAAGACGAACGACAGAATATTTCGGTTGTTTCAGGTGTTCCGATGAGTGTTCCGGTGGTTGATGCAAAACCTACCGAACGTCCAGGTGTTTTTACGGCATCAATTCCAGGTGCACCTGTTCTGAATATTTCAGTTAATAACAGTACGCCAGCAGTACAGACATTAAGCCCAGGTGTTACAAATAATACTGATAAGGATGTTCGCCCGGCAGGATTTACTCAGGGTGGTAATACCAGGGATGCAGTTATTCGATTCCCGAAGGACAGCGGTCATAATGCCGTATATGTTTCAGTGAGTGATGTTCTTAGTCCTGACCAGGTAAAACAACGTCAGGATGAAGAAAATCGCCGTCAGCAGGAATGGGATGCTACGCATCCGGTTGAAGCGGCTGAGCGAAATTATGAACGCGCGCGTGCAGAGCTGAATCAGGCAAATGAAGATGTTGCCAGAAATCAGGAGCGACAGGCTAAAGCTGTTCAGGTTTATAATTCGCGTAAAAGCGAACTTGATGCAGCGAATAAGACTCTTGCTGATGCAATAGCTGAAATAAAACAATTTAATCGATTTGCCCATGACCCAATGGCTGGCGGTCACAGAATGTGGCAAATGGCCGGGCTTAAAGCTCAGCGGGCGCAGACGGATGTAAATAATAAGCAGGCTGCAGATGCTGATGCTGCATTAAGTGCCGCGCAGGAGCGCCGCAAACAGAAGGAAAATAAAGAGAAGGACGCTAAGGATAAATTAGATAAGGAGAGTAAACGGAATAAGCCAGGGAAGGCGACAGGTAAAGGTAAACCAGTTGGTGATAAATGGCTGGATGATGCGGGTAAAGATTCAGGAGCGCCAATTCCAGATCGCATTGCTGATAAGTTGCGTGATAAAGAATTTAAAAACTTCGACGATTTTCGGAGAAAATTCTGGGAAGAAGTGTCGAAAGATCCTGAGTTAAGTAAGCAATTTAATCCAGGTAATAAAAAACGCTTGAGCCAAGGATTAGCTCCACGGGCTAGGAATAAAGATACTGTCGGTGGGCGTCGCTCATTTGAGCTTCATCACGACAAGCCAATCAGTCAGGATGTGGTGTTTATGATATGGACAACCTCCGTATCACTACGCCGAAACGCCATATTGATATTCACAGAGGTCAGTAA | MSGGDGRGHNTGAHSTSGNINGGPTGIGVSGGASDGSGWSSENNPWGGGSGSGIHWGGGSGRGNGGGNGNSGGGSGTGGNLSAVAAPVAFGFPALSTPGAGGLAVSISASELSAAIAGIIAKLKKVNLKFTPFGVVLSSLIPSEIAKDDPNMMSKIVTSLPADDITESPVSSLPLDKATVNVNVRVVDDVKDERQNISVVSGVPMSVPVVDAKPTERPGVFTASIPGAPVLNISVNNSTPAVQTLSPGVTNNTDKDVRPAGFTQGGNTRDAVIRFPKDSGHNAVYVSVSDVLSPDQVKQRQDEENRRQQEWDATHPVEAAERNYERARAELNQANEDVARNQERQAKAVQVYNSRKSELDAANKTLADAIAEIKQFNRFAHDPMAGGHRMWQMAGLKAQRAQTDVNNKQAADADAALSAAQERRKQKENKEKDAKDKLDKESKRNKPGKATGKGKPVGDKWLDDAGKDSGAPIPDRIADKLRDKEFKNFDDFRRKFWEEVSKDPELSKQFNPGNKKRLSQGLAPRARNKDTVGGRRSFELHHDKPISQDGGVYDMDNLRITTPKRHIDIHRGQ | ATGGAACTGAAAAACAGCATTAGTGATTACACTGAAACTGAATTCAAAAAAATTATTGAAGACATCATCAATTGTGAAGGTGATGAAAAAAAACAGGATGATAACCTCGAGCATTTTATAAGTGTTACTGAGCATCCTAGTGGTTCTGATCTGATTTATTACCCAGAAGGTAATAATGATGGTAGCCCTGAAGCTGTTATTAAAGAGATTAAAGAATGGCGAGCTGCTAACGGTAAGTCAGGATTTAAACAGGGCTGA | MELKNSISDYTETEFKKIIEDIINCEGDEKKQDDNLEHFISVTEHPSGSDLIYYPEGNNDGSPEAVIKEIKEWRAANGKSGFKQG | ATGAAAAAAATAACAGGGATTATTTTATTGCTTCTTGCAGTCATTATTCTGGCTGCATGTCAGGCAAACTATATCCGGGATGTTCAGGGCGGGACCGTATCTCCGTCATCAACAGCTGAAGTGACCGGATTAGCAACGCAGTAA | MKKITGIILLLLAVIILAACQANYIRDVQGGTVSPSSTAEVTGLATQ | ATGAGCGGTGGAGATGGACGCGGCCATAACACGGGCGCGCATAGCACAAGTGGTAACATTAATGGTGGCC  CGACCGGGATTGGTGTAAGTGGTGGTGCTTCTGATGGTTCAGGATGGAGTTCGGAAAATAACCCGTGGGG  TGGTGGTTCCGGTAGCGGCATTCACTGGGGAGGTGGCTCCGGTCGTGGTAATGGCGGGGGTAATGGCAAT  TCCGGTGGTGGCTCGGGAACAGGCGGTAATTTGTCAGCAGTAGCTGCGCCAGTGGCATTTGGTTTTCCGG  CTCTTTCCACTCCAGGAGCTGGCGGTCTGGCTGTCAGTATTTCTGCAAGCGAATTATCGGCAGCTATTGC  TGGTATTATTGCTAAATTAAAAAAAGTAAATCTTAAATTCACTCCTTTTGGGGTTGTCTTATCTTCATTA  ATTCCGTCGGAAATAGCGAAAGATGACCCCAATATGATGTCAAAGATTGTGACGTCATTACCCGCAGATG  ATATTACTGAATCACCTGTCAGTTCATTACCTCTCGATAAGGCAACAGTAAACGTAAATGTTCGTGTTGT  TGATGATGTAAAAGACGAACGACAGAATATTTCGGTTGTTTCAGGTGTTCCGATGAGTGTTCCGGTGGTT  GATGCAAAACCTACCGAACGTCCAGGTGTTTTTACGGCATCAATTCCAGGTGCACCTGTTCTGAATATTT  CAGTTAATAACAGTACGCCAGCAGTACAGACATTAAGCCCAGGTGTTACAAATAATACTGATAAGGATGT  TCGCCCGGCAGGATTTACTCAGGGTGGTAATACCAGGGATGCAGTTATTCGATTCCCGAAGGACAGCGGT  CATAATGCCGTATATGTTTCAGTGAGTGATGTTCTTAGTCCTGACCAGGTAAAACAACGTCAGGATGAAG  AAAATCGCCGTCAGCAGGAATGGGATGCTACGCATCCGGTTGAAGCGGCTGAGCGAAATTATGAACGCGC  GCGTGCAGAGCTGAATCAGGCAAATGAAGATGTTGCCAGAAATCAGGAGCGACAGGCTAAAGCTGTTCAG  GTTTATAATTCGCGTAAAAGCGAACTTGATGCAGCGAATAAGACTCTTGCTGATGCAATAGCTGAAATAA  AACAATTTAATCGATTTGCCCATGACCCAATGGCTGGCGGTCACAGAATGTGGCAAATGGCCGGGCTTAA  AGCTCAGCGGGCGCAGACGGATGTAAATAATAAGCAGGCTGCAGATGCTGATGCTGCATTAAGTGCCGCG  CAGGAGCGCCGCAAACAGAAGGAAAATAAAGAGAAGGACGCTAAGGATAAATTAGATAAGGAGAGTAAAC  GGAATAAGCCAGGGAAGGCGACAGGTAAAGGTAAACCAGTTGGTGATAAATGGCTGGATGATGCGGGTAA  AGATTCAGGAGCGCCAATTCCAGATCGCATTGCTGATAAGTTGCGTGATAAAGAATTTAAAAACTTCGAC  GATTTTCGGAGAAAATTCTGGGAAGAAGTGTCGAAAGATCCTGAGTTAAGTAAGCAATTTAATCCAGGTA  ATAAAAAACGCTTGAGCCAAGGATTAGCTCCACGGGCTAGGAATAAAGATACTGTCGGTGGGCGTCGCTC  ATTTGAGCTTCATCACGACAAGCCAATCAGTCAGGATGGTGGTGTTTATGATATGGACAACCTCCGTATC  ACTACGCCGAAACGCCATATTGATATTCACAGAGGTCAGTAAAAATGGAACTGAAAAACAGCATTAGTGA  TTACACTGAAACTGAATTCAAAAAAATTATTGAAGACATCATCAATTGTGAAGGTGATGAAAAAAAACAG  GATGATAACCTCGAGCATTTTATAAGTGTTACTGAGCATCCTAGTGGTTCTGATCTGATTTATTACCCAG  AAGGTAATAATGATGGTAGCCCTGAAGCTGTTATTAAAGAGATTAAAGAATGGCGAGCTGCTAACGGTAA  GTCAGGATTTAAACAGGGCTGAAATATGAATGCCAGTTGTTTAAGGATGAATGACTGGCATTCTTTCACA  ACAAGGAGTCGTTATGAAAAAAATAACAGGGATTATTTTATTGCTTCTTGCAGTCATTATTCTGGCTGCA  TGTCAGGCAAACTATATCCGGGATGTTCAGGGCGGGACCGTATCTCCGTCATCAACAGCTGAAGTGACCG  GATTAGCAACGCAGTAA | ATGAGCGGTGGAGATGGACGCGGCCATAACACGGGCGCGCATAGCACAAGTGGTAACATTAATGGTGGCCCGACCGGGATTGGTGTAAGTGGTGGTGCTTCTGATGGTTCAGGATGGAGTTCGGAAAATAACCCGTGGGGTGGTGGTTCCGGTAGCGGCATTCACTGGGGAGGTGGCTCCGGTCGTGGTAATGGCGGGGGTAATGGCAATTCCGGTGGTGGCTCGGGAACAGGCGGTAATTTGTCAGCAGTAGCTGCGCCAGTGGCATTTGGTTTTCCGGCTCTTTCCACTCCAGGAGCTGGCGGTCTGGCTGTCAGTATTTCTGCAAGCGAATTATCGGCAGCTATTGCTGGTATTATTGCTAAATTAAAAAAAGTAAATCTTAAATTCACTCCTTTTGGGGTTGTCTTATCTTCATTAATTCCGTCGGAAATAGCGAAAGATGACCCCAATATGATGTCAAAGATTGTGACGTCATTACCCGCAGATGATATTACTGAATCACCTGTCAGTTCATTACCTCTCGATAAGGCAACAGTAAACGTAAATGTTCGTGTTGTTGATGATGTAAAAGACGAACGACAGAATATTTCGGTTGTTTCAGGTGTTCCGATGAGTGTTCCGGTGGTTGATGCAAAACCTACCGAACGTCCAGGTGTTTTTACGGCATCAATTCCAGGTGCACCTGTTCTGAATATTTCAGTTAATAACAGTACGCCAGCAGTACAGACATTAAGCCCAGGTGTTACAAATAATACTGATAAGGATGTTCGCCCGGCAGGATTTACTCAGGGTGGTAATACCAGGGATGCAGTTATTCGATTCCCGAAGGACAGCGGTCATAATGCCGTATATGTTTCAGTGAGTGATGTTCTTAGTCCTGACCAGGTAAAACAACGTCAGGATGAAGAAAATCGCCGTCAGCAGGAATGGGATGCTACGCATCCGGTTGAAGCGGCTGAGCGAAATTATGAACGCGCGCGTGCAGAGCTGAATCAGGCAAATGAAGATGTTGCCAGAAATCAGGAGCGACAGGCTAAAGCTGTTCAGGTTTATAATTCGCGTAAAAGCGAACTTGATGCAGCGAATAAGACTCTTGCTGATGCAATAGCTGAAATAAAACAATTTAATCGATTTGCCCATGACCCAATGGCTGGCGGTCACAGAATGTGGCAAATGGCCGGGCTTAAAGCTCAGCGGGCGCAGACGGATGTAAATAATAAGCAGGCTGCAGATGCTGATGCTGCATTAAGTGCCGCGCAGGAGCGCCGCAAACAGAAGGAAAATAAAGAGAAGGACGCTAAGGATAAATTAGATAAGGAGAGTAAACGGAATAAGCCAGGGAAGGCGACAGGTAAAGGTAAACCAGTTGGTGATAAATGGCTGGATGATGCGGGTAAAGATTCAGGAGCGCCAATTCCAGATCGCATTGCTGATAAGTTGCGTGATAAAGAATTTAAAAACTTCGACGATTTTCGGAGAAAATTCTGGGAAGAAGTGTCGAAAGATCCTGAGTTAAGTAAGCAATTTAATCCAGGTAATAAAAAACGCTTGAGCCAAGGATTAGCTCCACGGGCTAGGAATAAAGATACTGTCGGTGGGCGTCGCTCATTTGAGCTTCATCACGACAAGCCAATCAGTCAGGATGTGGTGTTTATGATATGGACAACCTCCGTATCACTACGCCGAAACGCCATATTGATATTCACAGAGGTCAGTAA  ATGGAACTGAAAAACAGCATTAGTGATTACACTGAAACTGAATTCAAAAAAATTATTGAAGACATCATCAATTGTGAAGGTGATGAAAAAAAACAGGATGATAACCTCGAGCATTTTATAAGTGTTACTGAGCATCCTAGTGGTTCTGATCTGATTTATTACCCAGAAGGTAATAATGATGGTAGCCCTGAAGCTGTTATTAAAGAGATTAAAGAATGGCGAGCTGCTAACGGTAAGTCAGGATTTAAACAGGGCTGA | MSGGDGRGHNTGAHSTSGNINGGPTGIGVSGGASDGSGWSSENNPWGGGSGSGIHWGGGSGRGNGGGNGNSGGGSGTGGNLSAVAAPVAFGFPALSTPGAGGLAVSISASELSAAIAGIIAKLKKVNLKFTPFGVVLSSLIPSEIAKDDPNMMSKIVTSLPADDITESPVSSLPLDKATVNVNVRVVDDVKDERQNISVVSGVPMSVPVVDAKPTERPGVFTASIPGAPVLNISVNNSTPAVQTLSPGVTNNTDKDVRPAGFTQGGNTRDAVIRFPKDSGHNAVYVSVSDVLSPDQVKQRQDEENRRQQEWDATHPVEAAERNYERARAELNQANEDVARNQERQAKAVQVYNSRKSELDAANKTLADAIAEIKQFNRFAHDPMAGGHRMWQMAGLKAQRAQTDVNNKQAADADAALSAAQERRKQKENKEKDAKDKLDKESKRNKPGKATGKGKPVGDKWLDDAGKDSGAPIPDRIADKLRDKEFKNFDDFRRKFWEEVSKDPELSKQFNPGNKKRLSQGLAPRARNKDTVGGRRSFELHHDKPISQDGGVYDMDNLRITTPKRHIDIHRGQ  MELKNSISDYTETEFKKIIEDIINCEGDEKKQDDNLEHFISVTEHPSGSDLIYYPEGNNDGSPEAVIKEIKEWRAANGKSGFKQG | MSGGDGRGHNTGAHSTSGNINGGPTGIGVSGGASDGSGWSSENNPWGGGSGSGIHWGGGSGRGNGGGNGNSGGGSGTGGNLSAVAAPVAFGFPALSTPGAGGLAVSISASELSAAIAGIIAKLKKVNLKFTPFGVVLSSLIPSEIAKDDPNMMSKIVTSLPADDITESPVSSLPLDKATVNVNVRVVDDVKDERQNISVVSGVPMSVPVVDAKPTERPGVFTASIPGAPVLNISVNNSTPAVQTLSPGVTNNTDKDVRPAGFTQGGNTRDAVIRFPKDSGHNAVYVSVSDVLSPDQVKQRQDEENRRQQEWDATHPVEAAERNYERARAELNQANEDVARNQERQAKAVQVYNSRKSELDAANKTLADAIAEIKQFNRFAHDPMAGGHRMWQMAGLKAQRAQTDVNNKQAADADAALSAAQERRKQKENKEKDAKDKLDKESKRNKPGKATGKGKPVGDKWLDDAGKDSGAPIPDRIADKLRDKEFKNFDDFRRKFWEEVSKDPELSKQFNPGNKKRLSQGLAPRARNKDTVGGRRSFELHHDKPISQDGGVYDMDNLRITTPKRHIDIHRGQ  MELKNSISDYTETEFKKIIEDIINCEGDEKKQDDNLEHFISVTEHPSGSDLIYYPEGNNDGSPEAVIKEIKEWRAANGKSGFKQG  MKKITGIILLLLAVIILAACQANYIRDVQGGTVSPSSTAEVTGLATQ |
| Escherichia coli plasmid ColE9-J, complete sequence  <https://www.ncbi.nlm.nih.gov/nuccore/NC_011977.1?report=graph> | Colicin E9  *ceaI*  (582 aa)  *ceiI*  immunity  (86 aa)  *ceiE*  immunity of col E5  (83 aa)  Lysis  (26 aa)  Colicin E5 lysis  (47 aa) | ATGAGCGGTGGGGATGGACGCGGCCATAACACGGGCGCGCATAGCACAAGTGGTAACATTAATGGTGGCC  CGACCGGGATTGGTGTAAGTGGTGGTGCTTCTGATGGTTCAGGATGGAGTTCGGAAAATAACCCGTGGGG  TGGTGGTTCCGGTAGCGGCATTCACTGGGGAGGTGGCTCCGGTCGTGGTAATGGCGGGGGTAATGGCAAT  TCCGGTGGTGGCTCGGGAACAGGCGGTAATTTGTCAGCAGTAGCTGCGCCAGTGGCATTTGGTTTTCCGG  CTCTTTCCACTCCAGGAGCTGGCGGTCTGGCTGTCAGTATTTCTGCAAGCGAATTATCGGCAGCTATTGC  TGGTATTATTGCTAAATTAAAAAAAGTAAATCTTAAATTCACTCCTTTTGGGGTTGTCTTATCTTCATTA  ATTCCGTCGGAAATAGCGAAAGATGACCCCAATATGATGTCAAAGATTGTGACGTCATTACCCGCAGATG  ATATTACTGAATCACCTGTCAGTTCATTACCTCTCGATAAGGCAACAGTAAACGTAAATGTTCGTGTTGT  TGATGATGTAAAAGACGAACGACAGAATATTTCGGTTGTTTCAGGTGTTCCGATGAGTGTTCCGGTGGTT  GATGCAAAACCTACCGAACGTCCAGGTGTTTTTACGGCATCAATTCCAGGTGCACCTGTTCTGAATATTT  CAGTTAATGACAGTACGCCAGCAGTACAGACATTAAGCCCAGGTGTTACAAATAATACTGATAAGGATGT  TCGCCCGGCAGGATTTACTCAGGGTGGTAATACCAGGGATGCAGTTATTCGATTCCCGAAGGACAGCGGT  CATAATGCCGTATATGTTTCAGTGAGTGATGTTCTTAGTCCTGACCAGGTAAAACAACGTCAGGATGAAG  AAAATCGCCGTCAGCAGGAATGGGATGCTACGCATCCGGTTGAAGCGGCTGAGCGAAATTATGAACGCGC  GCGTGCAGAGCTCAATCAGGCAAATGAAGATGTTGCCAGAAATCAGGAGCGACAGGCTAAAGCTGTTCAG  GTTTATAATTCGCGTAAAAGCGAACTTGATGCAGCGAATAAAACTCTTGCTGATGCAATAGCTGAAATAA  AACAATTTAATCGATTTGCCCATGACCCAATGGCTGGCGGTCACAGAATGTGGCAAATGGCCGGGCTTAA  AGCTCAGCGGGCGCAGACGGATGTAAATAATAAGCAGGCTGCATTTGATGCTGCTGCAAAAGAGAAGTCA  GATGCTGATGCTGCATTAAGTGCCGCGCAGGAGCGCCGCAAACAGAAGGAAAATAAAGAAAAGGACGCTA  AGGATAAATTAGATAAGGAGAGTAAACGGAATAAGCCAGGGAAGGCGACAGGTAAAGGTAAACCAGTTGG  TGATAAATGGCTGGATGATGCAGGTAAAGATTCAGGAGCGCCAATTCCAGATCGCATTGCTGATAAGTTG  CGTGATAAAGAATTTAAAAGCTTCGACGATTTTCGGAAGGCTGTATGGGAAGAGGTGTCGAAAGATCCTG  AGCTTAGTAAAAATTTAAACCCAAGCAATAAGTCTAGTGTTTCAAAAGGTTATTCTCCGTTTACTCCAAA  GAATCAACAGGTCGGAGGGAGAAAAGTCTATGAACTTCATCATGACAAGCCAATTAGTCAAGGTGGTGAG  GTTTATGACATGGATAATATCCGAGTGACTACACCTAAGCGACATATCGATATTCACCGAGGTAAGTAA | MSGGDGRGHNTGAHSTSGNINGGPTGIGVSGGASDGSGWSSENNPWGGGSGSGIHWGGGSGRGNGGGNGN  SGGGSGTGGNLSAVAAPVAFGFPALSTPGAGGLAVSISASELSAAIAGIIAKLKKVNLKFTPFGVVLSSL  IPSEIAKDDPNMMSKIVTSLPADDITESPVSSLPLDKATVNVNVRVVDDVKDERQNISVVSGVPMSVPVV  DAKPTERPGVFTASIPGAPVLNISVNDSTPAVQTLSPGVTNNTDKDVRPAGFTQGGNTRDAVIRFPKDSG  HNAVYVSVSDVLSPDQVKQRQDEENRRQQEWDATHPVEAAERNYERARAELNQANEDVARNQERQAKAVQ  VYNSRKSELDAANKTLADAIAEIKQFNRFAHDPMAGGHRMWQMAGLKAQRAQTDVNNKQAAFDAAAKEKS  DADAALSAAQERRKQKENKEKDAKDKLDKESKRNKPGKATGKGKPVGDKWLDDAGKDSGAPIPDRIADKL  RDKEFKSFDDFRKAVWEEVSKDPELSKNLNPSNKSSVSKGYSPFTPKNQQVGGRKVYELHHDKPISQGGE  VYDMDNIRVTTPKRHIDIHRGK | ATGGAACTGAAGCATAGCATTAGTGATTATACAGAAGCTGAATTTTTACAACTTGTAACAACAATTTGTA  ATGCGGACACTTCCAGTGAAGAAGAACTGGTTAAATTGGTTACACACTTTGAGGAAATGACTGAGCACCC  TAGTGGTAGTGATTTAATATATTACCCAAAAGAAGGTGATGATGACTCACCTTCAGGTATTGTAAACACA  GTAAAACAATGGCGAGCCGCTAACGGTAAGTCAGGATTTAAACAGGGCTAA  ATGAAGTTATCACCAAAAGCTGCAATAGAAGTTTGTAATGAAGCAGCGAAAAAAGGCTTATGGATTTTGG  GCATTGATGGTGGGCATTGGCTGAATCCTGGATTCAGGATAGATAGTTCAGCATCATGGACATATGATAT  GCCGGAGGAATACAAATCAAAAACCCCTGAAAATAATAGATTGGCTATTGAAAATATTAAAGATGATATT  GAGAATGGATACACTGCTTTCATTATCACGTTAAAGATGTAA | MELKHSISDYTEAEFLQLVTTICNADTSSEEELVKLVTHFEEMTEHPSGSDLIYYPKEGDDDSPSGIVNT  VKQWRAANGKSGFKQG  MKLSPKAAIEVCNEAAKKGLWILGIDGGHWLNPGFRIDSSASWTYDMPEEYKSKTPENNRLAIENIKDDI  ENGYTAFIITLKM | ATGAAAAAAATAACAGGGATTATTTTATTGCTTCTTGCAGTCATTATTCTGTCTGCATGGGGTTCTAAGC  CGAAAACCTAG  ATGAAAAAAATAACAGGGATTATTTTATTGCTTCTTGCAGCCATTATTCTTGCTGCATGTCAGGCAAACT  ATATCCGCGATGTTCAGGGCGGGACTGTATCCCCGTCATCCTCAGCTGAACTGACCGGATTAGCAACGCA  GTAA | MKKITGIILLLLAVIILSAWGSKPKT  MKKITGIILLLLAAIILAACQANYIRDVQGGTVSPSSSAELTGLATQ | ATGAGCGGTGGGGATGGACGCGGCCATAACACGGGCGCGCATAGCACAAGTGGTAACATTAATGGTGGCC  CGACCGGGATTGGTGTAAGTGGTGGTGCTTCTGATGGTTCAGGATGGAGTTCGGAAAATAACCCGTGGGG  TGGTGGTTCCGGTAGCGGCATTCACTGGGGAGGTGGCTCCGGTCGTGGTAATGGCGGGGGTAATGGCAAT  TCCGGTGGTGGCTCGGGAACAGGCGGTAATTTGTCAGCAGTAGCTGCGCCAGTGGCATTTGGTTTTCCGG  CTCTTTCCACTCCAGGAGCTGGCGGTCTGGCTGTCAGTATTTCTGCAAGCGAATTATCGGCAGCTATTGC  TGGTATTATTGCTAAATTAAAAAAAGTAAATCTTAAATTCACTCCTTTTGGGGTTGTCTTATCTTCATTA  ATTCCGTCGGAAATAGCGAAAGATGACCCCAATATGATGTCAAAGATTGTGACGTCATTACCCGCAGATG  ATATTACTGAATCACCTGTCAGTTCATTACCTCTCGATAAGGCAACAGTAAACGTAAATGTTCGTGTTGT  TGATGATGTAAAAGACGAACGACAGAATATTTCGGTTGTTTCAGGTGTTCCGATGAGTGTTCCGGTGGTT  GATGCAAAACCTACCGAACGTCCAGGTGTTTTTACGGCATCAATTCCAGGTGCACCTGTTCTGAATATTT  CAGTTAATGACAGTACGCCAGCAGTACAGACATTAAGCCCAGGTGTTACAAATAATACTGATAAGGATGT  TCGCCCGGCAGGATTTACTCAGGGTGGTAATACCAGGGATGCAGTTATTCGATTCCCGAAGGACAGCGGT  CATAATGCCGTATATGTTTCAGTGAGTGATGTTCTTAGTCCTGACCAGGTAAAACAACGTCAGGATGAAG  AAAATCGCCGTCAGCAGGAATGGGATGCTACGCATCCGGTTGAAGCGGCTGAGCGAAATTATGAACGCGC  GCGTGCAGAGCTCAATCAGGCAAATGAAGATGTTGCCAGAAATCAGGAGCGACAGGCTAAAGCTGTTCAG  GTTTATAATTCGCGTAAAAGCGAACTTGATGCAGCGAATAAAACTCTTGCTGATGCAATAGCTGAAATAA  AACAATTTAATCGATTTGCCCATGACCCAATGGCTGGCGGTCACAGAATGTGGCAAATGGCCGGGCTTAA  AGCTCAGCGGGCGCAGACGGATGTAAATAATAAGCAGGCTGCATTTGATGCTGCTGCAAAAGAGAAGTCA  GATGCTGATGCTGCATTAAGTGCCGCGCAGGAGCGCCGCAAACAGAAGGAAAATAAAGAAAAGGACGCTA  AGGATAAATTAGATAAGGAGAGTAAACGGAATAAGCCAGGGAAGGCGACAGGTAAAGGTAAACCAGTTGG  TGATAAATGGCTGGATGATGCAGGTAAAGATTCAGGAGCGCCAATTCCAGATCGCATTGCTGATAAGTTG  CGTGATAAAGAATTTAAAAGCTTCGACGATTTTCGGAAGGCTGTATGGGAAGAGGTGTCGAAAGATCCTG  AGCTTAGTAAAAATTTAAACCCAAGCAATAAGTCTAGTGTTTCAAAAGGTTATTCTCCGTTTACTCCAAA  GAATCAACAGGTCGGAGGGAGAAAAGTCTATGAACTTCATCATGACAAGCCAATTAGTCAAGGTGGTGAG  GTTTATGACATGGATAATATCCGAGTGACTACACCTAAGCGACATATCGATATTCACCGAGGTAAGTAAA  ATGGAACTGAAGCATAGCATTAGTGATTATACAGAAGCTGAATTTTTACAACTTGTAACAACAATTTGTA  ATGCGGACACTTCCAGTGAAGAAGAACTGGTTAAATTGGTTACACACTTTGAGGAAATGACTGAGCACCC  TAGTGGTAGTGATTTAATATATTACCCAAAAGAAGGTGATGATGACTCACCTTCAGGTATTGTAAACACA  GTAAAACAATGGCGAGCCGCTAACGGTAAGTCAGGATTTAAACAGGGCTAAAATATGAGTGCCGGTTGTT  TAAGGATGAATGGCTGGCATTCTTTCACAACAAGGAGTCGTTATGAAAAAAATAACAGGGATTATTTTAT  TGCTTCTTGCAGTCATTATTCTGTCTGCATGGGGTTCTAAGCCGAAAACCTAGAAAATTCCGTAACCAAA  GCCAGTAATTGACAGATTTGCATGACGTTGAATAGGTGACGGGTTATGTGACGAAATCTGATGCAGAAAT  CGTTGTTTCAGTGACAGTCACTCAATCGGTCGTTTATGTGACAACCCACGCCGTTACTGGTCGCGGAAAA  ATCCAGGTTTCCGGCCTGGAACCCCGAAATGATCCAGCAACAGTGTATGGTTCTCCTGGTAAATATGTTG  TTGTCAATGATCGTACTGGTGAGGTTACTCAGATTAGTGATAAGACAGATCCGGGTTGGGTGGACGATTC  GAGAATTCAATGGGGAAATAAAAATGACCAATAAATTATTTGAACATACGGTGTTATATGATAGTGGTGA  TGCCTTTTTTGAATTAAAAGGAAATGCTTCTATGAAGTTATCACCAAAAGCTGCAATAGAAGTTTGTAAT  GAAGCAGCGAAAAAAGGCTTATGGATTTTGGGCATTGATGGTGGGCATTGGCTGAATCCTGGATTCAGGA  TAGATAGTTCAGCATCATGGACATATGATATGCCGGAGGAATACAAATCAAAAACCCCTGAAAATAATAG  ATTGGCTATTGAAAATATTAAAGATGATATTGAGAATGGATACACTGCTTTCATTATCACGTTAAAGATG  TAAATAGTGTTATAGAATTTTATGTTTCATGGATGATTTCAACCTTTGGATTTCAGGTTTTTATGGATGA  ATGCCTGAGTCCATATATACAAGGAACAGTATGAAAAAAATAACAGGGATTATTTTATTGCTTCTTGCAG  CCATTATTCTTGCTGCATGTCAGGCAAACTATATCCGCGATGTTCAGGGCGGGACTGTATCCCCGTCATC  CTCAGCTGAACTGACCGGATTAGCAACGCAGTAA | ATGAGCGGTGGGGATGGACGCGGCCATAACACGGGCGCGCATAGCACAAGTGGTAACATTAATGGTGGCC  CGACCGGGATTGGTGTAAGTGGTGGTGCTTCTGATGGTTCAGGATGGAGTTCGGAAAATAACCCGTGGGG  TGGTGGTTCCGGTAGCGGCATTCACTGGGGAGGTGGCTCCGGTCGTGGTAATGGCGGGGGTAATGGCAAT  TCCGGTGGTGGCTCGGGAACAGGCGGTAATTTGTCAGCAGTAGCTGCGCCAGTGGCATTTGGTTTTCCGG  CTCTTTCCACTCCAGGAGCTGGCGGTCTGGCTGTCAGTATTTCTGCAAGCGAATTATCGGCAGCTATTGC  TGGTATTATTGCTAAATTAAAAAAAGTAAATCTTAAATTCACTCCTTTTGGGGTTGTCTTATCTTCATTA  ATTCCGTCGGAAATAGCGAAAGATGACCCCAATATGATGTCAAAGATTGTGACGTCATTACCCGCAGATG  ATATTACTGAATCACCTGTCAGTTCATTACCTCTCGATAAGGCAACAGTAAACGTAAATGTTCGTGTTGT  TGATGATGTAAAAGACGAACGACAGAATATTTCGGTTGTTTCAGGTGTTCCGATGAGTGTTCCGGTGGTT  GATGCAAAACCTACCGAACGTCCAGGTGTTTTTACGGCATCAATTCCAGGTGCACCTGTTCTGAATATTT  CAGTTAATGACAGTACGCCAGCAGTACAGACATTAAGCCCAGGTGTTACAAATAATACTGATAAGGATGT  TCGCCCGGCAGGATTTACTCAGGGTGGTAATACCAGGGATGCAGTTATTCGATTCCCGAAGGACAGCGGT  CATAATGCCGTATATGTTTCAGTGAGTGATGTTCTTAGTCCTGACCAGGTAAAACAACGTCAGGATGAAG  AAAATCGCCGTCAGCAGGAATGGGATGCTACGCATCCGGTTGAAGCGGCTGAGCGAAATTATGAACGCGC  GCGTGCAGAGCTCAATCAGGCAAATGAAGATGTTGCCAGAAATCAGGAGCGACAGGCTAAAGCTGTTCAG  GTTTATAATTCGCGTAAAAGCGAACTTGATGCAGCGAATAAAACTCTTGCTGATGCAATAGCTGAAATAA  AACAATTTAATCGATTTGCCCATGACCCAATGGCTGGCGGTCACAGAATGTGGCAAATGGCCGGGCTTAA  AGCTCAGCGGGCGCAGACGGATGTAAATAATAAGCAGGCTGCATTTGATGCTGCTGCAAAAGAGAAGTCA  GATGCTGATGCTGCATTAAGTGCCGCGCAGGAGCGCCGCAAACAGAAGGAAAATAAAGAAAAGGACGCTA  AGGATAAATTAGATAAGGAGAGTAAACGGAATAAGCCAGGGAAGGCGACAGGTAAAGGTAAACCAGTTGG  TGATAAATGGCTGGATGATGCAGGTAAAGATTCAGGAGCGCCAATTCCAGATCGCATTGCTGATAAGTTG  CGTGATAAAGAATTTAAAAGCTTCGACGATTTTCGGAAGGCTGTATGGGAAGAGGTGTCGAAAGATCCTG  AGCTTAGTAAAAATTTAAACCCAAGCAATAAGTCTAGTGTTTCAAAAGGTTATTCTCCGTTTACTCCAAA  GAATCAACAGGTCGGAGGGAGAAAAGTCTATGAACTTCATCATGACAAGCCAATTAGTCAAGGTGGTGAG  GTTTATGACATGGATAATATCCGAGTGACTACACCTAAGCGACATATCGATATTCACCGAGGTAAGTAA  ATGGAACTGAAGCATAGCATTAGTGATTATACAGAAGCTGAATTTTTACAACTTGTAACAACAATTTGTA  ATGCGGACACTTCCAGTGAAGAAGAACTGGTTAAATTGGTTACACACTTTGAGGAAATGACTGAGCACCC  TAGTGGTAGTGATTTAATATATTACCCAAAAGAAGGTGATGATGACTCACCTTCAGGTATTGTAAACACA  GTAAAACAATGGCGAGCCGCTAACGGTAAGTCAGGATTTAAACAGGGCTAA  ATGAAGTTATCACCAAAAGCTGCAATAGAAGTTTGTAATGAAGCAGCGAAAAAAGGCTTATGGATTTTGG  GCATTGATGGTGGGCATTGGCTGAATCCTGGATTCAGGATAGATAGTTCAGCATCATGGACATATGATAT  GCCGGAGGAATACAAATCAAAAACCCCTGAAAATAATAGATTGGCTATTGAAAATATTAAAGATGATATT  GAGAATGGATACACTGCTTTCATTATCACGTTAAAGATGTAA | MSGGDGRGHNTGAHSTSGNINGGPTGIGVSGGASDGSGWSSENNPWGGGSGSGIHWGGGSGRGNGGGNGN  SGGGSGTGGNLSAVAAPVAFGFPALSTPGAGGLAVSISASELSAAIAGIIAKLKKVNLKFTPFGVVLSSL  IPSEIAKDDPNMMSKIVTSLPADDITESPVSSLPLDKATVNVNVRVVDDVKDERQNISVVSGVPMSVPVV  DAKPTERPGVFTASIPGAPVLNISVNDSTPAVQTLSPGVTNNTDKDVRPAGFTQGGNTRDAVIRFPKDSG  HNAVYVSVSDVLSPDQVKQRQDEENRRQQEWDATHPVEAAERNYERARAELNQANEDVARNQERQAKAVQ  VYNSRKSELDAANKTLADAIAEIKQFNRFAHDPMAGGHRMWQMAGLKAQRAQTDVNNKQAAFDAAAKEKS  DADAALSAAQERRKQKENKEKDAKDKLDKESKRNKPGKATGKGKPVGDKWLDDAGKDSGAPIPDRIADKL  RDKEFKSFDDFRKAVWEEVSKDPELSKNLNPSNKSSVSKGYSPFTPKNQQVGGRKVYELHHDKPISQGGE  VYDMDNIRVTTPKRHIDIHRGK  MELKHSISDYTEAEFLQLVTTICNADTSSEEELVKLVTHFEEMTEHPSGSDLIYYPKEGDDDSPSGIVNT  VKQWRAANGKSGFKQG  MKLSPKAAIEVCNEAAKKGLWILGIDGGHWLNPGFRIDSSASWTYDMPEEYKSKTPENNRLAIENIKDDI  ENGYTAFIITLKM | MSGGDGRGHNTGAHSTSGNINGGPTGIGVSGGASDGSGWSSENNPWGGGSGSGIHWGGGSGRGNGGGNGN  SGGGSGTGGNLSAVAAPVAFGFPALSTPGAGGLAVSISASELSAAIAGIIAKLKKVNLKFTPFGVVLSSL  IPSEIAKDDPNMMSKIVTSLPADDITESPVSSLPLDKATVNVNVRVVDDVKDERQNISVVSGVPMSVPVV  DAKPTERPGVFTASIPGAPVLNISVNDSTPAVQTLSPGVTNNTDKDVRPAGFTQGGNTRDAVIRFPKDSG  HNAVYVSVSDVLSPDQVKQRQDEENRRQQEWDATHPVEAAERNYERARAELNQANEDVARNQERQAKAVQ  VYNSRKSELDAANKTLADAIAEIKQFNRFAHDPMAGGHRMWQMAGLKAQRAQTDVNNKQAAFDAAAKEKS  DADAALSAAQERRKQKENKEKDAKDKLDKESKRNKPGKATGKGKPVGDKWLDDAGKDSGAPIPDRIADKL  RDKEFKSFDDFRKAVWEEVSKDPELSKNLNPSNKSSVSKGYSPFTPKNQQVGGRKVYELHHDKPISQGGE  VYDMDNIRVTTPKRHIDIHRGK  MELKHSISDYTEAEFLQLVTTICNADTSSEEELVKLVTHFEEMTEHPSGSDLIYYPKEGDDDSPSGIVNT  VKQWRAANGKSGFKQG  MKLSPKAAIEVCNEAAKKGLWILGIDGGHWLNPGFRIDSSASWTYDMPEEYKSKTPENNRLAIENIKDDI  ENGYTAFIITLKM  MKKITGIILLLLAVIILSAWGSKPKT  MKKITGIILLLLAAIILAACQANYIRDVQGGTVSPSSSAELTGLATQ |
| E.coli plasmid pColD-157 DNA  <https://www.ncbi.nlm.nih.gov/nuccore/Y10412.1?report=graph> | Colicin D  *cda*  (697 aa)  Immunity  *cdi*  (87 aa)  Lysis  *cdl*  (48 aa) | ATGAGTGATTACGAAGGTAGTGGTCCGACAGAAGGTATTGATTATGGGCACTCGATGGTCGTGTGGCCGT  CAACAGGACTGATTTCGGGTGGTGATGTGAAACCAGGAGGCTCATCAGGTATCGCTCCATCCATGCCTCC  GGGATGGGGGGATTACAGCCCACAGGGTATCGCACTTGTACAAAGTGTTCTTTTTCCGGGAATTATTCGC  CGGATTATTCTGGATAAGGAACTTGAAGAGGGAGACTGGTCGGGATGGTCTGTCAGTGTGCATAGCCCTT  GGGGAAACGAGAAAGTTTCCGCTGCACGAACAGTTCTTGAGAATGGTTTACGTGGTGGTTTGCCAGAACC  GTCTCGCCCGGCTGCTGTTTCTTTTGCCCGTCTGGAGCCTGCTTCCGGAAATGAGCAAAAAATTATTCGT  CTTATGGTTACACAGCAACTGGAACAGGTAACGGATATTCCGGCCAGCCAGTTACCAGCAGCGGGTAATA  ATGTACCGGTGAAATATCGTCTGATGGACCTTATGCAGAACGGTACGCAATATATGGCTATTATCGGAGG  TATTCCGATGACAGTGCCGGTAGTTGATGCCGTTCCAGTTCCGGACCGGAGTCGTCCGGGAACAAATATT  AAAGATGTTTACAGTGCCCCTGTATCACCAAATCTACCGGACCTGGTATTAAGTGTGGGTCAGATGAATA  CTCCGGTTCTGTCTAATCCCGAAATTCAGGAAGAAGGAGTTATTGCTGAGACAGGTAATTATGTTGAGGC  TGGTTATACGATGTCCAGTAATAATCATGATGTCATTGTTCGTTTTCCTGATGGCAGTGGTGTTTCTCCG  TTATATATTTCAACTGTAGAGATTCTGGACAGTAATGTTCTGAGTCAGCGTCAGGAAGCCGAAAATAAAG  CAAAGGATGATTTCAGAGTCAAGAAAGAAGAAGCGGTAGCCCGGGCTGAAGCTGAAAAAGCAAAAGCCGA  GTTATTTAGTAAAGCAGGTGTGAACCAGCCTCCTGTATATACACAGGAAATGATGGAAAGGGCTAATTCA  GTAATGAATGAGCAGGGAGCTCTTGTTCTGAATAACACTGCCAGTTCTGTACAACTGGCGATGACCGGAA  CTGGTGTCTGGACTGCTGCTGGTGATATTGCAGGCAACATAAGCAAGTTTTTCAGTAATGCTCTGGAGAA  AGTAACCATACCCGAAGTGAGTCCCCTGCTTATGCGGATTTCTCTTGGTGCTCTGTGGTTTCATTCAGAA  GAAGCTGGAGCAGGAAGTGATATCGTGCCGGGGCGAAATCTGGAGGCGATGTTTTCGCTGAGTGCTCAGA  TGTTGGCTGGACAGGGCGTGGTCATTGAACCTGGTGCGACGAGTGTAAATCTGCCTGTTCGTGGACAATT  GATAAACAGTAACGGGCAATTAGCTCTGGATTTACTGAAAACAGGGAATGAAAGTATCCCAGCTGCGGTT  CCTGTTCTTAATGCTGTCCGTGATACAGCAACAGGACTGGATAAAATCACGTTACCAGCAGTAGTAGGCG  CACCTTCCCGGACGATTCTGGTTAATCCGGTACCACAACCTTCGGTGCCAACAGATACAGGTAATCATCA  ACCCGTTCCGGTTACACCAGTGCATACAGGAACGGAAGTAAAACCGGTCGAAATGCCAGTAACGACGATT  ACTCCTGTTTCTGATGTTGGTGGACTACGAGATTTTATTTACTGGCGTCCTGATGCTGCTGGGACTGGTG  TTGAAGCTGTTTATGTGATGCTTAATGATCCTCTGGATTCAGGGCGATTTTCGCGTAAACAACTTGATAA  AAAATATAAACATGCTGGTGATTTTGGTATTAGTGATACAAAAAAGAATCGTGAAACTCTTACTAAATTC  AGGGATGCTATTGAGGAGCATTTATCGGATAAGGATACAGTAGAGAAAGGAACATACCGAAGAGAAAAAG  GTTCAAAAGTTTATTTTAATCCTAATACGATGAATGTGGTTATAATTAAGTCAAATGGTGAGTTCTTATC  TGGGTGGAAAATAAATCCAGATGCGGATAATGGTCGAATTTATTTAGAGACAGGTGAACTATGA | MSDYEGSGPTEGIDYGHSMVVWPSTGLISGGDVKPGGSSGIAPSMPPGWGDYSPQGIALVQSVLFPGIIR  RIILDKELEEGDWSGWSVSVHSPWGNEKVSAARTVLENGLRGGLPEPSRPAAVSFARLEPASGNEQKIIR  LMVTQQLEQVTDIPASQLPAAGNNVPVKYRLMDLMQNGTQYMAIIGGIPMTVPVVDAVPVPDRSRPGTNI  KDVYSAPVSPNLPDLVLSVGQMNTPVLSNPEIQEEGVIAETGNYVEAGYTMSSNNHDVIVRFPDGSGVSP  LYISTVEILDSNVLSQRQEAENKAKDDFRVKKEEAVARAEAEKAKAELFSKAGVNQPPVYTQEMMERANS  VMNEQGALVLNNTASSVQLAMTGTGVWTAAGDIAGNISKFFSNALEKVTIPEVSPLLMRISLGALWFHSE  EAGAGSDIVPGRNLEAMFSLSAQMLAGQGVVIEPGATSVNLPVRGQLINSNGQLALDLLKTGNESIPAAV  PVLNAVRDTATGLDKITLPAVVGAPSRTILVNPVPQPSVPTDTGNHQPVPVTPVHTGTEVKPVEMPVTTI  TPVSDVGGLRDFIYWRPDAAGTGVEAVYVMLNDPLDSGRFSRKQLDKKYKHAGDFGISDTKKNRETLTKF  RDAIEEHLSDKDTVEKGTYRREKGSKVYFNPNTMNVVIIKSNGEFLSGWKINPDADNGRIYLETGEL | ATGAACAAGATGGCAATGATCGATTTGGCGAAATTATTTTTAGCTTCGAAAATTACAGCAATTGAGTTTT  CAGAGCGAATTTGTGTTGAACGGAGAAGATTGTATGGTGTTAAGGATTTGTCTCCGAATATATTAAATTG  TGGGGAAGAGTTGTTTATGGCTGCTGAGCGATTTGAGCCTGATGCAGATAGGGCTGATTATGAAATTGAT  GATAATGGACTTAAGGTTGAGGTCCGATCTATCTTGGAAAAATTTAAATTATAA | MNKMAMIDLAKLFLASKITAIEFSERICVERRRLYGVKDLSPNILNCGEELFMAAERFEPDADRADYEID  DNGLKVEVRSILEKFKL | ATGAAAAAAAAGATAGGGGGATCTATGATTCTCGTTCTTGCTGTTTTGTGTCTGACTGCTTGCCAAGCTA  ACTATGTAAGGGATGTTCAGGGTGGAACAATTGCTCCATCATCATCTTCGAAACTAATCGGGGTGGCGGT  TCAGTGA | MKKKIGGSMILVLAVLCLTACQANYVRDVQGGTIAPSSSSKLIGVAVQ | ATGAGTGATTACGAAGGTAGTGGTCCGACAGAAGGTATTGATTATGGGCACTCGATGGTCGTGTGGCCGT  CAACAGGACTGATTTCGGGTGGTGATGTGAAACCAGGAGGCTCATCAGGTATCGCTCCATCCATGCCTCC  GGGATGGGGGGATTACAGCCCACAGGGTATCGCACTTGTACAAAGTGTTCTTTTTCCGGGAATTATTCGC  CGGATTATTCTGGATAAGGAACTTGAAGAGGGAGACTGGTCGGGATGGTCTGTCAGTGTGCATAGCCCTT  GGGGAAACGAGAAAGTTTCCGCTGCACGAACAGTTCTTGAGAATGGTTTACGTGGTGGTTTGCCAGAACC  GTCTCGCCCGGCTGCTGTTTCTTTTGCCCGTCTGGAGCCTGCTTCCGGAAATGAGCAAAAAATTATTCGT  CTTATGGTTACACAGCAACTGGAACAGGTAACGGATATTCCGGCCAGCCAGTTACCAGCAGCGGGTAATA  ATGTACCGGTGAAATATCGTCTGATGGACCTTATGCAGAACGGTACGCAATATATGGCTATTATCGGAGG  TATTCCGATGACAGTGCCGGTAGTTGATGCCGTTCCAGTTCCGGACCGGAGTCGTCCGGGAACAAATATT  AAAGATGTTTACAGTGCCCCTGTATCACCAAATCTACCGGACCTGGTATTAAGTGTGGGTCAGATGAATA  CTCCGGTTCTGTCTAATCCCGAAATTCAGGAAGAAGGAGTTATTGCTGAGACAGGTAATTATGTTGAGGC  TGGTTATACGATGTCCAGTAATAATCATGATGTCATTGTTCGTTTTCCTGATGGCAGTGGTGTTTCTCCG  TTATATATTTCAACTGTAGAGATTCTGGACAGTAATGTTCTGAGTCAGCGTCAGGAAGCCGAAAATAAAG  CAAAGGATGATTTCAGAGTCAAGAAAGAAGAAGCGGTAGCCCGGGCTGAAGCTGAAAAAGCAAAAGCCGA  GTTATTTAGTAAAGCAGGTGTGAACCAGCCTCCTGTATATACACAGGAAATGATGGAAAGGGCTAATTCA  GTAATGAATGAGCAGGGAGCTCTTGTTCTGAATAACACTGCCAGTTCTGTACAACTGGCGATGACCGGAA  CTGGTGTCTGGACTGCTGCTGGTGATATTGCAGGCAACATAAGCAAGTTTTTCAGTAATGCTCTGGAGAA  AGTAACCATACCCGAAGTGAGTCCCCTGCTTATGCGGATTTCTCTTGGTGCTCTGTGGTTTCATTCAGAA  GAAGCTGGAGCAGGAAGTGATATCGTGCCGGGGCGAAATCTGGAGGCGATGTTTTCGCTGAGTGCTCAGA  TGTTGGCTGGACAGGGCGTGGTCATTGAACCTGGTGCGACGAGTGTAAATCTGCCTGTTCGTGGACAATT  GATAAACAGTAACGGGCAATTAGCTCTGGATTTACTGAAAACAGGGAATGAAAGTATCCCAGCTGCGGTT  CCTGTTCTTAATGCTGTCCGTGATACAGCAACAGGACTGGATAAAATCACGTTACCAGCAGTAGTAGGCG  CACCTTCCCGGACGATTCTGGTTAATCCGGTACCACAACCTTCGGTGCCAACAGATACAGGTAATCATCA  ACCCGTTCCGGTTACACCAGTGCATACAGGAACGGAAGTAAAACCGGTCGAAATGCCAGTAACGACGATT  ACTCCTGTTTCTGATGTTGGTGGACTACGAGATTTTATTTACTGGCGTCCTGATGCTGCTGGGACTGGTG  TTGAAGCTGTTTATGTGATGCTTAATGATCCTCTGGATTCAGGGCGATTTTCGCGTAAACAACTTGATAA  AAAATATAAACATGCTGGTGATTTTGGTATTAGTGATACAAAAAAGAATCGTGAAACTCTTACTAAATTC  AGGGATGCTATTGAGGAGCATTTATCGGATAAGGATACAGTAGAGAAAGGAACATACCGAAGAGAAAAAG  GTTCAAAAGTTTATTTTAATCCTAATACGATGAATGTGGTTATAATTAAGTCAAATGGTGAGTTCTTATC  TGGGTGGAAAATAAATCCAGATGCGGATAATGGTCGAATTTATTTAGAGACAGGTGAACTATGAACAAGA  TGGCAATGATCGATTTGGCGAAATTATTTTTAGCTTCGAAAATTACAGCAATTGAGTTTTCAGAGCGAAT  TTGTGTTGAACGGAGAAGATTGTATGGTGTTAAGGATTTGTCTCCGAATATATTAAATTGTGGGGAAGAG  TTGTTTATGGCTGCTGAGCGATTTGAGCCTGATGCAGATAGGGCTGATTATGAAATTGATGATAATGGAC  TTAAGGTTGAGGTCCGATCTATCTTGGAAAAATTTAAATTATAATCATGAGTGATTTTAATATCTGGCAT  TTAAGTATAAATGTCAGATATTCACTCATGACAAGGAGATGTCATGAAAAAAAAGATAGGGGGATCTATG  ATTCTCGTTCTTGCTGTTTTGTGTCTGACTGCTTGCCAAGCTAACTATGTAAGGGATGTTCAGGGTGGAA  CAATTGCTCCATCATCATCTTCGAAACTAATCGGGGTGGCGGTTCAGTGA | ATGAGTGATTACGAAGGTAGTGGTCCGACAGAAGGTATTGATTATGGGCACTCGATGGTCGTGTGGCCGT  CAACAGGACTGATTTCGGGTGGTGATGTGAAACCAGGAGGCTCATCAGGTATCGCTCCATCCATGCCTCC  GGGATGGGGGGATTACAGCCCACAGGGTATCGCACTTGTACAAAGTGTTCTTTTTCCGGGAATTATTCGC  CGGATTATTCTGGATAAGGAACTTGAAGAGGGAGACTGGTCGGGATGGTCTGTCAGTGTGCATAGCCCTT  GGGGAAACGAGAAAGTTTCCGCTGCACGAACAGTTCTTGAGAATGGTTTACGTGGTGGTTTGCCAGAACC  GTCTCGCCCGGCTGCTGTTTCTTTTGCCCGTCTGGAGCCTGCTTCCGGAAATGAGCAAAAAATTATTCGT  CTTATGGTTACACAGCAACTGGAACAGGTAACGGATATTCCGGCCAGCCAGTTACCAGCAGCGGGTAATA  ATGTACCGGTGAAATATCGTCTGATGGACCTTATGCAGAACGGTACGCAATATATGGCTATTATCGGAGG  TATTCCGATGACAGTGCCGGTAGTTGATGCCGTTCCAGTTCCGGACCGGAGTCGTCCGGGAACAAATATT  AAAGATGTTTACAGTGCCCCTGTATCACCAAATCTACCGGACCTGGTATTAAGTGTGGGTCAGATGAATA  CTCCGGTTCTGTCTAATCCCGAAATTCAGGAAGAAGGAGTTATTGCTGAGACAGGTAATTATGTTGAGGC  TGGTTATACGATGTCCAGTAATAATCATGATGTCATTGTTCGTTTTCCTGATGGCAGTGGTGTTTCTCCG  TTATATATTTCAACTGTAGAGATTCTGGACAGTAATGTTCTGAGTCAGCGTCAGGAAGCCGAAAATAAAG  CAAAGGATGATTTCAGAGTCAAGAAAGAAGAAGCGGTAGCCCGGGCTGAAGCTGAAAAAGCAAAAGCCGA  GTTATTTAGTAAAGCAGGTGTGAACCAGCCTCCTGTATATACACAGGAAATGATGGAAAGGGCTAATTCA  GTAATGAATGAGCAGGGAGCTCTTGTTCTGAATAACACTGCCAGTTCTGTACAACTGGCGATGACCGGAA  CTGGTGTCTGGACTGCTGCTGGTGATATTGCAGGCAACATAAGCAAGTTTTTCAGTAATGCTCTGGAGAA  AGTAACCATACCCGAAGTGAGTCCCCTGCTTATGCGGATTTCTCTTGGTGCTCTGTGGTTTCATTCAGAA  GAAGCTGGAGCAGGAAGTGATATCGTGCCGGGGCGAAATCTGGAGGCGATGTTTTCGCTGAGTGCTCAGA  TGTTGGCTGGACAGGGCGTGGTCATTGAACCTGGTGCGACGAGTGTAAATCTGCCTGTTCGTGGACAATT  GATAAACAGTAACGGGCAATTAGCTCTGGATTTACTGAAAACAGGGAATGAAAGTATCCCAGCTGCGGTT  CCTGTTCTTAATGCTGTCCGTGATACAGCAACAGGACTGGATAAAATCACGTTACCAGCAGTAGTAGGCG  CACCTTCCCGGACGATTCTGGTTAATCCGGTACCACAACCTTCGGTGCCAACAGATACAGGTAATCATCA  ACCCGTTCCGGTTACACCAGTGCATACAGGAACGGAAGTAAAACCGGTCGAAATGCCAGTAACGACGATT  ACTCCTGTTTCTGATGTTGGTGGACTACGAGATTTTATTTACTGGCGTCCTGATGCTGCTGGGACTGGTG  TTGAAGCTGTTTATGTGATGCTTAATGATCCTCTGGATTCAGGGCGATTTTCGCGTAAACAACTTGATAA  AAAATATAAACATGCTGGTGATTTTGGTATTAGTGATACAAAAAAGAATCGTGAAACTCTTACTAAATTC  AGGGATGCTATTGAGGAGCATTTATCGGATAAGGATACAGTAGAGAAAGGAACATACCGAAGAGAAAAAG  GTTCAAAAGTTTATTTTAATCCTAATACGATGAATGTGGTTATAATTAAGTCAAATGGTGAGTTCTTATC  TGGGTGGAAAATAAATCCAGATGCGGATAATGGTCGAATTTATTTAGAGACAGGTGAACTATGA  ATGAACAAGATGGCAATGATCGATTTGGCGAAATTATTTTTAGCTTCGAAAATTACAGCAATTGAGTTTT  CAGAGCGAATTTGTGTTGAACGGAGAAGATTGTATGGTGTTAAGGATTTGTCTCCGAATATATTAAATTG  TGGGGAAGAGTTGTTTATGGCTGCTGAGCGATTTGAGCCTGATGCAGATAGGGCTGATTATGAAATTGAT  GATAATGGACTTAAGGTTGAGGTCCGATCTATCTTGGAAAAATTTAAATTATAA | MSDYEGSGPTEGIDYGHSMVVWPSTGLISGGDVKPGGSSGIAPSMPPGWGDYSPQGIALVQSVLFPGIIR  RIILDKELEEGDWSGWSVSVHSPWGNEKVSAARTVLENGLRGGLPEPSRPAAVSFARLEPASGNEQKIIR  LMVTQQLEQVTDIPASQLPAAGNNVPVKYRLMDLMQNGTQYMAIIGGIPMTVPVVDAVPVPDRSRPGTNI  KDVYSAPVSPNLPDLVLSVGQMNTPVLSNPEIQEEGVIAETGNYVEAGYTMSSNNHDVIVRFPDGSGVSP  LYISTVEILDSNVLSQRQEAENKAKDDFRVKKEEAVARAEAEKAKAELFSKAGVNQPPVYTQEMMERANS  VMNEQGALVLNNTASSVQLAMTGTGVWTAAGDIAGNISKFFSNALEKVTIPEVSPLLMRISLGALWFHSE  EAGAGSDIVPGRNLEAMFSLSAQMLAGQGVVIEPGATSVNLPVRGQLINSNGQLALDLLKTGNESIPAAV  PVLNAVRDTATGLDKITLPAVVGAPSRTILVNPVPQPSVPTDTGNHQPVPVTPVHTGTEVKPVEMPVTTI  TPVSDVGGLRDFIYWRPDAAGTGVEAVYVMLNDPLDSGRFSRKQLDKKYKHAGDFGISDTKKNRETLTKF  RDAIEEHLSDKDTVEKGTYRREKGSKVYFNPNTMNVVIIKSNGEFLSGWKINPDADNGRIYLETGEL  MNKMAMIDLAKLFLASKITAIEFSERICVERRRLYGVKDLSPNILNCGEELFMAAERFEPDADRADYEID  DNGLKVEVRSILEKFKL | MSDYEGSGPTEGIDYGHSMVVWPSTGLISGGDVKPGGSSGIAPSMPPGWGDYSPQGIALVQSVLFPGIIR  RIILDKELEEGDWSGWSVSVHSPWGNEKVSAARTVLENGLRGGLPEPSRPAAVSFARLEPASGNEQKIIR  LMVTQQLEQVTDIPASQLPAAGNNVPVKYRLMDLMQNGTQYMAIIGGIPMTVPVVDAVPVPDRSRPGTNI  KDVYSAPVSPNLPDLVLSVGQMNTPVLSNPEIQEEGVIAETGNYVEAGYTMSSNNHDVIVRFPDGSGVSP  LYISTVEILDSNVLSQRQEAENKAKDDFRVKKEEAVARAEAEKAKAELFSKAGVNQPPVYTQEMMERANS  VMNEQGALVLNNTASSVQLAMTGTGVWTAAGDIAGNISKFFSNALEKVTIPEVSPLLMRISLGALWFHSE  EAGAGSDIVPGRNLEAMFSLSAQMLAGQGVVIEPGATSVNLPVRGQLINSNGQLALDLLKTGNESIPAAV  PVLNAVRDTATGLDKITLPAVVGAPSRTILVNPVPQPSVPTDTGNHQPVPVTPVHTGTEVKPVEMPVTTI  TPVSDVGGLRDFIYWRPDAAGTGVEAVYVMLNDPLDSGRFSRKQLDKKYKHAGDFGISDTKKNRETLTKF  RDAIEEHLSDKDTVEKGTYRREKGSKVYFNPNTMNVVIIKSNGEFLSGWKINPDADNGRIYLETGEL  MNKMAMIDLAKLFLASKITAIEFSERICVERRRLYGVKDLSPNILNCGEELFMAAERFEPDADRADYEID  DNGLKVEVRSILEKFKL  MKKKIGGSMILVLAVLCLTACQANYVRDVQGGTIAPSSSSKLIGVAVQ |
| Escherichia coli K-12 P678-54 plasmid CloDF13, complete sequence  <https://www.ncbi.nlm.nih.gov/nuccore/NC_002119.1?report=graph> | Cloacin DF13  *ccl*  (561 aa)  Immunity  *cim*  (85 aa)  Lysis  *Cex*  (49 aa) | ATGAGTGGTGGAGACGGTCGAGGTCCGGGTAATTCAGGTCTGGGACATAATGGTGGTCAG  GCCAGTGGGAATGTGAACGGTACGTCTGGTAAAGGTGGCCCTTCATCAGGTGGTGGTACG  GATCCAAACAGCGGGCCGGGCTGGGGTACGACGCATACGCCGAATGGAGATATTCATAAC  TATAATCCGGGGGAGTTTGGTAACGGCGGGAGTAAACCCGGTGGTAATGGTGGTAACAGC  GGCAATCATAGCGGTAGCTCTGGTGGTGGACAGTCTTCGGCCACCGCGATGGCCTTCGGT  CTGCCTGCCCTGGCTACACCGGGGGCTGAAGGACCGGCTTTATCCTTTTCCGGCGATGCG  CTGTCGTCAGCCGTTGCCGATGTGCTGGCTGCCCTGAAAGGTCCGTTTAAGTTTGGTCTG  TGGGGGATTGCGATCTATGGTGTACTGCCTTCTGAGATTGCAAAAGATGATCCGAAAATG  ATGTCAAAAATTATGACGTCATTACCGGCCGATACGGTGACGGAGACTCCGGCAAGTACT  TTACCGCTGGACCAAGCGACGGTTCGTGTCAGACAACGGGTTGTGGATGTGGTGAAGGAT  GAACGGCAGCATATTGCGGTTGTCGCGGGTCGGCCAATGAGTGTTCCTGTGGTGGATGCG  AAACCGACAAAACGTCCGGGGGTATTCAGTGTGTCTATTCCGGGTCTCCCGGCTCTGCAG  GTGAGCGTACCGAAAGGTGTTCCGGCAGCGAAAGCCCCGCCAAAAGGCATTGTTGCTGAA  AAAGGTGATTCACGTCCGGCTGGTTTTACGGCCGGTGGTAACTCCCGTGAGGCCGTTATT  CGTTTCCCGAAAGAGACCGGACAGAAGCCGGTCTATGTGTCGGTGACAGATGTTCTTACC  CCGGCACAGGTAAAACAGCGTCAGGAGGAAGAAAAGCGTCGCCAGCAGGCATGGGACGCC  GCTCATCCGGAAGAGGGGCTGAAAAGAGAATATGATAAAGCGAAAGCTGAGCTGGATGCC  GAAGATAAAAATATTACGACTTTAAACGGCAGGATTACATCGACAGAAAAAGCGATTCCT  GGCGCCAGGGCTGCTGTCCAGGAAGCTGATAAAAAGGTGAAAGAGGCGGAGGCGAATAAG  GATGATTTTGTGACTTATAACCCTCCTCATGAATATGGCTCCGGGTGGCAGGATCAGGTT  CGCTATCTTGATAAGGATATTCAGAATCAGAATGCGAAATTAAAAGCTGCTCAGGCATCT  TTAAACGCAATGAATGATGCCTTATCCAGAGATAAGGCTGCGCTTTCCGGGGCGATGGAG  AGCCGGAAACAAAAGGAGAAAAAAGCGAAGGAGGCAGAAAATAAATTAAATGAGGAAAAG  AAAAAGCCTCGCAAGGGAACTAAAGATTATGGCCATGATTATTTTCCTGATCCCAAGACT  GAAGATATTAAGGGGTTGGGAGAGTTGAAAGAGGGAAAACCTAAAACCCCTAAACAAGGA  GGAGGTGGTAAGCGGGCCCGATGGTATGGTGATAAAAAGCGTAAAATTTATGAATGGGAT  TCCCAGCACGGTGAGCTTGAAGGATACCGCGCCAGTGATGGCGAACACCTCGGCGCATTC  GATCCAAAAACGGGTAAGCAGGTTAAAGGGCCGGATCCAAAACGAAACATTAAAAAATAT  CTTTAA | MSGGDGRGPGNSGLGHNGGQASGNVNGTSGKGGPSSGGGTDPNSGPGWGTTHTPNGDIHNYNPGEFGNGG  SKPGGNGGNSGNHSGSSGGGQSSATAMAFGLPALATPGAEGPALSFSGDALSSAVADVLAALKGPFKFGL  WGIAIYGVLPSEIAKDDPKMMSKIMTSLPADTVTETPASTLPLDQATVRVRQRVVDVVKDERQHIAVVAG  RPMSVPVVDAKPTKRPGVFSVSIPGLPALQVSVPKGVPAAKAPPKGIVAEKGDSRPAGFTAGGNSREAVI  RFPKETGQKPVYVSVTDVLTPAQVKQRQEEEKRRQQAWDAAHPEEGLKREYDKAKAELDAEDKNITTLNG  RITSTEKAIPGARAAVQEADKKVKEAEANKDDFVTYNPPHEYGSGWQDQVRYLDKDIQNQNAKLKAAQAS  LNAMNDALSRDKAALSGAMESRKQKEKKAKEAENKLNEEKKKPRKGTKDYGHDYFPDPKTEDIKGLGELK  EGKPKTPKQGGGGKRARWYGDKKRKIYEWDSQHGELEGYRASDGEHLGAFDPKTGKQVKGPDPKRNIKKY  L | ATGGGGCTTAAATTACATATTCATTGGTTTGATAAGAAAACCGAAGAGTTTAAAGGCGGT  GAATACTCAAAAGACTTCGGTGATGATGGTTCTGTCATTGAAAGTCTGGGGATGCCTTTA  AAGGATAATATTAATAATGGTTGGTTTGATGTTGAAAAACCATGGGTTTCGATATTACAG  CCACACTTTAAAAATGTAATCGATATTAGTAAATTTGATTACTTTGTATCCTTTGTTTAC  CGGGATGGTAACTGGTAA | MGLKLHIHWFDKKTEEFKGGEYSKDFGDDGSVIESLGMPLKDNINNGWFDVEKPWVSILQPHFKNVIDIS  KFDYFVSFVYRDGNW | ATGAAAAAGGCAAAGGCTATTTTTCTTTTTATATTGATAGTTTCAGGCTTTCTTCTGGTC  GCATGTCAGGCAAACTATATCCGGGATGTTCAGGGTGGAACGGTGGCACCATCGTCCTCC  TCTGAACTGACGGGGATCGCGGTTCAGTAG | MKKAKAIFLFILIVSGFLLVACQANYIRDVQGGTVAPSSSSELTGIAVQ | ATGAGTGGTGGAGACGGTCGAGGTCCGGGTAATTCAGGTCTGGGACATAATGGTGGTCAG  GCCAGTGGGAATGTGAACGGTACGTCTGGTAAAGGTGGCCCTTCATCAGGTGGTGGTACG  GATCCAAACAGCGGGCCGGGCTGGGGTACGACGCATACGCCGAATGGAGATATTCATAAC  TATAATCCGGGGGAGTTTGGTAACGGCGGGAGTAAACCCGGTGGTAATGGTGGTAACAGC  GGCAATCATAGCGGTAGCTCTGGTGGTGGACAGTCTTCGGCCACCGCGATGGCCTTCGGT  CTGCCTGCCCTGGCTACACCGGGGGCTGAAGGACCGGCTTTATCCTTTTCCGGCGATGCG  CTGTCGTCAGCCGTTGCCGATGTGCTGGCTGCCCTGAAAGGTCCGTTTAAGTTTGGTCTG  TGGGGGATTGCGATCTATGGTGTACTGCCTTCTGAGATTGCAAAAGATGATCCGAAAATG  ATGTCAAAAATTATGACGTCATTACCGGCCGATACGGTGACGGAGACTCCGGCAAGTACT  TTACCGCTGGACCAAGCGACGGTTCGTGTCAGACAACGGGTTGTGGATGTGGTGAAGGAT  GAACGGCAGCATATTGCGGTTGTCGCGGGTCGGCCAATGAGTGTTCCTGTGGTGGATGCG  AAACCGACAAAACGTCCGGGGGTATTCAGTGTGTCTATTCCGGGTCTCCCGGCTCTGCAG  GTGAGCGTACCGAAAGGTGTTCCGGCAGCGAAAGCCCCGCCAAAAGGCATTGTTGCTGAA  AAAGGTGATTCACGTCCGGCTGGTTTTACGGCCGGTGGTAACTCCCGTGAGGCCGTTATT  CGTTTCCCGAAAGAGACCGGACAGAAGCCGGTCTATGTGTCGGTGACAGATGTTCTTACC  CCGGCACAGGTAAAACAGCGTCAGGAGGAAGAAAAGCGTCGCCAGCAGGCATGGGACGCC  GCTCATCCGGAAGAGGGGCTGAAAAGAGAATATGATAAAGCGAAAGCTGAGCTGGATGCC  GAAGATAAAAATATTACGACTTTAAACGGCAGGATTACATCGACAGAAAAAGCGATTCCT  GGCGCCAGGGCTGCTGTCCAGGAAGCTGATAAAAAGGTGAAAGAGGCGGAGGCGAATAAG  GATGATTTTGTGACTTATAACCCTCCTCATGAATATGGCTCCGGGTGGCAGGATCAGGTT  CGCTATCTTGATAAGGATATTCAGAATCAGAATGCGAAATTAAAAGCTGCTCAGGCATCT  TTAAACGCAATGAATGATGCCTTATCCAGAGATAAGGCTGCGCTTTCCGGGGCGATGGAG  AGCCGGAAACAAAAGGAGAAAAAAGCGAAGGAGGCAGAAAATAAATTAAATGAGGAAAAG  AAAAAGCCTCGCAAGGGAACTAAAGATTATGGCCATGATTATTTTCCTGATCCCAAGACT  GAAGATATTAAGGGGTTGGGAGAGTTGAAAGAGGGAAAACCTAAAACCCCTAAACAAGGA  GGAGGTGGTAAGCGGGCCCGATGGTATGGTGATAAAAAGCGTAAAATTTATGAATGGGAT  TCCCAGCACGGTGAGCTTGAAGGATACCGCGCCAGTGATGGCGAACACCTCGGCGCATTC  GATCCAAAAACGGGTAAGCAGGTTAAAGGGCCGGATCCAAAACGAAACATTAAAAAATAT  CTTTAAGAGGTAAATATGGGGCTTAAATTACATATTCATTGGTTTGATAAGAAAACCGAA  GAGTTTAAAGGCGGTGAATACTCAAAAGACTTCGGTGATGATGGTTCTGTCATTGAAAGT  CTGGGGATGCCTTTAAAGGATAATATTAATAATGGTTGGTTTGATGTTGAAAAACCATGG  GTTTCGATATTACAGCCACACTTTAAAAATGTAATCGATATTAGTAAATTTGATTACTTT  GTATCCTTTGTTTACCGGGATGGTAACTGGTAAGCAAAAGTTATCAAGGATGAGTTGAAA  TACAGGCTTCCGGTGTTCACGGATGAATGCCGAAGCCTCTGAATACAAGGAAGGTATATG  AAAAAGGCAAAGGCTATTTTTCTTTTTATATTGATAGTTTCAGGCTTTCTTCTGGTCGCA  TGTCAGGCAAACTATATCCGGGATGTTCAGGGTGGAACGGTGGCACCATCGTCCTCCTCT  GAACTGACGGGGATCGCGGTTCAGTAG | ATGAGTGGTGGAGACGGTCGAGGTCCGGGTAATTCAGGTCTGGGACATAATGGTGGTCAG  GCCAGTGGGAATGTGAACGGTACGTCTGGTAAAGGTGGCCCTTCATCAGGTGGTGGTACG  GATCCAAACAGCGGGCCGGGCTGGGGTACGACGCATACGCCGAATGGAGATATTCATAAC  TATAATCCGGGGGAGTTTGGTAACGGCGGGAGTAAACCCGGTGGTAATGGTGGTAACAGC  GGCAATCATAGCGGTAGCTCTGGTGGTGGACAGTCTTCGGCCACCGCGATGGCCTTCGGT  CTGCCTGCCCTGGCTACACCGGGGGCTGAAGGACCGGCTTTATCCTTTTCCGGCGATGCG  CTGTCGTCAGCCGTTGCCGATGTGCTGGCTGCCCTGAAAGGTCCGTTTAAGTTTGGTCTG  TGGGGGATTGCGATCTATGGTGTACTGCCTTCTGAGATTGCAAAAGATGATCCGAAAATG  ATGTCAAAAATTATGACGTCATTACCGGCCGATACGGTGACGGAGACTCCGGCAAGTACT  TTACCGCTGGACCAAGCGACGGTTCGTGTCAGACAACGGGTTGTGGATGTGGTGAAGGAT  GAACGGCAGCATATTGCGGTTGTCGCGGGTCGGCCAATGAGTGTTCCTGTGGTGGATGCG  AAACCGACAAAACGTCCGGGGGTATTCAGTGTGTCTATTCCGGGTCTCCCGGCTCTGCAG  GTGAGCGTACCGAAAGGTGTTCCGGCAGCGAAAGCCCCGCCAAAAGGCATTGTTGCTGAA  AAAGGTGATTCACGTCCGGCTGGTTTTACGGCCGGTGGTAACTCCCGTGAGGCCGTTATT  CGTTTCCCGAAAGAGACCGGACAGAAGCCGGTCTATGTGTCGGTGACAGATGTTCTTACC  CCGGCACAGGTAAAACAGCGTCAGGAGGAAGAAAAGCGTCGCCAGCAGGCATGGGACGCC  GCTCATCCGGAAGAGGGGCTGAAAAGAGAATATGATAAAGCGAAAGCTGAGCTGGATGCC  GAAGATAAAAATATTACGACTTTAAACGGCAGGATTACATCGACAGAAAAAGCGATTCCT  GGCGCCAGGGCTGCTGTCCAGGAAGCTGATAAAAAGGTGAAAGAGGCGGAGGCGAATAAG  GATGATTTTGTGACTTATAACCCTCCTCATGAATATGGCTCCGGGTGGCAGGATCAGGTT  CGCTATCTTGATAAGGATATTCAGAATCAGAATGCGAAATTAAAAGCTGCTCAGGCATCT  TTAAACGCAATGAATGATGCCTTATCCAGAGATAAGGCTGCGCTTTCCGGGGCGATGGAG  AGCCGGAAACAAAAGGAGAAAAAAGCGAAGGAGGCAGAAAATAAATTAAATGAGGAAAAG  AAAAAGCCTCGCAAGGGAACTAAAGATTATGGCCATGATTATTTTCCTGATCCCAAGACT  GAAGATATTAAGGGGTTGGGAGAGTTGAAAGAGGGAAAACCTAAAACCCCTAAACAAGGA  GGAGGTGGTAAGCGGGCCCGATGGTATGGTGATAAAAAGCGTAAAATTTATGAATGGGAT  TCCCAGCACGGTGAGCTTGAAGGATACCGCGCCAGTGATGGCGAACACCTCGGCGCATTC  GATCCAAAAACGGGTAAGCAGGTTAAAGGGCCGGATCCAAAACGAAACATTAAAAAATAT  CTTTAA  ATGGGGCTTAAATTACATATTCATTGGTTTGATAAGAAAACCGAAGAGTTTAAAGGCGGT  GAATACTCAAAAGACTTCGGTGATGATGGTTCTGTCATTGAAAGTCTGGGGATGCCTTTA  AAGGATAATATTAATAATGGTTGGTTTGATGTTGAAAAACCATGGGTTTCGATATTACAG  CCACACTTTAAAAATGTAATCGATATTAGTAAATTTGATTACTTTGTATCCTTTGTTTAC  CGGGATGGTAACTGGTAA | MSGGDGRGPGNSGLGHNGGQASGNVNGTSGKGGPSSGGGTDPNSGPGWGTTHTPNGDIHNYNPGEFGNGG  SKPGGNGGNSGNHSGSSGGGQSSATAMAFGLPALATPGAEGPALSFSGDALSSAVADVLAALKGPFKFGL  WGIAIYGVLPSEIAKDDPKMMSKIMTSLPADTVTETPASTLPLDQATVRVRQRVVDVVKDERQHIAVVAG  RPMSVPVVDAKPTKRPGVFSVSIPGLPALQVSVPKGVPAAKAPPKGIVAEKGDSRPAGFTAGGNSREAVI  RFPKETGQKPVYVSVTDVLTPAQVKQRQEEEKRRQQAWDAAHPEEGLKREYDKAKAELDAEDKNITTLNG  RITSTEKAIPGARAAVQEADKKVKEAEANKDDFVTYNPPHEYGSGWQDQVRYLDKDIQNQNAKLKAAQAS  LNAMNDALSRDKAALSGAMESRKQKEKKAKEAENKLNEEKKKPRKGTKDYGHDYFPDPKTEDIKGLGELK  EGKPKTPKQGGGGKRARWYGDKKRKIYEWDSQHGELEGYRASDGEHLGAFDPKTGKQVKGPDPKRNIKKY  L  MGLKLHIHWFDKKTEEFKGGEYSKDFGDDGSVIESLGMPLKDNINNGWFDVEKPWVSILQPHFKNVIDIS  KFDYFVSFVYRDGNW | MSGGDGRGPGNSGLGHNGGQASGNVNGTSGKGGPSSGGGTDPNSGPGWGTTHTPNGDIHNYNPGEFGNGG  SKPGGNGGNSGNHSGSSGGGQSSATAMAFGLPALATPGAEGPALSFSGDALSSAVADVLAALKGPFKFGL  WGIAIYGVLPSEIAKDDPKMMSKIMTSLPADTVTETPASTLPLDQATVRVRQRVVDVVKDERQHIAVVAG  RPMSVPVVDAKPTKRPGVFSVSIPGLPALQVSVPKGVPAAKAPPKGIVAEKGDSRPAGFTAGGNSREAVI  RFPKETGQKPVYVSVTDVLTPAQVKQRQEEEKRRQQAWDAAHPEEGLKREYDKAKAELDAEDKNITTLNG  RITSTEKAIPGARAAVQEADKKVKEAEANKDDFVTYNPPHEYGSGWQDQVRYLDKDIQNQNAKLKAAQAS  LNAMNDALSRDKAALSGAMESRKQKEKKAKEAENKLNEEKKKPRKGTKDYGHDYFPDPKTEDIKGLGELK  EGKPKTPKQGGGGKRARWYGDKKRKIYEWDSQHGELEGYRASDGEHLGAFDPKTGKQVKGPDPKRNIKKY  L  MGLKLHIHWFDKKTEEFKGGEYSKDFGDDGSVIESLGMPLKDNINNGWFDVEKPWVSILQPHFKNVIDIS  KFDYFVSFVYRDGNW  MKKAKAIFLFILIVSGFLLVACQANYIRDVQGGTVAPSSSSELTGIAVQ |
